# Temporal Interference Stimulation Enhances Neural Regeneration

**DOI:** 10.1101/2025.08.18.670811

**Authors:** Sofia Peressotti, Maria Garcia Garrido, Patrycja Dzialecka, Rachel Man Hoi Law, Roberto Portillo-Lara, Bethany Geary, Elena Faillace, Marcelina Wojewska, Maria Otero-Jimenez, Martina Genta, Luqiao Tan, Karen Duff, Javier Alegre-Abarrategui, Rylie Green, Nir Grossmann

**Affiliations:** Bioengineering Department, Imperial College London, South Kensington, London, SW7 2AZ, UK; Department of Brain Sciences, Imperial College London, Hammersmith Hospital, London, W12 0NN, UK; UK Dementia Research Institute, London NW1 3BT, UK; Medical Research Council Protein Phosphorylation and Ubiquitylation Unit, University of Dundee, DD1 4HN, UK; Institute of Neurology, University College London, London WC1E 6BT, UK; Psychiatry and Fundamental Neuroscience Department, University of Geneva, Geneva 1202, CH

**Author notes:** These authors contributed equally (Sofia Peressotti, Maria Garcia Garrido). Shared last authorship (Rylie Green, Nir Grossmann).

## Abstract

Neural regeneration therapies aim to treat neurodegeneration by promoting the proliferation and maturation of exogenous or endogenous neural progenitor cells (NPCs). However, their efficacy has been limited. Deep brain stimulation (DBS) via implanted electrodes has been shown to promote neurogenesis. However, its invasiveness precludes deployment in research and widespread clinical use. Temporal interference (TI) has emerged as a strategy for non-invasive, high-precision DBS using multiple kHz-range electric fields, with a frequency difference within the range of neural activity. Here, we validate the potential of TI stimulation for neural regeneration augmentation. We demonstrate that TI stimulation with a theta-band frequency difference enhances the maturation of embryonic neural progenitor cells *in vitro*. We then demonstrate that theta-band TI stimulation targeting the hippocampus enhances endogenous hippocampal neurogenesis in an *in vivo* mouse model of Alzheimer’s disease. By uncovering frequency-specific control of stem cell fate, we propose a clinically relevant regeneration strategy which avoids pharmacological or genetic manipulation. Our results demonstrate focal, non-invasive augmentation of deep-brain neural regeneration.

**GRAPHICAL ABSTRACT:** 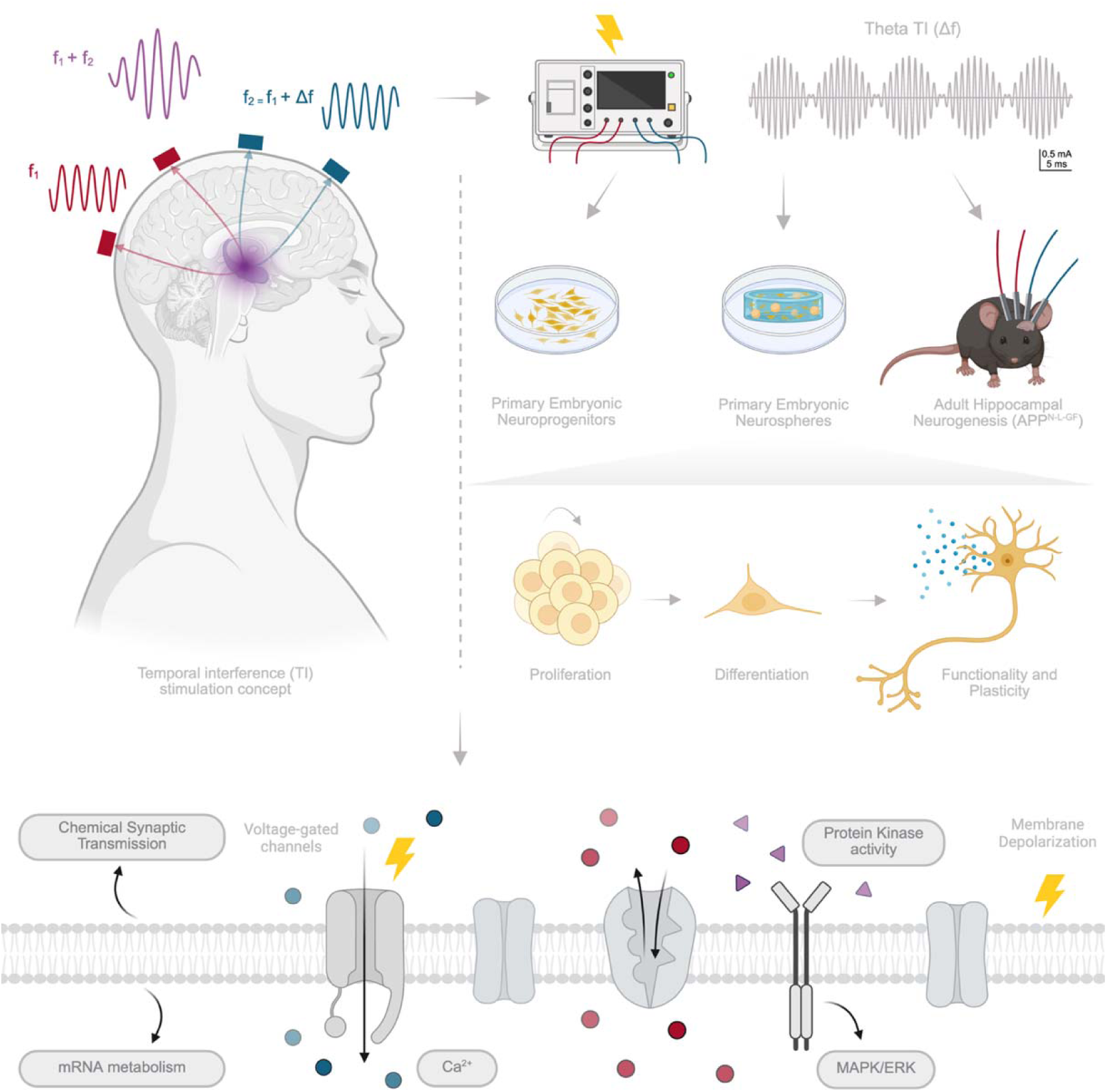

## 1. INTRODUCTION

Neurodegenerative diseases are characterized by a progressive loss of neural structure and function in the central nervous system^12,3^. Alzheimer’s disease (AD) and Parkinson’s disease (PD) are the most prevalent neurodegenerative diseases, approximately affecting 60 million patients worldwide^4,5^ and representing the leading cause of disability in elderly populations^2,3^. The complex and heterogeneous pathophysiology of neurodegenerative diseases, combined with the limited capacity of neural cells to regenerate spontaneously^6–9^, presents a significant challenge for the discovery and development of effective therapies^10^.

Neural regeneration therapies aspire to ameliorate neurodegeneration by directly counteracting neural loss and its associated cognitive impairment^6,7^. A promising approach is based on cell replacement therapies, where neural progenitor cells (NPCs), such as those derived from embryonic stem cells^11^, are transplanted into the affected brain regions to replace diseased cells and restore network functions^12^. An alternative therapeutic approach is based on boosting endogenous neural stem cells^9,13,14^. NPCs in the adult brain have the innate ability to differentiate and integrate into existing functional circuits across the human lifespan^9,13,15–20^. These cells are found in specific deep brain regions, such as the dentate gyrus (DG) of the hippocampus. Notably, the hippocampal formation is one of the most vulnerable regions in early-stage AD^21^. Adult hippocampal neurogenesis (AHN) is impaired in neurodegenerative diseases, such as AD and other neurological conditions^15–17,22,23^. The development of neural regeneration therapies leveraging exogenous or endogenous NPCs for neurodegenerative diseases is still a major challenge, limited by the trade-off between risks and effectiveness^6,24–26^.

Accumulating evidence has demonstrated that electrical stimulation of NPCs can promote their proliferation, differentiation, and migration through complex mechanisms, including calcium influx, mitochondrial activity^27,28^, and membrane depolarization^29–31^. Invasive electrical stimulation of deep brain regions such as the thalamus^32–35^, fornix^36,37^ or entorhinal cortex^38,39^ has been shown to promote AHN in animal models. However, the invasiveness of conventional DBS techniques bears the risk of serious surgical complications, limiting its therapeutic potential^40^. Thus, non-invasive electrical stimulation of the deep-brain neurogenic niches has been recently shown using transcranial direct current electrical stimulation (tDCS)^41,42^, transcranial alternating current stimulation (tACS)^43^, and transcranial magnetic stimulation (TMS)^44^. However, the use of these modalities induces stronger stimulation of the overlying cortical areas, pushing the limits of safety guidelines^45^ and making the spatial precision of DBS a key feature to target cell transplants and neurogenic niches.

We recently developed a strategy for non-invasive and focal electrical deep brain stimulation via temporal interference (TI) of kHz electric fields^46^. In TI stimulation, two or more electric fields are applied to the brain at different kHz frequencies, which are higher than the range of neural activity. These electric fields are superimposed at the target region to create a combined electric field, with an amplitude that changes periodically at the slow frequency difference between waveforms. This frequency difference is slower, and it falls within the range of neural activity. The strength of TI neural stimulation depends not only on the absolute amplitude, but also on the relative amplitude and orientation of the applied electric fields, allowing for pinpointing the stimulation target in three dimensions. We have demonstrated that non-invasive transcranial TI electrical stimulation can selectively modulate hippocampal activity in both rodents^46^ and humans^47^.

Herein, we investigate whether TI electrical stimulation can be used as a tool for neural regeneration therapy. First, we show that electrical TI stimulation of embryonic NPCs promotes their proliferation and differentiation in both 2D and 3D *in vitro* cultures, using immunofluorescence, calcium imaging, and mRNA transcriptomics analyses. Then, we demonstrate that electrical TI stimulation of the hippocampus enhances endogenous AHN in a mouse model of AD *in vivo*, using immunohistochemistry and proteomics analysis. This augmentation of both embryonic NPCs and endogenous AHN by TI stimulation occurs specifically at the theta frequency band. Given the clinical implementation of TI stimulation for precise and non-invasive hippocampal neuromodulation^47^, the demonstration of TI’s neurogenic effect opens an exciting therapeutic opportunity for the large patient population affected by AD and other neurodegenerative diseases.

## 2. RESULTS

### 2.1 Theta-band TI stimulation augments differentiation in 2D *in vitro* cultures of embryonic NPCs

First, we tested whether TI stimulation can modulate the differentiation of embryonic NPCs, as they are often employed in cell replacement therapies for neurodegenerative disease. We applied TI stimulation to primary cultures of embryonic NPCs from rat embryos and measured changes in neural differentiation relative to a sham stimulation at DIV10. The NPCs were harvested from the ventral-mesencephalon (VM) of day-14 rat embryos (E14) and grown in a monolayer culture for 7 days *in vitro* (DIV) prior to stimulation (culture characterization in **Figure S1**). Stimulation-induced changes in metabolic activity and cell differentiation relative to sham were measured at DIV10 (**Figure 1A**). The TI electrical stimuli were selected to recapitulate endogenous waveforms in the developing brain^46^. The currents were applied using a 1 kHz carrier frequency (f_1_), and a frequency difference between waveforms (f_2_= f1+Δf) at the theta-band (Δf=10 Hz; f_1_=1000 Hz + f_2_=1010 Hz), low-gamma band (Δf=40 Hz; f_1_=1000 Hz + f_2_=1040 Hz), or high-gamma band (Δf=100 Hz; f_1_=1000 Hz + f_2_=1100 Hz). A carrier frequency condition (sinusoidal waveform at f_1_=1000 Hz) was used as a control for the high frequency effect. The sham condition was a tissue culture well in contact with the electrodes, where no stimulation was delivered. The two sinusoidal frequencies (f_1,_ f_2_) were combined using MATLAB to minimize variability in electric field exposure across the cell monolayer, and a 1 mA current was delivered via two platinum electrodes (current density: 1.91 mA/mm^2^). Each stimulation condition was applied for three days and 6 h of stimulation per day, consisting of 2-second on and 2-second off periods, mimicking developmental neural patterns^48,49^(**Figure 1A**). At the end of the stimulation period (24 h after the last stimulation session), we measured cell density, metabolic activity via an Alamar Blue assay, and the state of cell pluripotency and differentiation using immunofluorescence (IF) staining for Sox2^50,51^, β-tubulin III^52^, and GFAP^53–55^.

**Figure 1:**
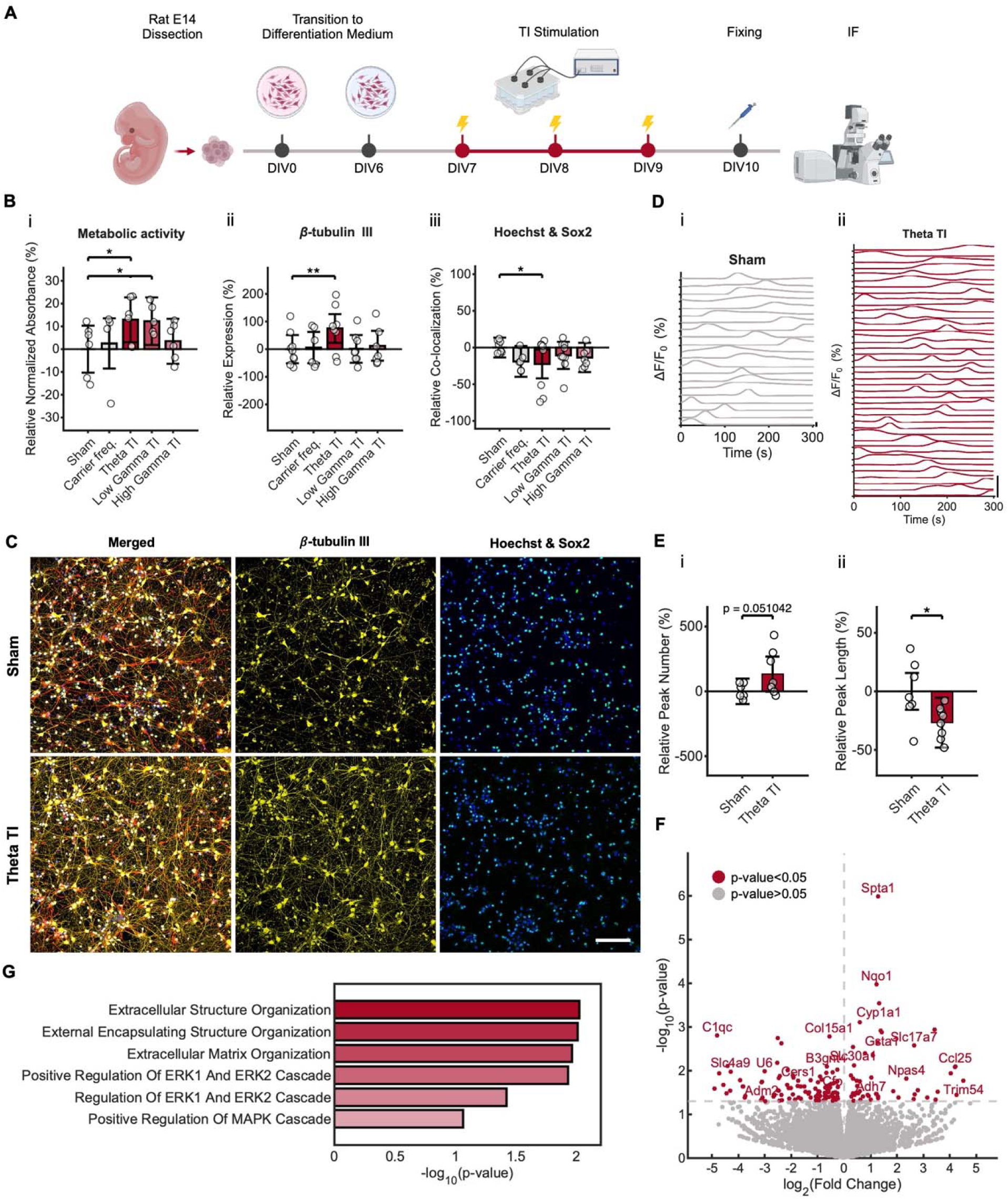
Theta-band TI stimulation augments differentiation in 2D in vitro cultures of embryonic NPCs. **(A)** Schematic of the experimental timeline. After dissociation of the VM cells at E14, the cells were transitioned from the proliferation to the differentiation medium (DIV0 to DIV6). Stimulation was delivered for 3 days (DIV7-DIV9), 6 h/day, 2s on-2s off. After the stimulation period (DIV10), the metabolic activity of NPCs was measured using an Alamar Blue assay and fixed for IF at DIV10. **(B)** Stimulation-induced change in NPC differentiation markers. (i) Alamar Blue assay (NPC metabolic activity), treatment effect F(4,25)=2.86, p=0.044, repeat effect χ^2^=5.41, p=0.020, linear mixed-effects model (LMM); Theta TI: p=0.013, Low-gamma TI: p=0.0304, High-gamma TI: p=0.9421, carrier: p=2.5716, Bonferroni-Holm (BH) correction. (ii) β-tubulin III expression (NPC differentiation), treatment effect F(4,36)=2.95, p=0.032, repeat effect χ^2^=6.77, p=0.009, LMM; Theta TI: p=0.0059, Low-gamma TI: p=3.8296, High-gamma TI: p=0.8868, Carrier freq.: p=1.6976, BH correction. (iii) Spatial co-localization quantified as Pearson’s correlation between Sox2 and β-tubulin III, treatment effect F(4,39)=1.63, p=0.185, repeat effect χ^2^=0, p=1, LMM; Theta TI: p=0.024, Low-gamma TI: p=1.027, High-gamma TI: p=0.35, Carrier freq.: p=0.1, BH correction. Values are expressed as a percentage relative to the mean of the sham. Data points show the average measurements extracted from each tissue culture well (n=6-9 wells per condition from 3 rats). Bar plots show the LMM predicted means ± 95% confidence interval (CI). See **Table S1** for details on the statistical analysis. **(C)** Representative fluorescent micrographs after theta-band stimulation and sham. Hoechst for nuclei (blue), Sox2 for pluripotent cells (green), β-tubulin III for neurons (yellow), GFAP for astrocytes (red). Scale bar: 100 µm. **(D)** Stimulation-induced change in NPC calcium activity. Representative ΔF/F_0_ calcium traces after (i) sham and (ii) theta-band stimulation. Scale bar: 50%. **(E)** Comparison of calcium peaks’ characteristics between conditions. (i) Calcium peaks number, treatment effect F(1,15)=4.61, p=0.048, repeat effect χ^2^=0, p=1, Theta TI: p=0.051. (ii) Calcium peak duration, treatment effect F(1,15)=7.30, p=0.016, repeat effect χ^2^=0, p=1; Theta TI: p=0.018, LMM. Values are expressed as a percentage relative to the mean of the sham. Data points show the average measurements extracted from each tissue culture well (n=6-9 recordings per condition from 3 rats). Bar plots show the LMM predicted means ± 95% confidence interval (CI). See **Table S2** for details on the statistical analysis. **(F)** Volcano plot showing genes differentially expressed after theta-band stimulation relative to sham (total 153 genes labelled in red; cutoff: p<0.05). **(G)** GO enrichment of biological process of the overexpressed genes (cutoff: p<0.05, log_2_(Fold Change)>0). The gene sets contain over 3 significantly overexpressed genes with an adjusted p<0.2. ^∗^p<0.05, ^∗∗^p<0.01, with p displaying the p-value adjusted for multiple comparisons with BH correction.

The stimulation conditions with frequency difference at the theta and low-gamma bands showed a higher metabolic activity (**Figure 1Bi**). Interestingly, the stimulation condition with a frequency difference Δf at the theta band, but not other bands, displayed a higher expression of β-tubulin III (**Figure 1Bii**). The β-tubulin III increase remained significant when normalizing the bulk expression by the total number of cells (**Figure S2**). The stimulation conditions did not affect the overall cell density or the Sox2+ cell density relative to sham (**Figure S2**). However, the theta-band condition also revealed a lower co-localization between Sox2 and the nuclei marker Hoechst (**Figure 1Biii**), which is associated with a more differentiated (less pluripotent) state of NPCs^50,51^. See **Figure 1C** for representative fluorescent micrographs. There was also no change in GFAP expression, a marker for glial differentiation (**Figure S2**).

To assess whether the stimulation-induced differentiation involved changes in cell activity, we repeated the experiment (1 kHz carrier frequency with theta-band difference and sham), and measured live calcium transients in the culture at 10 DIV, using the Fluo-4 AM fluorophore. The differentiation induced by the theta band stimulation was associated with shorter and more frequent calcium peaks (**Figure 1D-E**), which is consistent with earlier studies in NPC *in vitro* cultures^56–60^.

To investigate molecular pathways underlying stimulation-induced differentiation, we repeated the experiment (1 kHz carrier with a theta-band Δf and sham) and performed bulk mRNA sequencing 24 h after the stimulation period. We identified 153 genes that were differentially expressed in the theta-band stimulation compared to sham (Figure 1F, p<0.05, DESeq2 normalisation). The enrichment analysis revealed that the overexpressed gene sets were linked to the ERK/MAPK signalling pathway (**Figure 1G**, cut-off: p<0.05, log_2_(Fold Change)>0, Gene ontology (GO) enrichment of biological process), consistent with earlier studies examining the effect of electrical stimulation on neural differentiation^29,61–63^. We also found upregulated gene sets related to extracellular matrix reorganisation.

Overall, the 2D *in vitro* embryonic NPCs results demonstrated that TI stimulation at the theta-band frequency difference Δf boosts functional differentiation in 2D cultures of embryonic NPCs via molecular signalling pathways including the ERK/MAPK cascade.

### 2.2 Theta-band TI stimulation modulates differentiation in a 3D *in vitro* model of embryonic NPCs

Earlier studies have demonstrated that 3D *in vitro* models of NPCs display more physiologically accurate developmental dynamics and microenvironments compared to 2D cultures, resulting in longer differentiation timelines^64–66^. Thus, we repeated the stimulation paradigm (1 kHz carrier frequency with theta-band frequency difference Δf and sham) in a 3D *in vitro* culture of embryonic NPCs, to increase the physiological relevance of the therapeutic approach. Primary embryonic NPCs were cultured in suspension for four days to allow the formation of neurospheres. At DIV4, the neurospheres were encapsulated in a bioactive hydrogel functionalised with a laminin-derived epitope to support neural growth and differentiation. Identical culture conditions and stimulation protocols as in the original 2D NPC study were used. The 3-day stimulation sessions began at day of encapsulation (DOE) 7, and the cells were fixed at DOE 10 for IF characterization as before (**Figure 2A**).

**Figure 2:**
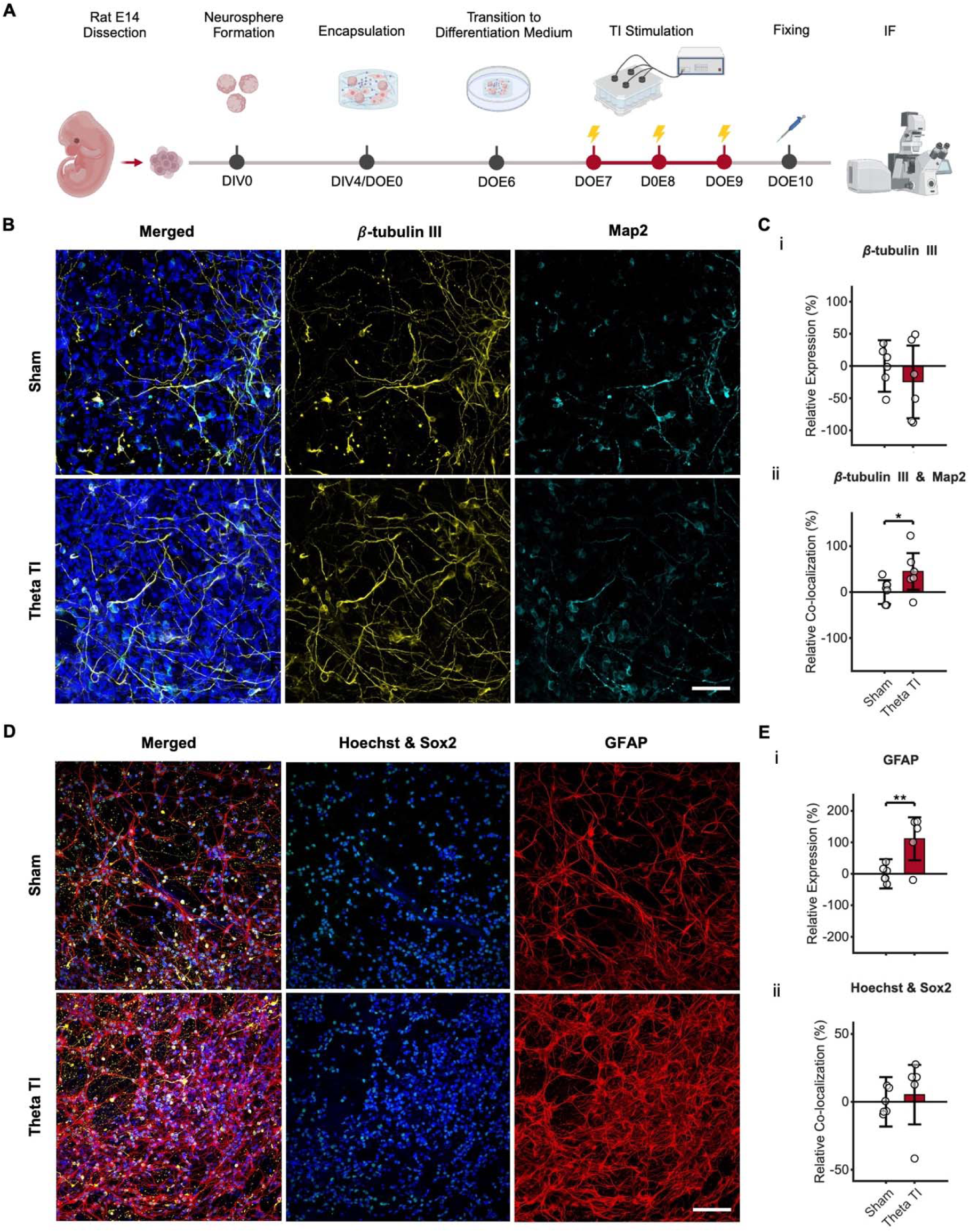
Theta-band TI stimulation modulates differentiation in a 3D in vitro model of embryonic NPCs. **(A)** Schematic of the experimental timeline. After dissociation of the VM cells at E14, the cells were grown in suspension to generate neurospheres, which were encapsulated at DIV4 (DOE0). The cells were transitioned from the proliferation to the differentiation medium (DOE0 to DOE6). Stimulation was delivered for 3 days (DOE7-DOE9), 6 h/day, 2s on-2s off. After the stimulation period (DIV10), NPCs were exposed to Alamar Blue to measure metabolic activity and fixed for IF. **(B)** Representative fluorescent micrographs after theta-band stimulation and sham. Hoechst for nuclei (blue), β-tubulin III for neurons (yellow), Map2 for mature neurons (cyan). Scale bar: 100 µm. **(C)** Comparison of IF markers between conditions. (i) β-tubulin III expression, treatment effect F(1,12)=0.92, p=0.36, repeat effect χ^2^=0, p=1, LMM, n=5-6 wells from 4 rats. (ii) Spatial co-localization quantified as Pearson’s correlation between β-tubulin III and Map2, treatment effect F(1,9)=6.22, p=0.033, repeat effect χ^2^=0.007, p=0.93, LMM, n=6 wells from 4 rats. Values are expressed as a percentage relative to the mean of the sham. Data points show the average measurements extracted from each tissue culture well (n=5-6 wells per condition from 4 rats). Bar plots show the LMM predicted means ± 95% confidence interval (CI). See **Table S3** for details on the statistical analysis. **(D)** Representative fluorescent micrographs after theta-band stimulation and sham. Hoechst for nuclei (blue), Sox2 for progenitor cells (green), β-tubulin III for neurons (yellow), GFAP for astrocytes (red). Scale bar: 100 µm. **(E)** Comparison of neural IF markers between conditions. (i) GFAP expression, treatment effect F(1,11)=13.62, p=0.004, repeat effect χ^2^=0, p=1, LMM, n=5-6 wells from 4 rats. (ii) Spatial co-localization quantified as Pearson’s correlation between Hoechst (nuclei) and Sox2, treatment effect F(1,8)=0.29, p=0.60, repeat effect χ^2^=0.435, p=0.509, LMM, n=5-6 wells from 4 rats. Values are expressed as a percentage relative to the mean of sham. Data points show the average measurements extracted from each tissue culture well (n=5-6 wells per condition from 4 rats). Bar plots show the LMM predicted means ± 95% confidence interval (CI). See **Table S3** for details on the statistical analysis. ^∗^p<0.05, ^∗∗^p<0.01, with p displaying the p-value adjusted for multiple comparisons with BH correction.

At DOE10, the 3D construct formed an interconnected tissue structure, resembling the morphology of embryonic tissue *in vivo* (**Figure S1**). We qualitatively observed a lower abundance of β-tubulin III+ cells, with a less extended morphology in cells grown in 3D (**Figure 2B**), consistent with 3D *in vitro* timelines and morphological features^64–66^. In this case, we did not find differences in cell density (**Figure S2**), Sox2+ cell density (**Figure S2**), metabolic activity (**Figure S2**), and β-tubulin III expression between the conditions (**Figure 2B-Ci**). There was also no difference in the correlation between Sox2 and the nuclear marker Hoechst (**Figure 2D-Eii**). To explore effects on a later maturation stage, we stained for microtubule-associated protein 2 (Map2)^52^. Although there was no change in Map2 expression (**Figure S2**), the stimulation resulted in a higher co-localization between Map2 and β-tubulin III, (**Figure 2B-Cii**), suggesting increased maturation in NPCs which began differentiating before the stimulation onset^52^. Interestingly, a higher expression of GFAP was detected after the theta-band stimulation (**Figure 2D**-**Ei**), which may entail an increase in astrocytic differentiation in parallel to neural differentiation^64,66–70^ (**Figure 2E**). We did not observe a qualitative change in astrocyte morphology (**Figure 2D**).

Taken together, these results strengthen the evidence that theta-band TI stimulation enhances neural differentiation in 3D cultures of embryonic NPCs. However, the effect size was small at the developmental stage we studied. Further investigations are needed to pinpoint the differentiation effect accounting for differences in developmental dynamics and field exposure timelines.

### 2.3 Theta-band TI stimulation augments adult hippocampal neurogenesis in an *in vivo* mouse model of AD

After establishing that electrical TI stimulation enhances embryonic NPC neural differentiation, we investigated whether this approach can also promote the differentiation of adult NPCs and their progeny in a neurodegenerative disease model *in vivo*^13,20,96^. Adult hippocampal neurogenesis (AHN) has been shown to play an essential role in cognitive processes such as learning and memory, mood regulation, stress and anxiety responses^13,22^. A potential therapeutic approach consists in boosting adult neurogenesis to enhance its neuroprotective effect and/or restore the associated cognitive functions^9,71^. To test whether AHN can be modulated by TI stimulation with frequency differences Δf endogenous to the developing brain^72^, we applied TI stimulation to the right hippocampus of a 6-7-month-old APP^N-L-GF^ mouse model^73^. Changes in hippocampal NPC proliferation and differentiation were measured in the ipsilateral DG. The TI stimulation electric currents were applied with a 2 kHz carrier frequency and a frequency difference at the delta-band (Δf=1 Hz; f_1_=2000 Hz + f_2_=2001 Hz), theta-band (Δf=8 Hz; f_1_=2000 Hz + f_2_=2008 Hz), or low-gamma band (Δf=40 Hz; f_1_=2000 Hz + f_2_=2040 Hz). The currents were applied at a current density of approximately 0.96 ± 0.29 mA/mm^2^ via two electrode pairs for eight days, 1 h of stimulation per day, while the mice were freely moving (**Figure 3A**). In the sham condition, mice underwent the minimally invasive electrode implantation procedure, and were connected to the TI stimulation setup, but no current was delivered. At the end of the stimulation period (1 h after the last stimulation session), we perfused the mice, fixed the hippocampal tissue, and measured the state of proliferation and differentiation using immunohistochemistry staining for Kiel 67 (Ki67)^74^ and doublecortin (DCX)^75^, respectively. We also explored changes in amyloid-beta aggregates using 12f4 and ct695^76^ staining, and changes in astrocyte and microglia using GFAP and Iba1 staining^77^, respectively.

**Figure 3:**
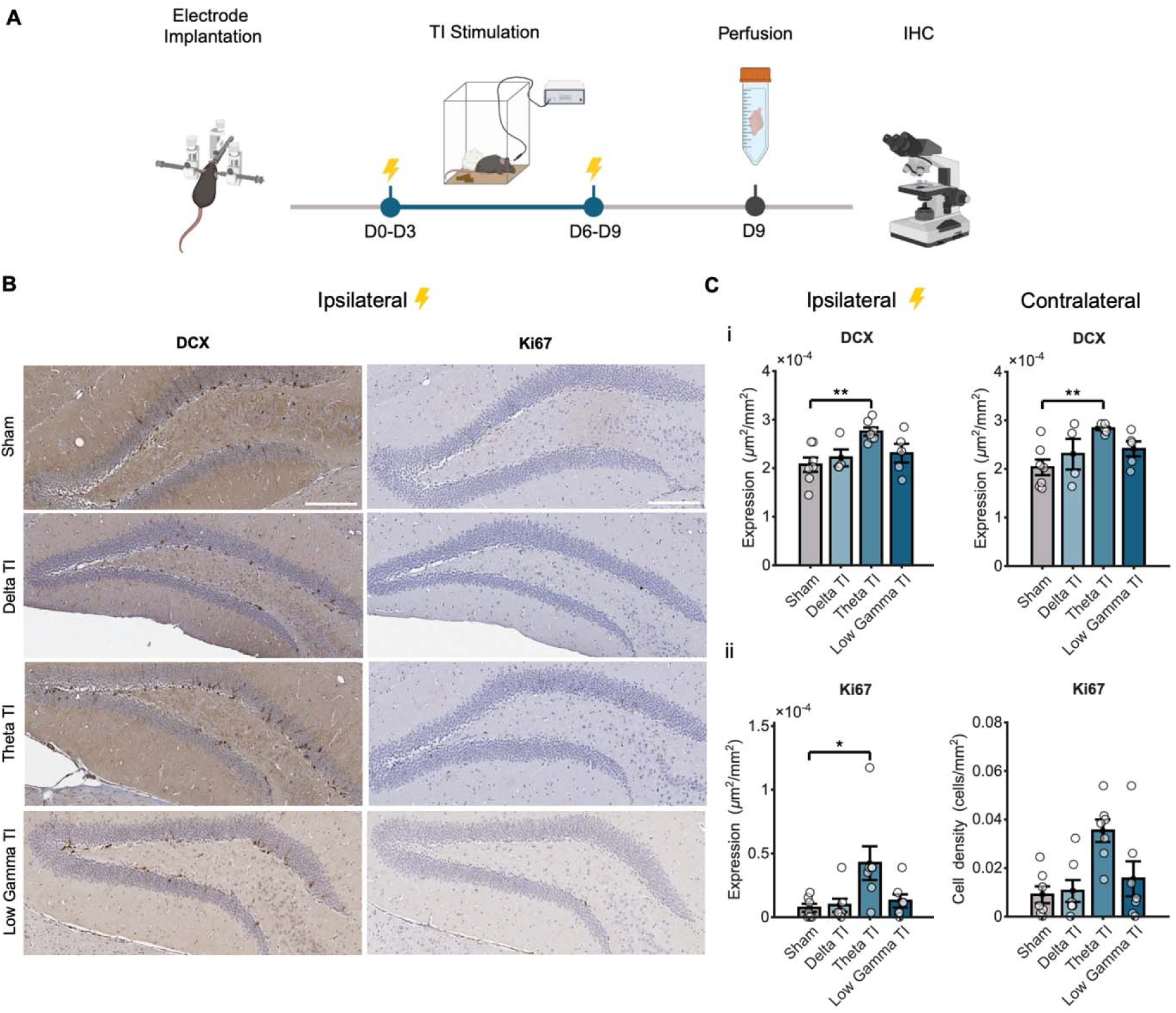
Theta-band TI stimulation augments adult hippocampal neurogenesis in an in vivo mouse model of AD. **(A)** Schematic of the experimental timeline. After recovery from surgery, stimulation was delivered to awake, free-moving APP^NL-G-F^ mice for 8 days (days 0 to 3 and 6 to 9), 1 h per day. All mice were perfused for immunohistochemistry (IHC) at day 9, 1 h after stimulation. **(B)** Representative brightfield images. Left: DCX expression (differentiation marker). Right: Ki67 expression (proliferation marker). Biomarkers were stained with DAB (brown), and cell nuclei were counterstained with haematoxylin (blue). Scale bar: 200 µm. **(C)** Comparison of IHC markers between conditions in the ipsilateral and contralateral DGs. (i) DCX expression in µm^2^ per mm^2^, DCX (ipsi): Delta TI: p=0.69, Theta TI: p=0.012, Low Gamma TI: p=0.69. DCX (contra), Delta TI: p=0.37, Theta TI: p=0.003, Low gamma TI: p=0.28; (ii) Ki67 expression in µm^2^ per mm^2^. Ki67 (ipsi): Delta TI: p=0.74, Theta TI: p=0.049, Low-gamma TI: p=0.58; Ki67 (contra): Delta TI: p=1, Theta TI: p=0.37, Low-gamma TI: p=1; sham: n=7 mice, delta: n=4, theta: n=6, gamma: n=5. Each data point represents an individual mouse. Results are expressed as mean ± standard error of the mean (SEM). Significance was calculated using an independent t-test if the data passed the normality test (Lilliefors, p>0.05), or a Wilcoxon rank-sum test otherwise, and a post-hoc BH correction; ^∗^p<0.05, ^∗∗^p<0.01. See **Table S4** for details on the statistical analysis.

Hippocampal TI stimulation with a frequency difference Δf at the theta-band, but not other bands, increased the expression of the neural differentiation marker DCX and the proliferation marker Ki67 in in the ipsilateral DG (**Figure 3B-C**). The contralateral DG was then quantified, where an increase in DCX and a non-significant increase in Ki67 expression compared to sham was detected specifically after theta TI stimulation (**Figure 3B-C**). Theta TI stimulation in the ipsilateral hippocampus was also associated with an increase in the microglia marker Iba1 (**Figure S3**). There were no changes in the astrocyte marker GFAP, nor in the amyloid beta markers 12f4 and ct695 (**Figure S3**).

To further investigate the effect of TI stimulation on adult NPC differentiation, changes in DCX expression in different regions of the granule cell layer (GCL) were quantified. The theta TI stimulation increased DCX expression in the ipsilateral and contralateral GCL combined (**Figure 4B-C**). When segmented into DG regions, the DCX increase was associated specifically to the dorsal granule cell layer (dGCL) and not the ventral granule cell layer (vGCL) (**Figure 4B-C**). The morphology of the DCX+ cells was then quantified to evaluate neural cell maturation (i.e., proliferative, intermediate, or postmitotic, **Figure 4A**)^78^. Here, the increase in DCX+ cell density was consistent across all maturation stages (**Figure 4D**), suggesting an enhancement of proliferation, differentiation and perhaps survival of the NPCs in the hippocampal dDGL niche.

**Figure 4.**
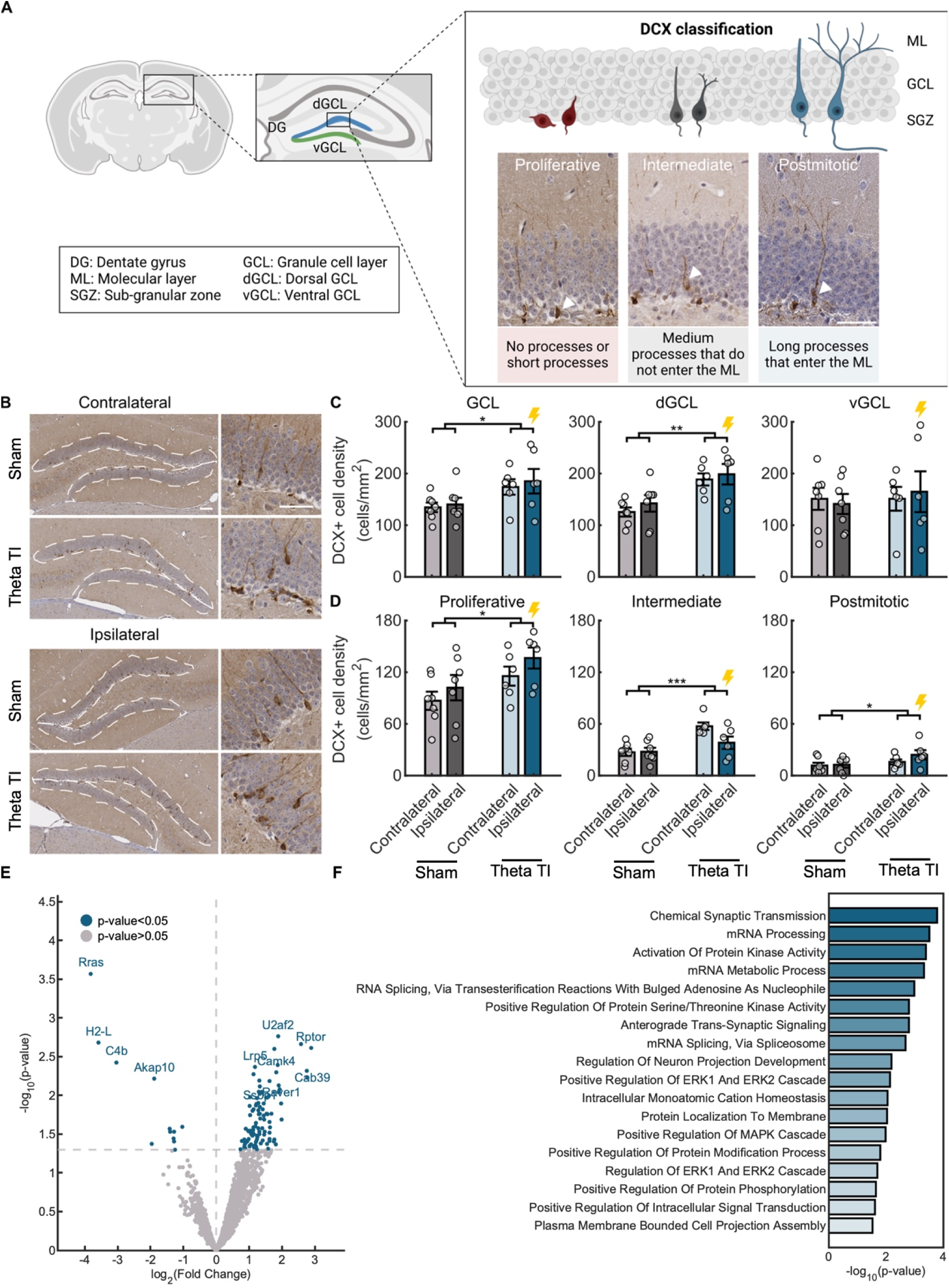
Theta TI stimulation leads to an increase in DCX+ cell density in the dorsal GCL across all maturation stages in the short term. **(A)** Schematic diagram of DCX+ maturation classification based on morphology, as described in Plümpe et al.^78^. **(B)** Representative brightfield images. Left: Sections of the contralateral and ipsilateral DG after theta-band stimulation and sham, stained with DCX (brown) and counterstained with haematoxylin (blue). Scale bar: 100 µm. Right: Zoomed-in view of the dGCL region from the images on the left. Scale bar: 50 µm. **(C)** Comparison of DCX+ cell density (in cells/mm^2^) across the DG sub-regions between conditions: granule cell layer (GCL), dorsal GCL (dGCL), ventral GCL (vGCL). GCL: F(1,11)=5.4, p=0.04, repeated-measures ANOVA, sham: n=7 mice, Theta TI: 6. dGCL: F(1,11)=14.6, p=0.0028, vGCL: F(1,11)=0.14, p=0.711, repeated-measures ANOVA, sham: n=7, Theta TI: n=6. **(D)** Comparison of DCX+ (in cells/mm^2^) across maturation stages between conditions. Proliferative (left), intermediate (middle), and postmitotic (right). Proliferative: F(1,11)=6.6, p=0.026, intermediate: F(1,11)=22.3, p=0.0006, postmitotic: F(1,11)=4.9, p=0.048, repeated-measures ANOVA, sham: n=7 mice, Theta TI: n=6. Within each bar in (C) and (D), each data point represents an individual mouse. Results are expressed as mean ± SEM. Significance was calculated using repeated measures ANOVA for all results presented in this figure (∗p<0.05, ∗∗p<0.01, ∗∗∗p<0.001). See **Table S5** for details on the statistical analysis. **(E)** Volcano plot showing proteins differentially expressed after theta-band stimulation relative to sham (total 111 proteins labelled in blue; cutoff: p<0.05). **(F)** GO enrichment of biological process of the overexpressed proteins (cutoff: p<0.05, log2(Fold Change)>0). The gene sets included contain over 3 significantly overexpressed genes with an adjusted p<0.2.

To explore the molecular pathways underlying the stimulation-induced differentiation, we performed bulk mass spectrometry on the ipsilateral DG, which underwent theta-band TI stimulation or sham. 111 proteins that were found to be differentially expressed in the DG after theta TI stimulation compared to sham (**Figure 4E**). The GO enrichment analysis for biological processes revealed increased expression of the ERK/MAPK cascade, consistent with the *in vitro* results (**Figure 1G**). Theta TI was also associated with pathways related to chemical synaptic transmission, RNA metabolic and homeostatic processes, and “regulation of neuron projection development”, among others (**Figure 4F**).

### 2.4 Long-term effects of theta-band TI stimulation on hippocampal neurogenesis in a mouse model of AD

Lastly, we aimed to explore the long-term effects of TI stimulation on AHN and its influence on hippocampal function. We repeated the experiment (2 kHz carrier with theta-band frequency difference Δf and sham), but now perfused the mice 4 weeks (instead of 1 h) after the last stimulation session (**Figure 5A**)^109,110^. To examine the fate of cells which proliferated during the stimulation period, we injected the DNA synthesis marker bromodeoxyuridine (BrdU)^79^ intraperitoneally, during the stimulation period (D3 and D10). The tissue was stained for BrdU, DCX, and the neural marker Neuron-Specific Nuclear Protein (NeuN)^80,81^. To assess hippocampal function, we performed an object pattern separation (OPS) test, known to depend on the hippocampal DG^82,83^. The OPS test started on D34, when the cells differentiated during the stimulation period were expected to undergo a critical period of enhanced synaptic plasticity (4-6 weeks post-neurogenesis initiation) ^20,84,85^.

**Figure 5.**
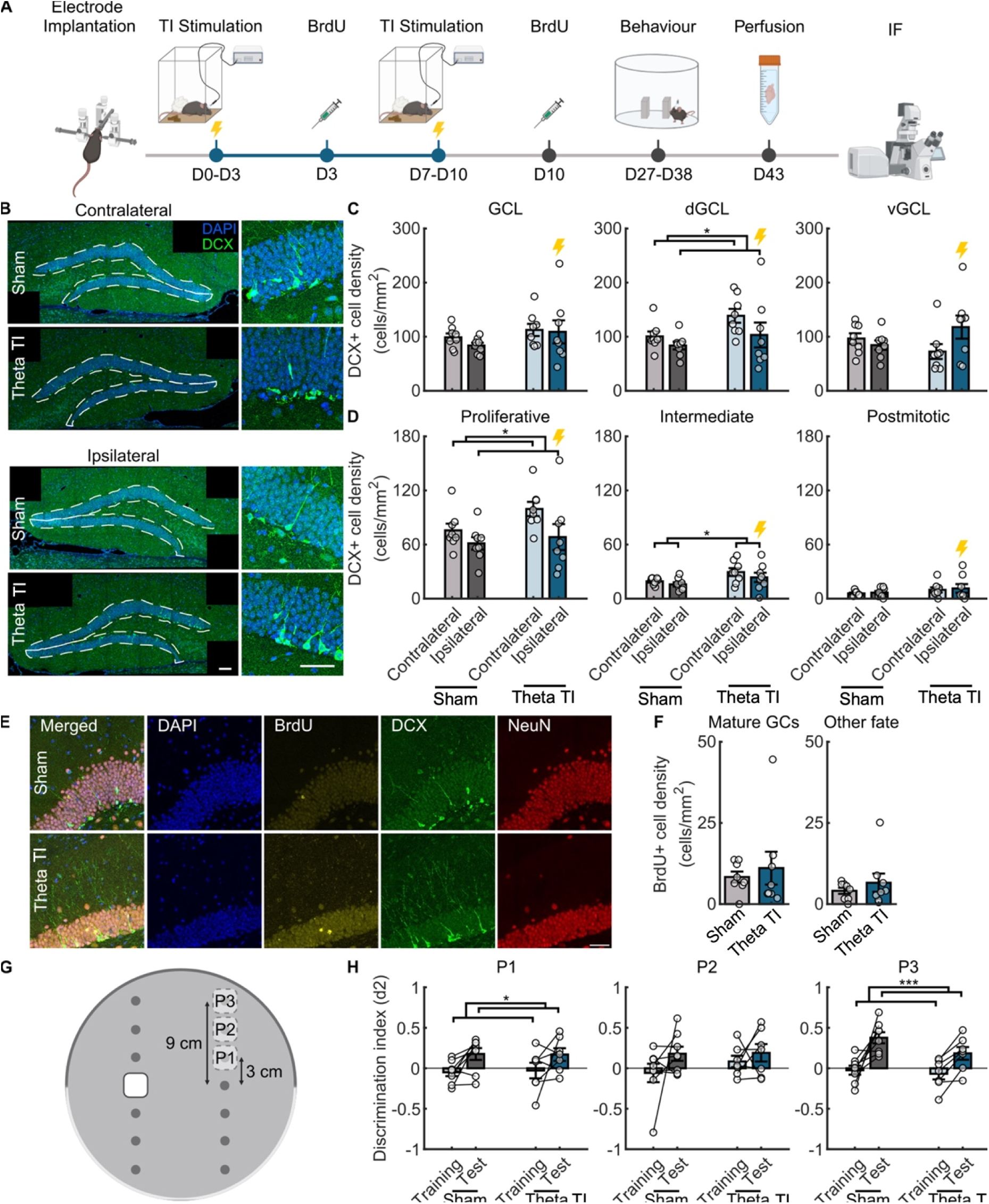
Long-term effects of theta-band TI stimulation on hippocampal neurogenesis in a mouse model of AD. **(A)** Schematic of the experimental timeline. After recovery from the surgery, stimulation was delivered to freely moving APP^N−L−GF^ mice for 8 days (days 0 to 3 and 7 to 10), 1 h/day. After the 4th and 8th stimulation sessions (days 3 and 7), all mice received a set of 4 BrdU injections, spaced 2 h apart. From days 27 to 42, mice performed the OPS task. All mice were perfused on day 43. **(B)** Representative fluorescent micrographs on the left show sections of the contralateral and ipsilateral DG after theta-band stimulation, stained with DCX (green) and counterstained with DAPI (blue), scale bar: 100 µm. Right: Zoomed-in view of the dGCL region from the images on the left, scale bar: 50 µm. **(C)** Comparison of DCX+ cell density (in cells/mm^2^) across the DG sub-regions between conditions. dGCL (contra vs ipsi): F(1,14)=5.1, p=0.041, repeated-measures ANOVA, sham: n=8, Theta TI: n=8. **(D)** Comparison of DCX+ (in cells/mm^2^) across maturation stages between conditions. Proliferative (left), intermediate (middle), and postmitotic (right). proliferative: F(1,14)=2.27, p=0.154, intermediate: F(1,14)=4.93, p=0.044, postmitotic: F(1,14)=1.31, p=0.271, repeated-measures ANOVA. proliferative (contra vs ipsi): F(1,14)=6.3, p=0.025, repeated-measures ANOVA, sham: n=8 mice, Theta TI: n=8 **(E)** Representative fluorescent micrographs showing structures stained by the antibodies BrdU (yellow), DCX (green), NeuN (red), DAPI (blue) from the GCL after theta-band stimulation and sham. Scale bar: 50 µm. **(F)** Comparison of BrdU+ cell density (in cells/mm^2^). Left: mature BrdU+ GCs. Right: BrdU+ with other (non-neuronal) fate in the dGCL after theta-band stimulation and sham. Mature DGCs: p=0.637, non-neural fate: p=0.875, Wilcoxon rank-sum test, sham: n=8 mice, Theta TI: n=8 **(G)** Schematic of the object pattern separation (OPS) task showing the three possible positions to which the non-static object could be moved (P1, P2 and P3). **(H)** Comparison of discrimination index (d2) for the OPS training and test sessions after theta-band stimulation and sham, obtained in positions P1 (left), P2 (middle) and P3 (right). Training vs test: P1: F(1,13)=8.3, p=0.013, P3: F(1,13)=25.2, p=0.00024, repeated-measures ANOVA. Theta TI vs sham: P1: F(1,13)=0.009, p=0.926, P3: F(1,13)=2.85, p=0.115, repeated-measures ANOVA, sham: n=8, Theta TI: 7 Within each bar, each data point represents an individual mouse in (C), (D), (F) and (H). Results are expressed as mean ± SEM. Significance was calculated using repeated measures ANOVA for the results presented in (C), (D) and (H). Two-sided unpaired two-samples Wilcoxon rank-sum test was used for the results presented in (F). (sham: n=8 mice, Theta TI: n=7, ∗p<0.05, ∗∗p<0.01, ∗∗∗p<0.001). See **Table S6** for details on the statistical analysis.

Four weeks after the end of the stimulation period, a residual increase in the expression of the differentiation marker DCX was still detected. Specifically, the ipsilateral and contralateral GCLs combined had a higher density of DCX+ cells at the intermediate maturation stage after theta TI stimulation compared to sham. No changes were observed at the proliferative or postmitotic stages in the dGCL (**Figure 5D**). As opposed to the short-term study, there was no difference in DCX+ cell density when segmenting the area in the GCL, dGCL or vGCL subregions (**Figure 5B-C**). However, a lower density of DCX+ cells was observed in the ipsilateral GCL compared to the contralateral GCL, in both theta TI stimulation and sham conditions, suggesting a potential effect of the minimally-invasive implantation surgery (**Figure 5C-D**).

We did not find differences between conditions in BrdU+ cells in the dGCL with a mature neuronal phenotype (i.e., BrdU+/NeuN+/DCX-cells^80,81^) or other phenotypes (i.e., BrdU+/NeuN-/DCX-cells^80,81^) at the 4-week post-stimulation time point (**Figure 5E-F**). No difference was detected in BrdU+ cell density in the hilus area, where ectopic differentiation may occur^86,87^ (**Figure S4**). These results could indicate that theta-band TI stimulation may have affected the survival of differentiating NPCs rather than the proliferation or differentiation process. However, the small number of BrdU+ cells detected lowered the sensitivity of the quantifications, rendering robust conclusions challenging. The performance of animals in the OPS test did not change with TI stimulation. However, the mice showed a higher discrimination index (d2) in the easiest (P1) and the hardest (P3) object positions, after both theta-TI stimulation and sham (**Figure 5G-H**), which is consistent with previous reports^88^.

## 3. DISCUSSION

Neural tissue repair strategies for neurodegenerative diseases have been facing critical challenges, due to the affected areas being located in the deep-brain. Here, we aimed to tackle this gap by using TI electrical stimulation as a non-invasive, drug-free tool to augment neurogenesis in the deep brain. Our results demonstrated that TI stimulation can be used to enhance neural differentiation in both embryonic and adult NPCs.

The neurogenic effect of TI stimulation was specific to frequency differences (Δf) at the theta-band, both in embryonic NPCs *in vitro* and in AHN *in vivo*. Theta band oscillation is a hallmark of early brain activity, emerging spontaneously as ‘spindle bursts’^89–92^ and shaping development through sensory-driven processes^90,92–95^. The presence of endogenous theta oscillations in developing brain organoids^96^ and adult brains^149–15^ further suggests their conserved role in neurogenesis across life stages. Such knowledge, combined with our findings on different NPC models, may entail that neural stem cells at different developmental stages respond to theta oscillatory patterns via a common mechanism. Earlier studies showed the neurogenic effects of electrical stimulation^29,30,97–100^. In particular, 100-130 Hz is the most widely employed frequency to increase neural differentiation *in vitro* and *in vivo* because of its widespread use in clinical DBS^29,30,97–107^. Theta stimulation has been reported to increase neural differentiation *in vitro*^108,109^ and *in vivo*, delivering DBS to rat models of PD^105^ and transcranial magnetic stimulation (TMS) in Swiss Webster^105^ mice. Other non-invasive models achieved increases in AHN, such as 40 Hz transcranial alternating current stimulation (tACS) in a 5xFAD mouse model^43^, and 40 Hz transcranial magnetic stimulation (TMS)^44^, transcranial direct current stimulation (tDCS)^110^ and 40 Hz audiovisual stimulation in WT mice^111^. Audiovisual gamma stimulation has been recently reported to increase adult neurogenesis in wild type^111^ and Down syndrome^112^ mice. However, no consensus has been reached on the optimal stimulation parameters to promote neural regeneration, largely because of the lack of systematic studies, and the variability between protocols and setups used. To allow for direct comparisons among protocols, we developed a standardized and electrically stable system for *in vitro* TI stimulation, and systematically assessed the influence of a range of TI frequencies in parallel. Our results showed that TI at the theta frequency exerts a neurogenic effect and increases metabolic rate in monolayer cultures of primary embryonic NPCs *in vitro*, as observed from independent increases in β-tubulin III expression and calcium signaling dynamics. The same embryonic cells grown in 3D neurospheres and encapsulated in a bioactive hydrogel did not display increases in β-tubulin III expression, likely because of the longer neural differentiation timelines of 3D *in vitro* cultures ^64–67,113,114^. However, the increased correlation between β-tubulin III and Map2 expression may suggest enhanced neural maturation.

Theta TI stimulation also boosted AHN *in vivo*. As AHN is known to be impaired in subjects with AD and mild cognitive decline (postmortem), it has been proposed as a therapeutic target to support hippocampal function and delay cognitive decline^19,22,23^. We employed a mouse model of AD, as opposed to WT mice, to provide a proof of concept of neural recovery in a diseased context with direct clinical translation. Theta TI stimulation of the hippocampus enhanced the differentiation of adult NPCs in both the stimulated ipsilateral and contralateral hippocampal regions. Such a bilateral response may be due to the electrical stimulation activating the contralateral hippocampus due to interhemispheric interconnectivity^115,116^. The theta TI stimulation may also affect the survival of differentiating NPCs. It is known that almost half of the NPCs in the DG undergoing neurogenesis do not survive to full differentiation (requiring 6-8 weeks in mice)^20,84,85^. Our results showed that theta TI hippocampal stimulation increases the expression of both proliferation and differentiation markers (Ki67 and DCX, respectively) in APP^N-L-GF^ mice immediately after the two weeks-stimulation session. A residual increase in the differentiation marker DCX was observed four weeks after the stimulation ended (six weeks from onset), suggesting that theta-TI stimulation may not only promote cell differentiation, similarly to the *in vitro* results, but also cell survival. This increase was specifically observed in proximity to the TI stimulation target (CA1), which could be indicative of a localised effect. Importantly, while it is likely that Ki67+ cells were undergoing initial stages of neurogenesis, it remains possible that the increase in proliferation was not specific to AHN cells^117^. For example, the proliferation increase may be related to the observed increase in microglia (via Iba1 expression).

Four weeks after stimulation ended, the TI-induced enhancement of DCX+ NPCs differentiation was not reflected in an increase in BrdU+ cell density, or improvements in hippocampal-dependent memory. Thus, it is also plausible that NPCs that displayed augmented differentiation at the end of the stimulation period did not survive the maturation process later, which would explain the lack of behavioural improvements. It is also possible that theta TI did not promote the generation of new neurons, but it only supported neuronal development. However, this is unlikely considering the increase in proliferation observed with Ki67. The lack of a significant increase in BrdU+ cells at the 4-week post-stimulation time point may be due to limitations in the methodology: the very low number of BrdU+ cells detected may have impaired the sensitivity of the quantification. In addition, induced AHN increases may become impaired in APP^NL-G-F^ models at 8-9 months^81^. Because this is also the age of our long-term study animals, changes in TI-induced neurogenesis may also have been affected.

Together, these results indicate that theta-TI stimulation can influence NPC behaviour *in vitro*, and support early-stage AHN and cell survival *in vivo*. Although cells undergoing adult and embryonic neurogenesis significantly differ in function and phenotype, the gene expression pathways and electrophysiological properties involved in neural differentiation show significant similarities^118^. However, despite recent discoveries^91,93^, the developmental pathways underlying oscillatory activity remain largely unknown. Our results suggest a potential common response to theta-frequency neuromodulation across species, models, and developmental stages. This investigation reports that the MAPK/ERK1 and 2 cascades are overexpressed upon theta-TI stimulation, in both embryonic NPCs *in vitro*, and in AHN in an AD *in vivo* environment. The MAPK family is highly conserved across species, and it contains extracellular signal-regulated kinases (ERKs), which transduce environmental signals to the nucleus upon phosphorylation, regulating cell proliferation, differentiation and survival^119,120^. ERK1/2 are known to be involved in neural stem cell differentiation, and their suppression leads to developmental impairments^119,120^. The upregulation of these kinases has been previously correlated with electrical activity and electrical stimulation, via binding to upregulated growth factors such as the epidermal growth factor receptor (EGF receptor) and the brain-derived neurotrophic factor (BDNF), or via ligand-independent activation triggered by electrical stimulation^30,63,92,121–123^. BDNF is an immediate upstream regulator of ERK1/2 via tyrosine phosphorylation^124^, and it is known to play a role in stimulation-induced neurogenesis, differentiation, and plasticity both *in vitro*^29,125^ and *in vivo*^111,126–131^. Although additional evidence and causation correlations are required to establish a link between the ERK1/2 pathway and the theta-TI stimulation, our results hint to a species- and development-conserved neurogenic mechanism, activated by the theta-TI stimulation and driven by the ERK1/2 cascade.

TI stimulation modulates the neural membrane at the frequency difference Δf ^132–135^. Hence, it is conceivable that the theta TI stimulation induced periodic changes in the NPCs membrane potential at the theta-band, that in turn modulated the ion influx rate, known to impact proliferation and differentiation^27,28^. Yet, the distinct electrophysiological properties of NPCs compared to neurons may affect their neuromodulation dynamics^20,136–139^. To the best of our knowledge, no previous study has looked at the effect of TI or kHz frequency stimulation on neural stem cells or NPCs. The neural membrane response to TI is primarily affected by the time-responses of voltage-gated sodium and potassium channels^133,135^. TI and kHz frequencies may have a different influence on membrane depolarization dynamics in NPCs, due to the expression patterns of ion channels during neural differentiation^140–142,20,137,136,139^. Such properties imply that neural stem cells may be able to follow oscillations at higher frequencies and be influenced by both low and high frequency stimulation simultaneously. Here, the carrier control (1 kHz) was not associated with any change in NPC differentiation or proliferation, although a combined effect of the two frequencies could not be excluded because an AC (non-TI) theta condition was not investigated. Further insight is required to elucidate the electrophysiological response of neural stem cells and NPCs to TI and kHz stimulation at the cellular and sub-cellular level.

We cannot exclude that the observed neurogenic phenomena and upregulated pathways are caused by mechanisms independent from NPC membrane oscillations. In this study, all models consisted of a hetero-cellular population, where non-neuronal cell types might respond to the stimulation and in turn, influence neural differentiation. For example, astrocytes are known to participate actively in brain signalling^53,54,143^ and to respond to electrical stimulation^144–146^. Our results showed a stimulation-induced increase in GFAP expression in the 3D *in vitro* model, along with increased neuronal maturation. Although further evidence is required, theta TI stimulation could have influenced the phenotype of astrocytes in the culture via increased glutamate release, increased calcium signaling, or other mechanisms^55,147,148^. Ahtiainen *et al.*^149^ have recently reported that the electrophysiological response of primary rat neurons to theta TI stimulation *in vitro* was suppressed when stimulating a co-culture of neurons and astrocytes (1:1 ratio), suggesting a key role of glial cells in the neuromodulatory response to TI stimulation. The increased expression of GFAP in the 3D NPC model upon theta TI stimulation might have also accelerated glial differentiation. Although hydrogel functionalization with the IKVAV epitope is known to promote neural differentiation and inhibition of gliogenesis^150–152^, we observed a large increase in astrocyte presence along with an enhanced neuronal maturation. This result may entail that the 3D structure combined with electrical stimulation enhanced glial development or astrocytic activation^153^, prompting further investigations. The increase in AHN observed *in vivo* might also be related to the neuromodulation of non-neuronal cell types. Flickering-light^154^ and transcranial AC^155^ 40 Hz stimulation can influence microglia phenotype in AD models, demonstrating the potential of modulating non-neuronal cell types. Microglia activation could also explain the overexpression of the MAPK pathway^156^, which could be correlated with the increase in Iba1 expression detected at the theta frequency in our *in vivo* study. It is also plausible that the TI-induced neurogenesis effect is mediated by endogenous neuronal stimulation, which, in turn, may influence NPC differentiation. For example, several studies have shown that parvalbumin (PV) interneurons regulate adult neurogenesis via GABAergic signalling^157,158^. Hippocampal PV interneurons control and preferentially respond to theta oscillations^159,160^, which could explain the frequency-specific neurogenic effect we observed. Future investigations on the influence of theta TI stimulation on NPCs *in vitro* and *in vivo,* using single-cell omics techniques and electrophysiology, could deepen the knowledge on this mechanism. In addition, replicating the results on a more translational *in vitro* model, such as using human induced pluripotent stem cell (iPSC)-derived cultures or organoids, would improve the translatability of the approach and detect more human-relevant molecular pathways.

These results highlight the importance of a bio-inspired approach in neuromodulation for regenerative medicine. The interventional field has currently explored stimulation parameters and protocols based on historical applications and studies. Conversely, neurodevelopmental investigations have provided knowledge on the biological mechanisms generating and responding to endogenous activity. Here, we propose to utilize bio-mimetic stimulation protocols to recapitulate species- and phase-specific endogenous activity, rooting experimental designs on current knowledge of neurodevelopmental activity patterns.

Overall, this initial evidence on the neurogenic potential of theta TI stimulation positions this methodology as a promising, non-pharmacological approach to bridge a critical translational gap in brain regeneration for neurodegenerative diseases. Although electrical stimulation is known to augment neurogenesis, previous technologies could not reach deep brain structures with spatial precision and without surgical electrode implantations. TI stimulation allows for non-invasive, focal electrical stimulation of deep brain structures in humans^47,161^, strengthening the translational potential of this approach^47,161^. For example, TI may complement existing interventions, such as cell replacement therapies, or be used to enhance adult neurogenesis. As the field shifts toward multimodal, individualized interventions, TI could emerge as both a therapeutic tool and a platform for probing stem cell behavior in response to bioelectric cues in the deep brain.

## 4. LIMITATIONS OF THE STUDY

While our results highlight the neurogenic potential of theta-band TI stimulation *in vitro* and *in vivo*, some limitations should be acknowledged. First, the *in vitro* and *in vivo* stimulation models used are different in species, developmental stage and related stimulation timelines. This hinders robust comparisons between models, which was beyond the scope of this investigation. The rationale for using different models was strengthening the evidence on the neurogenic effect of TI stimulation across species. The stimulation protocols and timelines were adapted to recapitulate the endogenous waveforms of the model selected, and the biomarkers were chosen based on the most established characterizations for the developmental stage and conditions considered. Future studies utilizing the same quantification methods on additional time points and models, would elucidate the biological similarities in neurogenic responses to TI across species.

Additionally, we did not directly confirm focal stimulation of the hippocampal CA1 region *in vivo*. We investigated the expression of c-Fos, a neural activity marker which has been previously reported in correlation with acute TI hippocampal stimulation^46^. However, we could not draw solid conclusions due to the sparse expression of the marker and the lack of knowledge on the influence of chronic TI stimulation, for which a dedicated investigation is required. Direct electrophysiological recordings are needed to confirm targeted stimulation, and whole-brain readouts are required to fully rule out off-target effects. However, our electrode configuration was closely adapted from previous work, where hippocampal targeting was validated with electrophysiological recordings^162,163^.

The observation of higher hippocampal DCX and Ki67 expression suggests augmentation of endogenous NPCs proliferation and differentiation. However, we cannot exclude the possibility of de-maturation of hippocampal neural cells that re-express DCX and Ki67^87,164^. Nevertheless, the increase in proliferative and intermediate-stage NPCs, along with the absence of ectopic differentiation in the hilus, renders the probability of de-maturation low. The lack of improvement in the hippocampal memory test suggests that longer stimulation periods may be needed to induce behavioral benefits^187,188,189,190,117,186^. The use of an AD mouse model may inherently limit the neurogenic response and, in turn, the sensitivity of the experimental tests, especially at later disease stages^81^. Therefore, repeating the experiments in wild-type mice would further validate the findings and improve translatability.

## 5. METHOD DETAILS

### Primary NPCs

#### Dissociation of eNPCs

Primary cultures were obtained from the ventral-mesencephalon (VM) of Sprague-Dawley rat embryos. These cells were selected as an established model to recapitulate neural differentiation *in vitro*^165^. Tissue dissociation was first performed as described by Thompson and Parish^166^. Briefly, E14 were extracted from the embryonic sac and dissected in cold Dulbecco’s phosphate buffered saline (DPBS) under a stereomicroscope. The midbrain was carefully dissected from the embryo and the neural tube was opened to expose the ventral region. The surrounding vasculature and connective meningeal tissue were carefully removed from the VM midline to eliminate as many contaminant cells as possible. The dissected tissues were then incubated for 5 min at 37°C in 0.1% trypsin and mechanically dissociated with a 1 ml pipette tip, followed by a 23 G needle. Single cells were then counted and plated in 24-well plates coated with Poly-L-lysine (PLL) for 20 min, rinsed twice with DPBS and air dried.

#### *In Vitro* Culture of Primary eNPCs

Cells were plated in 24-well plates at a density of 60,000 cells/cm^2^ with 0.75 ml of proliferation medium (DMEM)/F12 supplemented with 33 mM D-glucose, 1% L-glutamine, 1% penicillin/streptomycin (P/S), 2% B27, 1% fetal bovine serum (FBS), 20 ng/ml EGF, 20 ng/ml FGF). The cultures were then gradually transitioned from proliferation medium to differentiation medium: Dulbecco’s modified Eagle’s medium (DMEM)/F12 supplemented with 33 mM D-glucose, 1% L-glutamine, 1% penicillin/streptomycin (P/S), 50 mg/ml BSA, 1% N2. 75% of the medium was replaced every 2 days, increasing the ratio of differentiation medium by a third at every first medium change, until the transition was complete. Three pregnant rats were dissected for the monolayer NPC experiment (**Figure 1**), and three tissue culture wells per condition (technical replicas) were cultured and quantified, to ensure replicability and reduce variability in the dataset.

For experiments involving NPC-derived NS, cells were plated in ultra-low adhesion T75 flasks at a density of 1.5 − 2 · 10^5^ cells/ml in proliferation medium. NS cultures were maintained *in vitro* for 3-4 days until an average diameter of 50-100 μm was achieved. The NS were then encapsulated as described below, and transitioned from proliferation medium to differentiation in the same way as the monolayer cultures. Four pregnant rats were dissected for the 3D experiment (**Figure 2**), and 2-3 tissue culture wells per condition (technical replicas) were cultured and quantified. The 3D constructs which moved from their initial positioning during the experiment were discarded from the quantification because of the different stimulation voltage they may have experienced by moving across the well.

In all the *in vitro* experiments, we discarded the tissue culture wells where the recorded stimulation voltage did not meet the inclusion criteria to minimise variability and DC offset (described below).

### Animals

Homozygous APP^N−L−GF^ mice (RRID:IMSRRBRC06344, Charles River Laboratories, United States) were used in all *in vivo* experiments^73^. For the short-term *in vivo* study (**Figure 3**, **Figure 4**), male and female 5–6-month-old homozygous APP^N−L−GF^ mice (n=27; 8m, 19f) were assigned to four experimental groups: sham (control, n=7, 2m, 5f), delta (1 Hz stimulation, n=4, 1m, 3f), theta (8 Hz, n=6, 2m, 4f) and gamma (40 Hz, n=5, 2m, 3f). The animals were divided into five cohorts of mice for experimental procedures, each containing 4-7 mice. Mice from this study were donated by the laboratory of Prof Karen Duff at UCL. For the long-term *in vivo* study (**Figure 5**), 5-6 months old homozygous APP^N−L−GF^ mice (n=16; 11m, 5f) were assigned to two experimental groups: sham (control, n=8, 6m, 2f) and theta (8 Hz, n=8, 5m, 3f). Mice were split into 2 cohorts for experiments, with each containing 8 subjects. Mice from this study were bred at our isolator at Charles River Laboratories and derived from the breeding colony established by collaborators.

All mice were single housed after surgery and maintained on a 12 h light: 12 h dark cycle with ad libitum access to water and food. Experiments were approved by the Animal Welfare and Ethical Review Body (AWERB) at Imperial College and conformed to the UK Animals (Scientific Procedures) Act 1986.

### Electrical stimulation *in vitro*

#### *In vitro* electrical stimulation device

An *in vitro* stimulation device was used to deliver the TI stimulation frequencies on the monolayer NPCs and the 3D model. The format of the device consisted of a custom lid designed for a standard 24-well tissue culture plate (Corning Costar, United States). The tip of two insulated Pt wire electrodes were exposed to the culture medium at symmetrical distances in the well. Pt was selected as the electrode material for its electrical stability and cytocompatibility. The case containing the Pt electrodes and the electrical connections to deliver ES was made with computerised numerical control (CNC)-machined PEEK, due to its stability at high temperatures, pressure, and humidity, which are required for autoclave sterilisation^167^.

The device was designed with 18 independent stimulation channels, with one channel per culture well. The 2 Pt-wire electrodes were positioned to fit into each culture well through two PEEK electrode holders, which were built into the device to keep them in place and isolate them from the culture medium (**Figure S5**). Pt wires were then connected to high temperature-bearing cables (Farnell, 0047105) through metal crimps (RS Components, 670-6290) inside of the device. The connections were arranged along the internal wells of the device and covered in medical-grade silicone (NuSil, MED4-4220) to avoid water contact and damage to the connections. The internal cables were then soldered to autoclavable six-pin connectors (RS Components, 536-5605). All connections and all non-exposed electrode surfaces were insulated using medical-grade silicone.

#### Electrical stimulation protocol

The device was connected to a multichannel stimulator (STG4002-1.6 mA, Multichannel Systems, Reutlingen, Germany) through multiwire cables and compatible connector plugs (RS Components, 536-5706). The stimulation paradigm was created and delivered via the Software Multichannel Systems II (Multichannel Systems, Reutlingen, Germany). The stimulation applied was current-driven to minimise the damaging effects of charge injection. A capacitor was added in line to both the stimulation and ground electrodes to remove the DC offset artifacts^168^. The voltage was measured with an oscilloscope at the start and at the end of the stimulation to confirm the stability of the connections over time.

The eNPCs were stimulated for three days (DIV 7, 8, 9) and then fixed or imaged at DIV 10. The stimulation was delivered at a current of 1 mA (current density: 1.91 mA/mm^2^) for 6h per day, with 2s stimulation periods and 2s resting periods (2s on-2s off). To model the TI stimulation and deliver it through one stimulation channel, two carrier frequency waves (F_1_ and F_2=_F_1_ + ΔF) were digitally summed in anti-phase using MATLAB before being delivered to the cells, as in Grossman et al.^46^ (**Equation 1**):

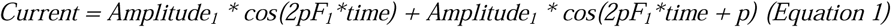

Here, f_1_=1000 Hz (carrier frequency), and f_2_=f_1_ + Δf, where Δf corresponds to one of the TI frequencies selected. The selected TI frequencies for stimulation in the monolayer NPC study were 10 Hz (theta burst), 40 Hz (low gamma) and 100 Hz (high gamma). The carrier frequency (1000 Hz sinusoid) was used as a control for the high frequencies. Only the theta burst stimulation was delivered in the 3D model study. The stimulation amplitudes were set to 2 mA peak-to-peak for both interfering waveforms. A Multichannel System Stimulator (STG4008 – 1.6mA from Multichannel Systems, Reutlingen, Germany) was used to deliver the stimulation.

Two 200 nF capacitors were added in line with each stimulation channel to eliminate possible DC offsets and the stimulation cables were shielded from environmental noise by grounded aluminium foil wrapping. The voltage of each channel was measured at the beginning and at the end of the stimulation with an oscilloscope to ensure the voltage variability was under 25% and the DC offset was under 10% of the peak-to-peak voltage. If the criteria were not met, the sample was excluded from the analysis.

### 3D In Vitro model

#### Synthesis of the Self-Assembling Hydrogel

A self-assembling hydrogel was used because of its mechanical properties recapitulating the brain environment^152^. The gel was functionalised with the laminin-derived epitope Isoleucin, lysin, valin, alanin, valine (IKVAV) because of it is known to support neural differentiation^169^. The hydrogel (molecular formula: Palmitoyl-VVAAEEEEGIKVAV-COOH) was produced via solid-phase peptide synthesis (SPPS). Briefly, a Wang resin pre-loaded with Valine was swollen in a 10-mL syringe with dichloromethane (DCM) for 10 mins on an action shaker. For the deprotection step, a 20%-piperidine in N-Methyl-2-pyrrolidone (NMP) solution was added to the syringe and shaken twice. The resin beads were washed with NMP 3 times, once with DCM, once with methanol and once again with DCM after every deprotection and coupling step. For the coupling step, 3 eq of each amino acid were dissolved in 9 eq of 1-[Bis(dimethylamino)methylene]-1H-1,2,3-triazolo[4,5-b]pyridinium 3-oxide hexafluorophosphate (HATU) and 5 eq of N,N-Diisopropylethylamine (DIPEA) in an excess of NMP and shaken for a minimum of 1 h. For the final palmitol coupling, 8 eq of Palmitol, 8 eq of HATU and 12 eq of DIPEA were mixed instead. For the cleavage of the peptide from the resin, a solution of 95% trifluoroacetic acid (TFA), 2.5% H20 and 2.5% triisopropyl silane (TIPS) was shaken in the syringe for 2.5 h. The TFA was subsequently evaporated from the cleavage solution under nitrogen gas or with a rotovap. The peptide was precipitated in cold diethyl ether, vacuum-filtered and dissolved in MilliQ water before freeze-drying. The above steps were performed at room temperature (15-25°C). The PA-IKVAV was purified by high-performance liquid chromatography (Analytical HPLC, Agilent Technologies 6130 Quadrupole LC/MS, United States) and the final purity was validated with liquid chromatography–mass spectrometry (LC/MS, Agilent Technologies, United States).

#### Encapsulation of NPC-derived neurospheres

The purified hydrogel was used at 1% w/v and dissolved in 80% neutral solution and 20% cell solution. First, the peptide was dissolved in the neutral solution (150 mM NaCl and 3 mM KCl in DI water). The pH was adjusted to 7.2-7.4 with 1% NH_4_OH solution and the solution was heated to 80°C for 30 min for annealing. After cooling to room temperature, the NS suspension was mixed into a density of 60,000 NS/mL. To form each gel, 20 μL of the solution were added to a circular PDMS mold (5 mm diameter) placed on a 13 mm round glass coverslip in a 24-well plate. These glass coverslips were coated with CaCl_2_ crystals via overnight incubation in gelling solution (neutral solution supplemented with 50 mM CaCl_2_) at 37°C. Lastly, the gelling solution was sprayed on the gels 15 times (approximately 10 mL) with a spray bottle to ensure complete gelation and the gels were incubated in diluted gelling solution (neutral solution with 25 mM CaCl_2_) for 1 min at room temperature. The gels were washed with DPBS and covered in 750 μL of cell medium.

### In Vitro Assays

#### Assessment of Metabolic Activity

The metabolic activity of cultures was evaluated using a commercial Alamar Blue assay (ThermoFisher, DAL1025). Samples were incubated for 5 h in 10 % AlamarBlue solution in differentiation medium. The supernatant was collected and analysed using a plate reader (544-590 nm). The absorbance (A) values difference was calculated as a proxy for the total metabolic reduction in the well. The values were then normalized by the ratio between the difference in absorbance between the positive control (Alamar Blue-only well, AB) and negative controls (no Alamar Blue, C) at 540 nm and at 590 nm (**Equation 2**):

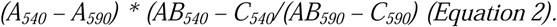

#### Immunofluorescent staining

Samples were stained via indirect double-immunofluorescent labelling using antibodies against relevant biomarkers. Briefly, samples were fixed with 4% paraformaldehyde (PFA) for 20 min at room temperature. Cell cultures were then washed twice in DPBS and incubated in cold permeabilization buffer (0.5 ml Triton X-100, 10.3 g sucrose, 0.292 g NaCl, 0.06 g MgCl2, 0.476 g HEPES buffer, in 100 ml water, pH 7.2) for 5 min at room temperature.

For monolayer experiments, non-specific binding sites were blocked with 1% BSA for 30 min at 37 °C, and samples were incubated at 4°C overnight with a chicken anti-GFAP+ antibody (1:800, Abcam 4674), rabbit anti-β tubulin III antibody (1:800, ThermoFisher Scientific T2200), and mouse anti-Sox2 antibody (1:500, Abcam 79351). Following incubation with the primary antibodies, samples were washed 3 times with 0.05% Tween-20/DPBS and incubated at room temperature for 1 h with Alexa Fluor 647 goat anti-Chicken IgY (H+L) (1:1000, ThermoFisher Scientific A-21449), Alexa Fluor 594 goat anti-Rabbit IgG (H+L) (1:1000, ThermoFisher Scientific A-11012) and Alexa Fluor 488 goat anti-Mouse IgG (H+L) (1:500, ThermoFisher Scientific A-10667) antibodies.

For encapsulated neurospheres, non-specific binding sites were blocked with 5% BSA for 4 h at room temperature, and samples were incubated at room temperature for 24h in 3% BSA with two sets of antibodies: a cell-specific panel and a functionality panel. The cell-specific panel included the same antibodies as the 2D experiments and the functionality panel contained a chicken anti-Map2 antibody (1:800, PA1-100), rabbit anti-synaptophysin antibody (1:400, Abcam 52636), and mouse anti-β tubulin III antibody (1:800, Abcam 78078). Following incubation with the primary antibodies, samples were washed 3 times with 0.05% Tween-20/DPBS for 10 min on a shaker and incubated at room temperature overnight with the same secondary antibodies as the 2D samples (concentrations in Table 4.2). After washing 3 times with DPBS, cell nuclei were counterstained with Hoechst 33342 (1:2000, ThermoFisher Scientific 62249), and samples were mounted on glass slides.

#### Immunofluorescence Microscopy and Image Analysis

Samples were imaged using a Leica SP8 inverted confocal microscope with a fixed scan of 1024×1024 pixels. On average, 3 z-stacked images (1 - 1.5 μm thickness) at 20x magnification were acquired for each 2D sample, and 3 z-stacked images at 20x and 40x were acquired from the 3D cell-type panel and cell-functionality panel, respectively. Image processing and analysis was performed using the ImageJ software (National Institutes of Health, United States) in combination with MATLAB (Release 2023b, The MathWorks, Inc., Natick, Massachusetts, United States). For image processing, a gaussian filter was applied for noise reduction, and a watershed function was applied when quantifying nuclear staining. The number of nuclei was quantified from the Hoechst channel and divided by the image area to determine cell density. Biomarker expression was calculated by applying a pre-defined threshold to all images. The co-localization was derived from the Pearson’s co-localization coefficient^170^.

#### mRNA extraction and Sequencing

The mRNA was extracted using a commercial RNeasy Micro Kit (Cat. 74004, Quiagen). Briefly, cells were washed with DPBS and lysed using Buffer RLT. The lysate was homogenised by centrifugation at full speed in a QIAshredder spin column for 2 mins on ice. Then, a 70% ethanol solution was added to the lysate and centrifuged in a RNeasy Minelute column for 15 s at >8000xg. After washing the column membrane with Buffer RW1, the RNeasy MinElute spin column membrane was incubated with DNase I mix (12.5% DNase I stock solution to 87.5% Buffer RDD) for 15 min at room temperature to remove DNA contamination. The membrane was subsequently washed once with Buffer RW1, once with Buffer RPE and once with 80% ethanol. The RNeasy MinElute spin column was then centrifuged at full speed for 5 min to allow for ethanol evaporation. Lastly, RNase-free water was added to the spin column and centrifuged for 1 min at full speed to elute the RNA. To quantify mRNA content and purity, the eluate absorbance at 260 nm, 280 nm, and 230 nm was measured using a Implen NanoPhotometer N60/N50 Nanodrop. Only samples with a purity ratio above 1.9 were included in the dataset (**Figure S7**).

The library preparation, sequencing, and analysis was performed in collaboration with Novogene (Novogene UK Company Limited, United Kingdom). Poly(A) enrichment was utilised for library preparation followed by 150 bp paired-end (40 million reads) sequencing using an Illumina Sequencing PE150 system. The data quality control was performed using a base calling error rate distribution of Q30 (99.9%), a GC content distribution control (with cut-off at 50%), and non-clean reads were removed from the analysis (defined as containing adapter contamination, a percentage of uncertain nucleotides >10%, and a percentage of low-quality nucleotides >50%). The reference genome used was mRatBN7.2 available from Ensembl^171^.

The expression analysis pipeline was carried out by first determining the gene expression level distribution, followed by a Pearson’s correlation analysis, and a principal component analysis (PCA) on the gene expression value (Fragments Per Kilobase of transcript per Million mapped reads, FPKM) of all samples^172^. The gene expression cut-off for the Venn diagram was set at 1 FPKM. The DESeq2 software was used for differential gene expression analysis^173^. For this, the DESeq2 normalisation method was applied, the p-value calculation model followed a negative binomial distribution, and the false discovery rate (FDR) method was Benjamini-Hochberg^174^. The Enrichr website^175^ was used to implement the GO enrichment analysis of biological process (2023), with a cut-off of p <0.05 and log_2_(Fold Change)>0.

#### Calcium Activity Imaging and Analysis

Calcium imaging was performed using a Fluo-4 AM assay (ThermoFisher Scientific F14201). Samples were washed with DPBS and incubated for 1 h in the staining solution (1 μM Fluo-4 diluted in recording medium, composed of HBSS supplemented with 128 mM NaCl, 1 mM MgCl2, 45 mM sucrose, 10 mM glucose, and 0.01 M HEPES). The staining solution was removed, and samples were washed with fresh recording medium and then imaged on a Zeiss Axio Observer widefield fluorescence microscope at 37 °C and 5% CO2 using a 10x air objective. Each ROI was imaged for 10 min with an acquisition rate of 8.7 Hz and at least two ROIs per sample were imaged. The overall imaging time did not exceed 60 min and the imaging order of the samples was randomised.

For the analysis of the recordings, the first step was correcting the photobleaching, defined as the reduction of the fluorophore fluorescence due to photon-induced chemical damage during imaging^176^. To correct for this effect, an exponential curve was fitted to the signal derived from the mean of each frame and the derived points were subtracted from the corresponding frame of the recording. Subsequently, the cells showing calcium activation were detected by selecting all the pixels with a standard deviation of their signal over time higher than twice the mean of the standard deviation. The values were then manually adjusted to keep the pixel range selected in a range of 0.3-0.6% of the total image area. This threshold value was selected by optimising the parameter to detect visible signals. All the detected pixels were then grouped, and all ROIs with an area lower than 20 μm^2^ were excluded from the analysis, based on a visual observation. The traces of each ROI were computed as the average signal from all pixels in the ROI. A moving average of 5 frames and a low pass-filter (normalised Nyquist frequency: 0.03) was applied for smoothing the visualisation. The peaks were manually selected by two blinded experimenters, based on the following criteria: the trace should display a one or more clear peaks with one ascending and one descending sides, each peak should display one maximum value, without noise at the peak, the peak should be shorter than 120 s, the peak should also be visible in the trace derived from the signal normalised by the average of the frame for each frame. These criteria allowed for a conservative selection of the peaks, which was based on previously reported calcium transients of NPCs^58,60^. The peaks were automatically detected by finding a local maximum^177^. The peak width was extracted from the traces under optimised conditions (minimum height: 50 u, Minimum width: 10 frames, Minimum horizontal distance between neighbouring peaks: 200 frames, minimum prominence: 6 u, relative peak height to quantify prominence: 0.98).

### In Vivo Methods

#### Electrode implantation

The electrode implantation procedure was adapted from previous studies from Missey^162^ and Acerbo et al^163^. Briefly, animals were first anaesthetised with 3% (vol/vol) isoflurane in oxygen in an induction chamber, and transferred to a nose cone for shaving where isoflurane was kept at 2%. All mice were then injected with carprofen (5 mg/kg), buprenorphine (0.1mg/kg) and dexamethasone (0.01 ml). Petroleum Jelly (Vaseline Original) was applied to the eyes and the hair was removed with a shaving cream (Veet). After shaving, the animal was head-fixed in a stereotaxic frame (Kopf Instruments). The scalp was removed with small scissors, and the skull was cleaned and scraped thoroughly to remove all tissue. The surrounding skin was then glued to the skull with super glue (Loctite). After this, a red activator (Super Bond), containing phosphoric acid, was applied to the exposed skull for 30s and then carefully washed away with saline to ensure adherence of the cement to the skull. The electrode implantation coordinates were marked with ink on the skull (AP −2, ML +0.7, −0.7, −3.1 and −4.5 mm, **Figure S6**), and the skull area was then covered with a thin layer of clear dental cement (Super Bond). Four 0.7 mm holes in the marked locations were drilled through the cement and the skull down to the dura mater (**Figure S6**). Four stainless steel screws (TX000-1.5FH, 0.86 mm diameter, Component Supply), previously attached in pairs to micro-connector sockets, were screwed into the holes (**Figure S6**). The impedance of the connection was measured during this procedure, reaching a value of 0.5 – 1 MΩ DC between all pairs. After implantation, the skull and the screws were secured with dental cement (Super Bond). Mice received carprofen in water for two days post-surgery and were allowed to recover for at least 6 days before the first stimulation session.

#### Hippocampal TI Stimulation

Four days before the start of the TI stimulation protocol, all mice were habituated to the experimenter for 4 days using the cup handling method, in accordance with Hurst and West^178^. The TI stimulation waveforms were generated using MATLAB and delivered via a data acquisition system (DAQ NI USB-6216 BNC, National Instrument) connected to two linear current isolators (Soterix LCI1107 H Precision, Soterix Medical, United States), one per electrode pair. The stimulation was delivered to free-moving animals using custom-made wires and a commutator (**Figure S6**). Two custom-made transformer boxes (Hammond Manufacturing L1140-LN-B) were added in line with the stimulation channels as high-pass filters. The direct impedance of each channel was tested with a multimeter (289 Fluke, United States) before and during stimulation. The carrier frequency was set at 2 kHz, applying f_1_=2 kHz with one electrode pair and f_2_=f_1_+Δf, with the second electrode pair. The three TI stimulation frequencies selected were theta (8 Hz), gamma (40 Hz) and a delta pulsed TI protocol (1 Hz pulses of 100 ms pulse width).

In pulsed TI, two sinusoidal waveforms (f_1_ and f_2_) are applied via two electrode pairs that are out-of-phase, with a phase difference of 180^◦^. Frequency f is fixed, and frequency f_2_ is modulated such that f =f + 1/pulse width during the pulse period, generating a pulse at the given frequency. During the interpulse interval, the two frequencies are equal (f_2_=f_1_), causing destructive interference of the resulting signal. The delta stimulation was applied as a pulsed TI protocol generated by setting both frequencies at 2 kHz during interpulse interval (f_1_=f_2_=2 kHz), and the pulse width at 100 ms (f_2_=f_1_ + 1/(pulse width)=2010 Hz) during the 1 Hz pulse period. This generated a pulsed 1 Hz (delta) stimulation, delivering 10 Hz-bursts. This stimulation pattern was chosen because it has been shown to improve the efficacy of the stimulation^179^ and influence long-term synaptic plasticity^180^. The initial current amplitude was set at a range of 0.3-0.35 mA per channel (initial summed current of 0.7 mA in the short-term study in **Figure 3** and **Figure 4**, and 0.6 mA in the long-term study in **Figure 5**), and reduced when the mice showed signs of sensitisation on an individual basis (current range: 0.2875 ± 0.0875 mA per channel; approximate current density range, dependent on the exact exposed electrode surface: 0.96 ± 0.29 mA/mm^2^). The stimulation was delivered in a 10 s-on-10 s-off fashion with a 0.5 s of linear ramp up and ramp down. The animals were stimulated for 1h per day, for 8 days spread across two weeks. In the short-term study, mice were stimulated for 4 consecutive days, after which they had 2 days of rest and, subsequently, 4 consecutive days of stimulation. In the long-term study, mice had 3 rest days between both 4-day stimulation blocks, instead of 2. Video recordings were collected for each stimulation session from at least 5 min before to 5 min after the stimulation paradigm started, to monitor animal behaviour. The voltage of the two channel pairs was measured before stimulation and every 10 min during stimulation with a multimeter to determine the total impedance values between electrode pairs and ensure stable connection.

#### BrdU Injections

Mice in the long-term *in vivo* study were injected with BrdU (50 mg/kg) intraperitoneally after the 4^th^ and 8^th^ stimulation sessions. BrdU (B5002-500MG, Sigma Aldrich) was previously prepared by diluting it in 0.85% sterile saline (13225649, Thermo Scientific™) and passing the solution through a 0.2 µm filter (725-2520, Thermo Scientific™). The first BrdU injection was delivered immediately after the stimulation session finished. Three additional BrdU injections were delivered every ∼2 h, resulting in a total of 4 injections per mouse and day.

#### Object Pattern Separation Task

The OPS^181^ task is a variation of the object location task (OLT), with the difference of testing several specific object locations that differ by a small distance. For this task, we used a circular acrylic open field that consisted of a 40 cm height and 40 cm diameter cylinder, half of which was opaque grey and half was transparent, and a grey 60 x 60 cm square base. On the base, the possible positions the objects could take were marked with engraved dots arranged forming two vertical lines (**Figure 5G**). The two middle dots, one on each of the two vertical lines, were placed in the middle of the square base, and served as the objects starting positions during the training phase. On the horizontal axis, the middle two dots were placed 22.5 cm from the right and left borders of the square base, respectively, and 15 cm from each other. The remaining engraved dots represented locations to which the objects could be moved to and were equally distributed across the two vertical lines, with half of them being placed above and half of them below each of the middle dots. The vertical spacing between all dots was 3 cm. The open field was placed on a surface 40 cm above the floor and surrounded by a black 4-panel room divider, leaving an opening at the front. Two tall floor lamps, placed at either side of the open field, illuminated the room. Two different sets of identical objects were used, for which we obtained the files to 3D print them in house from Blackmore et al.^182^. The first set of objects consisted of two white 3D printed cuboids with a base of 4 x 4 cm and a height of 10 cm. The second set of objects were two white 3D printed prolate spheroids on a rounded pedestal of 5.8 cm of diameter, and had a total height of 10 cm.

On the first week of the OPS task, mice were habituated for 5 minutes to the circular open field for 2 consecutive days, which they were allowed to explore freely. Subsequently, on OPS day 3, mice were familiarised for 5 minutes to a given set of identical objects, either the first or the second set, and this was randomly allocated and counterbalanced between groups. Leaving a rest day in between, mice were familiarised for 5 minutes to the opposite set of identical objects on OPS day 5. During the object habituation, the objects were placed inside the circular open field and their locations were determined at random within the possible engraved positions. Positions were counterbalanced between both experimental groups. The order in which mice entered the open field was randomised and kept consistent across all sessions.

On the second week of the OPS task, mice performed the task on three different experimental days (OPS days 8, 10 and 12). Each experimental day consisted of two phases, a training phase and a test phase. Mice were allowed to freely explore the open field and objects in each phase for 4 minutes. During the training phase, mice were exposed to either set of identical objects placed at the starting positions (middle engraved dots on the open field). In the test phase, which started 60 minutes after the training phase, one of the identical objects was moved to one of three possible positions (P1=3 cm, P2=6 cm and P3=9 cm, Figure 5G) while the other remained static. The object that was moved (either left or right), the position to which it was moved (P1, P2 or P3), and the direction in which it was moved (backwards or forwards) was randomised across all days. Each mouse performed the test with all three positions.

All sessions were video recorded with a ceiling mounted monochrome industrial camera (DMK 33UX273, The Imaging Source) at a frequency of 30 Hz. Once behavioural sessions were completed, videos were processed offline with DeepLabCut^183^ to label different parts of the mouse’s body, including the nose position, in each videoframe. After this, all data was analysed with a custom-made MATLAB script which first identified the edges of the two objects in the open field, during the training and test phases, and defined two areas to which a border of 1 cm was added. We quantified the time that the mouse’s nose was inside each of these two object areas, with quantification only including the first 20 seconds of combined object exploration time. Mice with a combined object exploration time lower than 7 seconds during the training phase, and/or lower than 10 seconds during the test phase, were excluded from analysis. The time mice spent exploring the static object (T_static object_) and the moved object (T_moved object_) were used to calculate a discrimination index (d2), which was obtained as ‘d2=(T_moved object_ - T_static object_)/ (T_moved object_ + T_static object_)’, and used for analysis.

#### Brain Tissue Processing

For the short-term *in vivo* study, all mice were perfused ∼60 minutes after the last stimulation session and were kept in total darkness during this time. For the long-term *in vivo* study, mice were perfused 33 days after the last stimulation session. Briefly, mice were anaesthetised with 0.03 mL ketamine and 0.01 mL xylazine and perfused with 20 ml of PBS. The brains were then excised and fixed in 4% PFA for 18 – 24 h. After this, the brains were washed in PBS, split in two halves using a standardised matrix, and washed again with PBS. Subsequently, the brains were processed and embedded in paraffin blocks. Coronal brain slices were cut with a microtome (Leica Biosystems) at a thickness of 7 μm, mounted on SuperFrost plus microscope slides (VWR) and allowed to air dry for a minimum of 24 h before staining.

### In Vivo Assays

#### Immunohistochemistry and imaging

One slice per mouse was stained for every biomarker. The biomarkers used were DCX (ab18723, Abcam diluted at 1:4000) for immature cells, Ki67 for active cell proliferation (ab16667, Abcam, diluted at 1:500).

The rehydration procedure consisted of two 5 min washes in xylene (534056-4L, Honeywell), two in 100% absolute ethanol (Eth, 20821.365), one in 90% Eth and one in 70% Eth. Slides were washed for 5 min in deionised water (DI water) and incubated in the dark for 30 min in 0.3% H2O2 (23622.298, VWR) to prevent endogenous peroxidase activity. Antigen retrieval was performed by incubating slices in citrate buffer pH 6.0 (AB93678, Abcam) for 20 min in an electric steamer (Argos) as per Table 5.1. After a 5 min PBS wash, protein blocking was performed for 60 min with 10% normal horse serum (S-2000-20, Vector Laboratories) in PBS (003002, Invitrogen). Slides were then washed twice with PBS and incubated with the primary antibody diluted in 10% NHS/PBS overnight at 4°C. Slides were washed three times in PBS for 5 min and incubated with secondary antibodies ImmPRESS[R] HRP Horse Anti-Rabbit IgG Polymer Detection Kit, Peroxidase (MP-7401-50, Vector Laboratories) for 30 min at room temperature. After three 5 min PBS washes, the staining was visualized with 3,3’-Diaminobenzidine (ImmPACT [R] DAB Substrate, Peroxidase (HRP), SK-4105, Vector Laboratories). Slides were washed in DI water for 5 min and incubated in hematoxylin (MHS32-1L, Mayer’s) for 2 to 3 minutes. Lastly, the slides were washed in tap water and dehydrated with three 5min washes of 70% Eth, 90% Eth, 100% Eth and three washes of xylene. The slides were mounted with DPX mounting medium (D/5330/05, Fisher Scientific) and imaged at 20x magnification using a bright-field scanner (Leica Aperio AT2), obtaining a pixel dimension of 0.5040 μm.

#### Immunofluorescence staining and imaging

Primary antibodies used for co-staining were rabbit anti-DCX (1:200, ab18723, Abcam) for immature neurons, mouse anti-NeuN (1:4000, Ab104224, Abcam) for mature neurons, and rat anti-BrdU (1:250, Ab6326, Abcam). Secondary antibodies used include Alexa Fluor™ 488 Goat anti-Rabbit IgG (H+L) (1:200, A-11034, Invitrogen), Alexa Fluor™ 647 Goat anti-Mouse IgG (H+L) (1:200, A-21236, Invitrogen) and Alexa Fluor™ 555 Goat anti-Rat IgG (H+L) (1:750, A-21434, Invitrogen). Two slices per mouse were stained to capture more BrdU+ cells for statistical quantification. Two slices were at least 0.1mm away from each other along the AP axis to avoid quantifying the same cell twice.

Rehydration steps were identical to the immunohistochemistry staining. After washing the slides in DI water for 5 min, antigen retrieval and BrdU exposure were performed simultaneously by incubating the slides in citrate buffer pH 6.0 (Ab93678, Abcam) for 40 min in the electrical steamer (Argos). Slides were subsequently cooled down in an ice water bath for 15 min. After two 5 min washes with PBS (003002, Invitrogen), slices were blocked with 10% Normal Goat Serum (Ab7481, Abcam) in PBS for one h at room temperature. Slices were then incubated with a primary antibody diluent of 5% Normal Goat Serum and 0.1% Triton-X100 (93443, Sigma-Aldrich) in PBS at 4°C overnight. The next day, slides were washed with PBS for 5 min three times. They were next incubated with a secondary antibody diluent, which had the same composition as the primary diluent, for 1 h at room temperature in the dark. After three 5 min PBS washes, the slices were incubated with 1:1000 DAPI (62248, ThermoFisher) in PBS. Following two 5 min PBS washes, slices were mounted with Anti-Fade Fluorescence Mounting Medium (Ab104135, Abcam) and kept in a dark and damp environment at 4°C. Imaging was performed within 2 weeks post-staining.

Slices were imaged with Leica SP8 Lightning Confocal Microscope and Leica Application Suite X (v3.5.7) with a z-stack tiled scan. The Regions of Interest (ROI) were outlined to cover the whole dentate gyrus, including the molecular layers. Depending on the sample, fifteen to twenty tiles were acquired to cover the ROI. Both left and right dentate gyri were imaged and analysed. Scans were 1024×1024 pixels, with a pixel size of 284.09nm x 284.09nm, taken with an HC PL APO CS2 40x/1.30 Oil objective (Leica) in 16-bit mode. Images had optical sections of approximately 1 μm and a pixel dwell time of approximately 1.5 μs. Fifteen z-stacks with z-steps of 1.3-2.0 μm were imaged for each sample. Tiles were later merged with the Suite’s mosaic merge function. Images were then z-projected with maximum intensity using ImageJ software. Acquired images were later analysed with MATLAB.

#### Proteomics sample preparation

Mass spectrometry-based proteomics was performed on the Formalin-fixed paraffin-embedded (FFPE) tissues. The protocol was adapted from the method reported by Coscia et al ^183^. Slices were rehydrated using the same steps in immunohistochemistry and immunofluorescence. To visualise cells and structures, samples were treated with hematoxylin (MHS32-1L, Mayer’s) for 2 min. Bluing was performed by washing the slices with running tap water for 5 min. The slides were submerged for 5 min each in 70% Eth, 90% Eth and two 100% Eth baths for dehydration. Afterwards, the slices were microdissected under a stereo microscope (Olympus SZ-PT). Each dentate gyrus was outlined with a knife scalpel (221-4438, RS Components), and 0.3μL of Milli-Q water (Merck Milli-Q) was added to the ROI for tissue collection with a pipette. Each sample was then added to 100μL of freshly prepared lysis buffer that consisted of 50% v/v 2,2,2-Trifluoroethanol (SIAL96924, Sigma-Aldrich), 300mM Tris-hydrochloride (10812846001, Roche) and Milli-Q water. The tissues in the buffer were snap-frozen with dry ice and kept at −80 °C until sent to the collaborator for analysis.

#### Protein digestion

10ug of protein samples were digested with S-trap micro columns (ProtiFi) following the recommended manufacturer’s protocols. Samples were alkylated with Ttris(2-carboxyethyl)phosphine (TCEP) at 56° for 30 minutes before alkylation for 30 minutes using Iodoacetamide in the dark. Enzymatic digestion was performed using Trypsin / Lys-C mix (Pierce) using an enzyme to protein ratio of 1:20. Digested peptides were eluted from the S-trap columns and were lyophilized using a vacuum concentrator (Thermo) before storage at −20° before mass spectrometry analysis.

#### Mass spectrometry

Digested peptide samples were resuspended in 0.1 % formic acid prior to preparation on EvoTips (EvoSep, Odense, Denmark). Following the manufacturer’s instructions, EvoTips were first washed with 100% Acetonitrile 0.1% formic acid. Tips were then conditioned using 1-propanol prior to an equilibration step using 0.1% formic acid. Loading was performed by addition of 100ng of sample to the tip prior to washing that was repeated 3 times using 0.1% formic acid. 100uL of 0.1% formic acid was left in the tip to avoid drying out. Prepared tips were then injected onto the MS using an EvoSep One LC system, using the 30 samples per day method. The analytical column used was an “endurance” column (C18, 1.9 µm, 15 cm x 150 µm). Samples were acquired on a Bruker timsTOF SCP in DIA PASEF mode using the standard method settings (**Figure S7**).

#### Data searching and analysis

Mass spectra files were then searched using DIA-NN v.1.9 with the library free method enabled. The protein database was a human FASTA file (Uniprot, downloaded June 2024). The default parameters were enabled with the “Heuristic protein inference” switched off and the “double pass” classifier being used instead of the single pass. Downstream normalization and imputation were performed in R (version 4.1.3). The Enrichr website^175^ was used to implement the GO enrichment analysis of biological process (2023), with a cut-off of non-adjusted p<0.05 and log_2_(Fold Change)>0.

### Image Analysis

#### Biomarker Density and Cell Count

The IHC bright-field images were analysed using a custom MATLAB pipeline. Images were deconvolved into separate Haematoxylin and DAB channels using built-in ‘Colour deconvolution’ function in Fiji software^184^. Two regions of interest (ROIs) corresponding to the left and right DG of the hippocampus were selected on both hemispheres in a semi-automated manner. Briefly, the centre of each DG was manually selected, and a standardised ROI window of size 1300 μm by 500 μm was centred around each selected point to fit the whole DG. Large artefacts such as tissue folds were then removed by manually drawing an exclusion area per image. Areas containing no tissue were also excluded. A slice mask containing only the included tissue was then generated for each image. After this, the brightness of the image was corrected by adjusting its histogram, with the ‘imhistmatch’ function, such that the pixel intensity matches the histogram of a common reference image for the corresponding antibody^185^. The function ‘imfilter’ was then used to apply a box filter to all images (kernel size: 3×3 pixels) for spatial smoothing and noise reduction^186^. The pixel values were then binarized according to a pre-selected threshold value specific to each biomarker. For Ki67 staining, a watershed algorithm was applied to separate connected cells^187^. Lastly, all particles in the mask were excluded if outside the pre-selected size boundaries (3-30 μm for DCX and Ki67; 4-50 μm GFAP and Iba1). The area covered by each antibody (μm^2^) was normalised by the total ROI area (mm^2^) and expressed as μm^2^/mm^2^. The total cell number in each ROI was quantified by identifying individual objects in the antibody binary mask, using the function ‘regionprops’, and normalised by the total ROI area (mm^2^) to obtain cell density.

#### Quantification of Cellular Maturation Stage

A MATLAB pipeline was developed to quantify the maturation stage of DCX+ cells based on morphology in a semi-manual manner. First, IHC or IF images of each pre-selected DG ROI were loaded into the MATLAB workspace. All DCX+ cells were then selected manually and assigned to a maturation category as described by Plümpe et al^78^ (**Figure 4A**), with some modifications. Due to the thinness of the brain slices used (7 µm), continuous cell processes may appear as discontinuous segments. During cell classification, segments were considered part of the same process if they followed a consistent trajectory and the gap between them was smaller than the average cell nucleus size. DCX+ cells with no processes or processes shorter than one cell body were assigned to the proliferative stage. DCX+ cells with a process longer than one cell body that does not project into the molecular layer were assigned to the intermediate stage. Finally, DCX+ cells with a long process projecting into the molecular layer, with or without branches, were assigned to the postmitotic stage. After cell classification was finalised, a variable number of points were selected by the user to systematically trace the inner and outer border of the GCL of the DG (Figure 4B). These outlines also allowed to distinguish between dGCL and vGCL. The points used to trace the inner part of the GCL were placed with a gap, consisting of the space equivalent to two cell nuclei, in order to also include the SGZ. The number of cells per category was extracted for the dGCL and vGCL of both the contralateral and ipsilateral ROIs. Cell number in each category was normalised to the corresponding total, dGCL or vGCL area (mm^2^) to obtain the cell density. Three different experimenters analysed all images independently, following the same guidelines, and their results were averaged to minimise experimenter bias. For the short-term *in vivo* study, only one brain slice per animal was quantified. However, two brain slices per animal were quantified in the long-term *in vivo* study, as these were available due to the low number of BrdU+ cells observed, and results were averaged.

#### Quantification of BrdU+ Cells

The number of BrdU+ cells was semi-manually quantified using a custom-made MATLAB pipeline. First, IF images of each pre-selected DG ROI were loaded into the MATLAB workspace. All BrdU+ cells were then selected manually and the presence of NeuN and/or DCX co-expression was visually assessed (Figure 5E). Cells with BrdU partial incorporation were also quantified as BrdU+ cells. Subsequently, a number of points were selected by the user to systematically trace the inner and outer border of the GCL of the DG and the boundary between the hilus and CA3 (**Figure 5B, Figure S4**). These outlines also allowed to distinguish between dGCL and vGCL. The points used to trace the inner layer of the GCL were placed with a gap consisting of the space equivalent to two cell nuclei, in order to also include the SGZ. The markers DCX and NeuN were used to distinguish between two categories: mature GCs (BrdU+/DCX-/NeuN+) and cells with other non-neuronal fate (BrdU+/DCX-/NeuN-). The number of BrdU+ cells per category was extracted for the dGCL and vGCL and the hilus of both the contralateral and ipsilateral ROIs. Cell number in each category was normalised to the corresponding GCL area or hilus area (mm^2^) to obtain the cell density. Three different experimenters analysed all images independently, following the same guidelines, and their results were averaged to minimise experimenter bias. Two brain slices per animal were quantified, due to the low number of BrdU+ cells observed, and results were averaged.

### Statistical analysis

#### In vitro analyses

All data represent the mean of at least 3 biological repeats (n≥3) derived from 3 animals, each of them consisting of at least 2 technical repeats (6-9 tissue culture wells per condition). To account for the variability between animals, a linear mixed effect model was implemented^188^, where the stimulation treatments were considered fixed variables and the sham control was used as a reference. The biological repeat was modelled as a random intercept, and technical repeats were nested as withing biological repeats and also modeled as a random intercept. We used the “fitlme” function, applying the maximum likelihood estimation (ML) method, in MATLAB (Version R2023b, Statistics and Machine Learning Toolbox). Model residuals were examined for normality and homoscedasticity (via Q-Q plots and residual distribution plots). All results are expressed as the model estimated means ± the 95% CI. The data points represent the percentual change compared to the average of the sham. The mixed model predictions for the fixed and random variable were considered significant at a significance level of 5% (^∗^p<0.05, ^∗∗^p<0.01, ^∗∗∗^p<0.001). The F statistics and degrees of freedom were calculated with an ANOVA, Satterthwaite method^189^. To evaluate the significance of the biological repeat effect, we performed a likelihood ratio test^190^ by comparing the full model to a reduced model excluding the biological repeat (random effect, where c^2^=2 × (log-likelihood of the full model – log-likelihood of the model without random effect). This test determined whether inclusion of the biological effect significantly improved model fit. In case of multiple comparisons, the results were corrected with Bonferroni-Holm (BH) post-hoc correction^191^. We identified outliers using a threshold of z-score >3 on the model residuals. If outliers were detected, we ran the analysis both with and without outliers. Here, we report the most conservative results. The results of all statistical tests are reported in **Table S1, S2, S3**.

#### *In vivo* analyses

The short-term study dataset was assessed for normality with the Lilliefors test^192^. If the data were found to be normally distributed (p>0.05), an independent t-test was used, to compare the means of each stimulation group to the sham group, for a total of four comparisons^193^. Otherwise, a non-parametric Wilcoxon rank-sum test was employed in the same way. The results were corrected for multiple comparisons with Bonferroni-Holm (BH) correction^191^

All results obtained with the manual quantification of DCX+ cells (**Figures 4**, **Figure 5**), as well as all the OPS task results (**Figure 5H**), were analysed with repeated measures ANOVA using the function ‘aov_car’ from the ‘afex’ package in R. We identified outliers with the function ‘check_outliers’ from the ‘performance’ package in R, using a threshold of 3 with the Z-score method. If outliers were detected, we ran the analysis both with and without outliers. Here we report the most conservative results from both analyses. We assessed the normality of the residuals with the Shapiro-Wilk test. Although we observed some rare normality violations, these were mostly present in the analysis performed including outliers and usually resolved in the analysis without outliers. In addition to this, the repeated measures ANOVA test has been shown to be robust against this kind of violation^194^. All results obtained with the manual quantification of BrdU+ cells (Figure 5F) were analysed with two-sided unpaired two-samples Wilcoxon rank-sum tests in R due to the presence of outliers.

Differences were considered significant at a significance level of 5% (^∗^p<0.05, ^∗∗^p<0.01, ^∗∗∗^p<0.001). The results of all statistical tests are reported in **Table S4, S5, S6**.

## 6. RESOURCE AVAILABILITY

### Lead contact

Further information and requests for resources and reagents should be directed to and will be fulfilled by the lead contact, Sofia Peressotti.

### Materials availability

This study did not generate new unique reagents.

### Data and code availability

Any additional information is available from the lead contact upon request.

## 7. ACKNOWLEDGMENTS

The authors thank Dr. Josef Goding (Imperial College London, UK), Prof. Sandrine Thuret (King’s College London, UK), Dr. Robert Toth, Dr. Estelle Cuttaz, Dr. Erol Hassan, Dr. Kjara Pilch for expert technical assistance, Gillian Kohel for the support in the biomaterial synthesis and gelling. We acknowledge the support of the Facility for Imaging by Light Microscopy (FILM) at the faculty of Medicine, Imperial College London. We thank the technicians and staff of the central biomedical services (CBS) at Imperial College London, Hammersmith Campus, and the UK dementia research institute (DRI) for the resources provided for the proteomics analysis.

The authors acknowledge funding support from: the European Research Council through the Consolidator Grant 771985, the UK Dementia Research Institute, Engineering and Physical Sciences Research Council, UK, EP/W004844/1 (NG), National Institute for Health and Care Research, Imperial Biomedical Research Centre, the Imperial College Bioengineering Department (S.P.), the EPSRC PhD scholarship via Center for Doctoral Training in Neurotechnology at Imperial College London (P.D.).

Figures and graphical abstract were partly created using Biorender (https://BioRender.com).

## 8. AUTHOR CONTRIBUTIONS

S.P. led the work, conceived and performed the experiments for the in vitro sections, and wrote the first draft of the manuscript. S.P. and R.P.L. conceptualized the work. S.P. and M.G.G. conceived, performed the experiments and wrote the *in vivo* sections of the manuscript. P.D., conceived and performed the *in vivo* experiments and related analyses. R.M.L., B.G., E.F., M.W., M.O.J., M.G., L.T. performed experiments or analyses. K.D., J.A.A. provided materials and expertise. M.G.G., N.G. and R.P.L. edited the manuscript. All authors reviewed the manuscript. N.G. and R.G. provided funding and supervision.

## 10. SUPPLEMENTAL INFORMATION

Refer to Document S1. Figures S1-S7 and Tables S1-S6.

## REFERENCES

1. Hansson, O. Biomarkers for neurodegenerative diseases. Nat Med 27, 954–963 (2021).

2. Feigin, V. L. et al. The global burden of neurological disorders: translating evidence into policy. Lancet Neurol 19, 255–265 (2020).

3. Hou, Y. et al. Ageing as a risk factor for neurodegenerative disease. Nat Rev Neurol 15, 565–581 (2019).

4. Livingston, G. et al. Dementia prevention, intervention, and care: 2024 report of the Lancet standing Commission. The Lancet 404, 572–628 (2024).

5. Bloem, B. R., Okun, M. S. & Klein, C. Parkinson’s disease. The Lancet 397, 2284–2303 (2021).

6. Temple, S. Advancing cell therapy for neurodegenerative diseases. Cell Stem Cell 30, 512–529 (2023).

7. Varadarajan, S. G., Hunyara, J. L., Hamilton, N. R., Kolodkin, A. L. & Huberman, A. D. Central nervous system regeneration. Cell 185, 77–94 (2022).

8. Yamanaka, S. Pluripotent Stem Cell-Based Cell Therapy—Promise and Challenges. Cell Stem Cell 27, 523–531 (2020).

9. Navarro Negredo, P., Yeo, R. W. & Brunet, A. Aging and Rejuvenation of Neural Stem Cells and Their Niches. Cell Stem Cell 27, 202–223 (2020).

10. Sun, M. K. & Alkon, D. L. Treating Alzheimer’s Disease: Focusing on Neurodegenerative Consequences. Journal of Alzheimer’s Disease 101, S263–S274 (2024).

11. Tabar, V. et al. Phase I trial of hES cell-derived dopaminergic neurons for Parkinson’s disease. Nature 2025 641:8064 641, 978–983 (2025).

12. Sivandzade, F. & Cucullo, L. Regenerative Stem Cell Therapy for Neurodegenerative Diseases: An Overview. Int J Mol Sci 22, 2153 (2021).

13. Babcock, K. R., Page, J. S., Fallon, J. R. & Webb, A. E. Adult Hippocampal Neurogenesis in Aging and Alzheimer’s Disease. Stem Cell Reports 16, 681–693 (2021).

14. Kempermann, G. et al. Human Adult Neurogenesis: Evidence and Remaining Questions. Cell Stem Cell 23, 25–30 (2018).

15. Anacker, C. & Hen, R. Adult hippocampal neurogenesis and cognitive flexibility — linking memory and mood. Nature Reviews Neuroscience 2017 18:6 18, 335–346 (2017).

16. Baptista, P. & Andrade, J. P. Adult hippocampal neurogenesis: Regulation and possible functional and clinical correlates. Front Neuroanat 12, 365041 (2018).

17. Toda, T. & Gage, F. H. Review: adult neurogenesis contributes to hippocampal plasticity. Cell and Tissue Research 2017 373:3 373, 693–709 (2017).

18. Toda, T., Parylak, S. L., Linker, S. B. & Gage, F. H. The role of adult hippocampal neurogenesis in brain health and disease. Mol Psychiatry 24, 67–87 (2019).

19. Kempermann, G., Song, H. & Gage, F. H. Neurogenesis in the Adult Hippocampus. Cold Spring Harb Perspect Biol 7, a018812 (2015).

20. Gonçalves, J. T., Schafer, S. T. & Gage, F. H. Adult Neurogenesis in the Hippocampus: From Stem Cells to Behavior. Cell 167, 897–914 (2016).

21. Dubois, B. et al. Clinical diagnosis of Alzheimer’s disease: recommendations of the International Working Group. The Lancet Neurology Preprint at 10.1016/S1474-4422(21)00066-1 (2021).

22. Moreno-Jiménez, E. P. et al. Adult hippocampal neurogenesis is abundant in neurologically healthy subjects and drops sharply in patients with Alzheimer’s disease. Nature Medicine 2019 25:4 25, 554–560 (2019).

23. Tobin, M. K. et al. Human Hippocampal Neurogenesis Persists in Aged Adults and Alzheimer’s Disease Patients. Cell Stem Cell 24, 974–982.e3 (2019).

24. Skidmore, S. & Barker, R. A. Challenges in the clinical advancement of cell therapies for Parkinson’s disease. Nature Biomedical Engineering 2023 7:4 7, 370–386 (2023).

25. Barker, R. A. et al. Designing stem-cell-based dopamine cell replacement trials for Parkinson’s disease. Nature Medicine 2019 25:7 25, 1045–1053 (2019).

26. Parmar, M., Grealish, S. & Henchcliffe, C. The future of stem cell therapies for Parkinson disease. Nature Reviews Neuroscience 2020 21:2 21, 103–115 (2020).

27. Iwasa, S. N. et al. Electrical Stimulation for Stem Cell-Based Neural Repair: Zapping the Field to Action. eNeuro 11, (2024).

28. Balasubramanian, S., Weston, D. A., Levin, M. & Davidian, D. C. C. Charging Ahead: Examining the Future Therapeutic Potential of Electroceuticals. Adv Ther (Weinh) 7, 2300344 (2024).

29. Zhu, R. et al. Electrical stimulation affects neural stem cell fate and function in vitro. Exp Neurol 319, 112963 (2019).

30. Cheng, H., Huang, Y., Yue, H. & Fan, Y. Electrical Stimulation Promotes Stem Cell Neural Differentiation in Tissue Engineering. Stem Cells Int 2021, (2021).

31. Thompson, D. M., Koppes, A. N., Hardy, J. G. & Schmidt, C. E. Electrical Stimuli in the Central Nervous System Microenvironment. Annu Rev Biomed Eng 16, 397–430 (2014).

32. Wu, Z. et al. Impact of subthalamic nucleus deep brain stimulation at different frequencies on neurogenesis in a rat model of Parkinson’s disease. Heliyon 10, (2024).

33. Chamaa, F. et al. Long-term stimulation of the anteromedial thalamus increases hippocampal neurogenesis and spatial reference memory in adult rats. Behavioural Brain Research 402, (2021).

34. Chamaa, F., Sweidan, W., Nahas, Z., Saade, N. & Abou-Kheir, W. Thalamic Stimulation in Awake Rats Induces Neurogenesis in the Hippocampal Formation. Brain Stimul 9, 101–108 (2016).

35. Toda, H., Hamani, C., Fawcett, A. P., Hutchison, W. D. & Lozano, A. M. The regulation of adult rodent hippocampal neurogenesis by deep brain stimulation: Laboratory investigation. J Neurosurg 108, 132–138 (2008).

36. Hao, S. et al. Forniceal deep brain stimulation rescues hippocampal memory in Rett syndrome mice. Nature 2015 526:7573 526, 430–434 (2015).

37. Wang, Q. et al. Forniceal deep brain stimulation in a mouse model of Rett syndrome increases neurogenesis and hippocampal memory beyond the treatment period. Brain Stimul 16, 1401–1411 (2023).

38. Stone, S. S. D. et al. Stimulation of entorhinal cortex promotes adult neurogenesis and facilitates spatial memory. Journal of Neuroscience 31, 13469–13484 (2011).

39. Sun, Y. W. et al. Multiple Sessions of Entorhinal Cortex Deep Brain Stimulation in C57BL/6J Mice Increases Exploratory Behavior and Hippocampal Neurogenesis. *Proceedings of the Annual International Conference of the IEEE Engineering in Medicine and Biology Society*, EMBS 2021-January, 6390–6393 (2021).

40. Lozano, A. M. et al. Deep brain stimulation: current challenges and future directions. Nature Reviews Neurology 2019 15:3 15, 148–160 (2019).

41. Dumontoy, S. et al. Repeated Anodal Transcranial Direct Current Stimulation (RA-tDCS) over the Left Frontal Lobe Increases Bilateral Hippocampal Cell Proliferation in Young Adult but Not Middle-Aged Female Mice. Int J Mol Sci 24, (2023).

42. Yu, T. H., Wu, Y. J., Chien, M. E. & Hsu, K. Sen. Multisession Anodal Transcranial Direct Current Stimulation Enhances Adult Hippocampal Neurogenesis and Context Discrimination in Mice. Journal of Neuroscience 43, 635–646 (2023).

43. Liu, Q. et al. Intracranial alternating current stimulation facilitates neurogenesis in a mouse model of Alzheimer’s disease. Alzheimers Res Ther 12, 1–12 (2020).

44. Cuccurazzu, B. et al. Exposure to extremely low-frequency (50 Hz) electromagnetic fields enhances adult hippocampal neurogenesis in C57BL/6 mice. Exp Neurol 226, 173–182 (2010).

45. Deng, Z. De, Lisanby, S. H. & Peterchev, A. V. Electric field depth-focality tradeoff in transcranial magnetic stimulation: Simulation comparison of 50 coil designs. Brain Stimul (2013) doi:10.1016/j.brs.2012.02.005.

46. Grossman, N. et al. Noninvasive Deep Brain Stimulation via Temporally Interfering Electric Fields. Cell 169, 1029–1041.e16 (2017).

47. Violante, I. R. et al. Non-invasive temporal interference electrical stimulation of the human hippocampus. Nature Neuroscience 2023 26:11 26, 1994–2004 (2023).

48. Yang, J. W., Hanganu-Opatz, I. L., Sun, J. J. & Luhmann, H. J. Three Patterns of Oscillatory Activity Differentially Synchronize Developing Neocortical Networks In Vivo. Journal of Neuroscience 29, 9011–9025 (2009).

49. Luhmann, H. J. et al. Spontaneous neuronal activity in developing neocortical networks: From single cells to large-scale interactions. Front Neural Circuits 10, 197897 (2016).

50. Graham, V., Khudyakov, J., Ellis, P. & Pevny, L. SOX2 Functions to Maintain Neural Progenitor Identity.Neuron 39, 749–765 (2003).

51. Pagin, M. et al. Sox2 controls neural stem cell self-renewal through a Fos-centered gene regulatory network. Stem Cells 39, 1107–1119 (2021).

52. Visan, A. et al. Neural differentiation of mouse embryonic stem cells as a tool to assess developmental neurotoxicity in vitro. Neurotoxicology 33, 1135–1146 (2012).

53. Baldwin, K. T., Murai, K. K. & Khakh, B. S. Astrocyte morphology. Trends Cell Biol 34, 547–565 (2024).

54. Murphy-Royal, C., Ching, S. N. & Papouin, T. A conceptual framework for astrocyte function. Nat Neurosci 26, 1848–1856 (2023).

55. McNeill, J., Rudyk, C., Hildebrand, M. E. & Salmaso, N. Ion Channels and Electrophysiological Properties of Astrocytes: Implications for Emergent Stimulation Technologies. Front Cell Neurosci 15, 644126 (2021).

56. Hansen, M. G., Tornero, D., Canals, I., Ahlenius, H. & Kokaia, Z. In vitro functional characterization of human neurons and astrocytes using calcium imaging and electrophysiology. in Methods in Molecular Biology vol. 1919 73–88 (Humana Press Inc., 2019).

57. de Groot, M. W. G. D. M., Dingemans, M. M. L., Rus, K. H., de Groot, A. & Westerink, R. H. S. Characterization of Calcium Responses and Electrical Activity in Differentiating Mouse Neural Progenitor Cells In Vitro. Toxicological Sciences 137, 428–435 (2014).

58. Sharma, Y., Saha, S., Joseph, A., Krishnan, H. & Raghu, P. In vitro human stem cell derived cultures to monitor calcium signaling in neuronal development and function. Wellcome Open Res 5, (2020).

59. Hayashi, H., Edin, F., Li, H., Liu, W. & Rask-Andersen, H. The effect of pulsed electric fields on the electrotactic migration of human neural progenitor cells through the involvement of intracellular calcium signaling. Brain Res 1652, 195–203 (2016).

60. Forostyak, O., Romanyuk, N., Verkhratsky, A., Sykova, E. & Dayanithi, G. Plasticity of calcium signaling cascades in human embryonic stem cell-derived neural precursors. Stem Cells Dev 22, 1506–1521 (2013).

61. Liu, Q. & Song, B. Electric field regulated signaling pathways. Int J Biochem Cell Biol 55, 264–268 (2014).

62. Cui, M. et al. Electromagnetic Fields for the Regulation of Neural Stem Cells. Stem Cells Int 2017, (2017).

63. Wolf-Goldberg, T., Barbul, A., Ben-Dov, N. & Korenstein, R. Low electric fields induce ligand-independent activation of EGF receptor and ERK via electrochemical elevation of H+ and ROS concentrations. Biochimica et Biophysica Acta (BBA) - Molecular Cell Research 1833, 1396–1408 (2013).

64. Khan, J. et al. Neurosphere Development from Hippocampal and Cortical Embryonic Mixed Primary Neuron Culture: A Potential Platform for Screening Neurochemical Modulator. ACS Chem Neurosci 9, 2870–2878 (2018).

65. Roth, J. G. et al. Advancing models of neural development with biomaterials. Nature Reviews Neuroscience 2021 22:10 22, 593–615 (2021).

66. Simão, D. et al. Recapitulation of Human Neural Microenvironment Signatures in iPSC-Derived NPC 3D Differentiation. Stem Cell Reports 11, 552–564 (2018).

67. Chandrasekaran, A. et al. Comparison of 2D and 3D neural induction methods for the generation of neural progenitor cells from human induced pluripotent stem cells. Stem Cell Res 25, 139–151 (2017).

68. Balasubramanian, S., Packard, J. A., Leach, J. B. & Powell, E. M. Three-Dimensional Environment Sustains Morphological Heterogeneity and Promotes Phenotypic Progression During Astrocyte Development. https://home.liebertpub.com/tea 22, 885–898 (2016).

69. Freitag, K. et al. Diverse but unique astrocytic phenotypes during embryonic stem cell differentiation, culturing and development. Communications Biology 2023 6:1 6, 1–12 (2023).

70. Galland, F. et al. Astrocyte culture models: Molecular and function characterization of primary culture, immortalized astrocytes and C6 glioma cells. Neurochem Int 131, (2019).

71. Toda, T., Parylak, S. L., Linker, S. B. & Gage, F. H. The role of adult hippocampal neurogenesis in brain health and disease. Molecular Psychiatry 2018 24:1 24, 67–87 (2018).

72. Colgin, L. L. Rhythms of the hippocampal network. Nature Reviews Neuroscience 2016 17:4 17, 239–249 (2016).

73. Saito, T. et al. Single App knock-in mouse models of Alzheimer’s disease. Nature Neuroscience 2014 17:5 17, 661–663 (2014).

74. Kee, N., Sivalingam, S., Boonstra, R. & Wojtowicz, J. M. The utility of Ki-67 and BrdU as proliferative markers of adult neurogenesis. J Neurosci Methods 115, 97–105 (2002).

75. Couillard-Despres, S. et al. Doublecortin expression levels in adult brain reflect neurogenesis. European Journal of Neuroscience 21, 1–14 (2005).

76. Iaccarino, H. F. et al. Gamma frequency entrainment attenuates amyloid load and modifies microglia. Nature 540, 230–235 (2016).

77. Sasaki, Y., Ohsawa, K., Kanazawa, H., Kohsaka, S. & Imai, Y. Iba1 Is an Actin-Cross-Linking Protein in Macrophages/Microglia. Biochem Biophys Res Commun 286, 292–297 (2001).

78. Plümpe, T. et al. Variability of doublecortin-associated dendrite maturation in adult hippocampal neurogenesis is independent of the regulation of precursor cell proliferation. BMC Neurosci 7, 1–14 (2006).

79. Taupin, P. BrdU immunohistochemistry for studying adult neurogenesis: Paradigms, pitfalls, limitations, and validation. Brain Res Rev 53, 198–214 (2007).

80. Mahmoud, R., Wainwright, S. R. & Galea, L. A. M. Sex hormones and adult hippocampal neurogenesis: Regulation, implications, and potential mechanisms. Front Neuroendocrinol 41, 129–152 (2016).

81. Walgrave, H. et al. Restoring miR-132 expression rescues adult hippocampal neurogenesis and memory deficits in Alzheimer’s disease. Cell Stem Cell 28, 1805–1821.e8 (2021).

82. van Goethem, N. P., van Hagen, B. T. J. & Prickaerts, J. Assessing spatial pattern separation in rodents using the object pattern separation task. Nature Protocols 2018 13:8 13, 1763–1792 (2018).

83. Pofahl, M. et al. Synchronous activity patterns in the dentate gyrus during immobility. Elife 10, (2021).

84. Ge, S., Yang, C. hao, Hsu, K. sen, Ming, G. li & Song, H. A Critical Period for Enhanced Synaptic Plasticity in Newly Generated Neurons of the Adult Brain. Neuron 54, 559–566 (2007).

85. Snyder, J. S. Recalibrating the Relevance of Adult Neurogenesis. Trends Neurosci 42, 164–178 (2019).

86. Chen, P., Chen, F., Wu, Y. & Zhou, B. New Insights Into the Role of Aberrant Hippocampal Neurogenesis in Epilepsy. Front Neurol 12, 727065 (2021).

87. Hagihara, H. et al. Expression of progenitor cell/immature neuron markers does not present definitive evidence for adult neurogenesis. Mol Brain 12, 1–6 (2019).

88. Foltran, R. B., Stefani, K. M., Bonafina, A., Resasco, A. & Diaz, S. L. Differential Hippocampal Expression of BDNF Isoforms and Their Receptors Under Diverse Configurations of the Serotonergic System in a Mice Model of Increased Neuronal Survival. Front Cell Neurosci 13, 466190 (2019).

89. Cellier, D., Riddle, J., Petersen, I. & Hwang, K. The development of theta and alpha neural oscillations from ages 3 to 24 years. Dev Cogn Neurosci 50, (2021).

90. Luhmann, H. J. & Khazipov, R. Neuronal activity patterns in the developing barrel cortex. Neuroscience 368, 256–267 (2018).

91. Martini, F. J., Guillamón-Vivancos, T., Moreno-Juan, V., Valdeolmillos, M. & López-Bendito, G. Spontaneous activity in developing thalamic and cortical sensory networks. Neuron 109, 2519–2534 (2021).

92. Kilb, W., Kirischuk, S. & Luhmann, H. J. Electrical activity patterns and the functional maturation of the neocortex. European Journal of Neuroscience 34, 1677–1686 (2011).

93. Luhmann, H. J. et al. Spontaneous neuronal activity in developing neocortical networks: From single cells to large-scale interactions. Front Neural Circuits 10, 197897 (2016).

94. An, S., Kilb, W. & Luhmann, H. J. Sensory-evoked and spontaneous gamma and spindle bursts in neonatal rat motor cortex. Journal of Neuroscience 34, 10870–10883 (2014).

95. Lindemann, C., Ahlbeck, J., Bitzenhofer, S. H. & Hanganu-Opatz, I. L. Spindle Activity Orchestrates Plasticity during Development and Sleep. Neural Plast 2016, (2016).

96. Sharf, T. et al. Functional neuronal circuitry and oscillatory dynamics in human brain organoids. Nature Communications 2022 13:1 13, 1–20 (2022).

97. Chang, H. F., Lee, Y. S., Tang, T. K. & Cheng, J. Y. Pulsed DC Electric Field–Induced Differentiation of Cortical Neural Precursor Cells. PLoS One 11, e0158133 (2016).

98. Sordini, L. et al. Effect of Electrical Stimulation Conditions on Neural Stem Cells Differentiation on Cross-Linked PEDOT:PSS Films. Front Bioeng Biotechnol 9, 591838 (2021).

99. Thompson, D. M., Koppes, A. N., Hardy, J. G. & Schmidt, C. E. Electrical Stimuli in the Central Nervous System Microenvironment. 10.1146/annurev-bioeng-121813-120655 16, 397–430 (2014).

100. Huang, Y., Li, Y., Chen, J., Zhou, H. & Tan, S. Electrical stimulation elicits neural stem cells activation: New perspectives in CNS repair. Front Hum Neurosci 9, 156639 (2015).

101. Zhou, H. et al. The antidepressant effect of nucleus accumbens deep brain stimulation is mediated by parvalbumin-positive interneurons in the dorsal dentate gyrus. Neurobiol Stress 21, 100492 (2022).

102. Bambico, F. R. et al. Neuroplasticity-dependent and -independent mechanisms of chronic deep brain stimulation in stressed rats. Translational Psychiatry 2015 5:11 5, e674–e674 (2015).

103. Hao, S. et al. Forniceal deep brain stimulation rescues hippocampal memory in Rett syndrome mice. Nature 2015 526:7573 526, 430–434 (2015).

104. Encinas, J. M., Hamani, C., Lozano, A. M. & Enikolopov, G. Neurogenic hippocampal targets of deep brain stimulation. Journal of Comparative Neurology 519, 6–20 (2011).

105. Ramírez-Rodríguez, G. B., Juan, D. M. S. & González-Olvera, J. J. 5□Hz of repetitive transcranial magnetic stimulation improves cognition and induces modifications in hippocampal neurogenesis in adult female Swiss Webster mice. Brain Res Bull 186, 91–105 (2022).

106. Hescham, S. et al. Fornix deep brain stimulation induced long-term spatial memory independent of hippocampal neurogenesis. Brain Struct Funct 222, 1069–1075 (2017).

107. Stone, S. S. D. et al. Stimulation of Entorhinal Cortex Promotes Adult Neurogenesis and Facilitates Spatial Memory. Journal of Neuroscience 31, 13469–13484 (2011).

108. Lim, J. H., McCullen, S. D., Piedrahita, J. A., Loboa, E. G. & Olby, N. J. Alternating Current Electric Fields of Varying Frequencies: Effects on Proliferation and Differentiation of Porcine Neural Progenitor Cells. https://home.liebertpub.com/cell 15, 405–412 (2013).

109. Matos, M. A. & Cicerone, M. T. Alternating current electric field effects on neural stem cell viability and differentiation. Biotechnol Prog 26, 664–670 (2010).

110. Yu, T. H., Wu, Y. J., Chien, M. E. & Hsu, K. Sen. Multisession Anodal Transcranial Direct Current Stimulation Enhances Adult Hippocampal Neurogenesis and Context Discrimination in Mice. Journal of Neuroscience 43, 635–646 (2023).

111. Trinchero, M. F. et al. Audiovisual gamma stimulation enhances hippocampal neurogenesis and neural circuit plasticity in aging mice. bioRxiv 2025.01.13.632794 (2025) doi:10.1101/2025.01.13.632794.

112. Islam, M. R. et al. Multisensory gamma stimulation enhances adult neurogenesis and improves cognitive function in male mice with Down Syndrome. PLoS One 20, e0317428 (2025).

113. Landers, J. et al. Carbon Nanotube Composites as Multifunctional Substrates for In Situ Actuation of Differentiation of Human Neural Stem Cells. Adv Healthc Mater 3, 1745–1752 (2014).

114. Seidlits, S. K. et al. Peptide-modified, hyaluronic acid-based hydrogels as a 3D culture platform for neural stem/progenitor cell engineering. J Biomed Mater Res A 107, 704–718 (2019).

115. Krautwald, K., Mahnke, L. & Angenstein, F. Electrical stimulation of the lateral entorhinal cortex causes a frequency-specific BOLD response pattern in the rat brain. Front Neurosci 13, 437573 (2019).

116. Abe, Y., Tsurugizawa, T., Le Bihan, D. & Ciobanu, L. Spatial contribution of hippocampal BOLD activation in high-resolution fMRI. Sci Rep 9, 1–9 (2019).

117. Kee, N., Sivalingam, S., Boonstra, R. & Wojtowicz, J. M. The utility of Ki-67 and BrdU as proliferative markers of adult neurogenesis. J Neurosci Methods 115, 97–105 (2002).

118. Hochgerner, H., Zeisel, A., Lönnerberg, P. & Linnarsson, S. Conserved properties of dentate gyrus neurogenesis across postnatal development revealed by single-cell RNA sequencing. Nature Neuroscience 2018 21:2 21, 290–299 (2018).

119. Lavoie, H., Gagnon, J. & Therrien, M. ERK signalling: a master regulator of cell behaviour, life and fate. Nat Rev Mol Cell Biol 21, 607–632 (2020).

120. Li, Z., Theus, M. H. & Wei, L. Role of ERK 1/2 signaling in neuronal differentiation of cultured embryonic stem cells. Dev Growth Differ 48, 513–523 (2006).

121. He, L. et al. Electrical stimulation at nanoscale topography boosts neural stem cell neurogenesis through the enhancement of autophagy signaling. Biomaterials 268, 120585 (2021).

122. Ryu, Y. et al. Cellular signaling pathways in the nervous system activated by various mechanical and electromagnetic stimuli. Front Mol Neurosci 17, 1427070 (2024).

123. Joo, M. C. et al. Effect of electrical stimulation on neural regeneration via the p38-RhoA and ERK1/2-Bcl-2 pathways in spinal cord-injured rats. Neural Regen Res 13, 340–346 (2018).

124. Kowiański, P. et al. BDNF: A Key Factor with Multipotent Impact on Brain Signaling and Synaptic Plasticity. Cell Mol Neurobiol 38, 579–593 (2018).

125. Caldeira, M. V. et al. BDNF regulates the expression and traffic of NMDA receptors in cultured hippocampal neurons. Molecular and Cellular Neuroscience 35, 208–219 (2007).

126. Chen, Y. H. et al. Quetiapine and repetitive transcranial magnetic stimulation ameliorate depression-like behaviors and up-regulate the proliferation of hippocampal-derived neural stem cells in a rat model of depression: The involvement of the BDNF/ERK signal pathway. Pharmacol Biochem Behav 136, 39–46 (2015).

127. Dalise, S. et al. Biological effects of dosing aerobic exercise and neuromuscular electrical stimulation in rats. Sci Rep 7, 1–13 (2017).

128. Luo, J. et al. High-frequency repetitive transcranial magnetic stimulation (rTMS) improves functional recovery by enhancing neurogenesis and activating BDNF/TrKB signaling in ischemic rats. Int J Mol Sci 18, (2017).

129. Mercerón-Martínez, D. et al. Amygdala electrical stimulation inducing spatial memory recovery produces an increase of hippocampal bdnf and arc gene expression. Brain Res Bull 124, 254–261 (2016).

130. Willand, M. P. et al. Electrical muscle stimulation elevates intramuscular BDNF and GDNF mRNA following peripheral nerve injury and repair in rats. Neuroscience 334, 93–104 (2016).

131. Ghorbani, M. et al. Impacts of epidural electrical stimulation on Wnt signaling, FAAH, and BDNF following thoracic spinal cord injury in rat. J Cell Physiol 235, 9795–9805 (2020).

132. Esmaeilpour, Z., Kronberg, G., Reato, D., Parra, L. C. & Bikson, M. Temporal interference stimulation targets deep brain regions by modulating neural oscillations. Brain Stimul 14, 55–65 (2021).

133. Luff, C. E. et al. The neuron mixer and its impact on human brain dynamics In brief. (2024) doi:10.1016/j.celrep.2024.114274.

134. Luff, C. E., Dzialecka, P., Acerbo, E., Williamson, A. & Grossman, N. Pulse-width modulated temporal interference (PWM-TI) brain stimulation. Brain Stimul 17, 92–103 (2024).

135. Mirzakhalili, E., Barra, B., Capogrosso, M. & Lempka, S. F. Biophysics of Temporal Interference Stimulation. Cell Syst 11, 557–572.e5 (2020).

136. Zhao, Y. et al. Electrical Property Characterization of Neural Stem Cells in Differentiation. PLoS One 11, e0158044 (2016).

137. Prè, D. et al. A Time Course Analysis of the Electrophysiological Properties of Neurons Differentiated from Human Induced Pluripotent Stem Cells (iPSCs). PLoS One 9, e103418 (2014).

138. Franz, D., Olsen, H. L., Klink, O. & Gimsa, J. Automated and manual patch clamp data of human induced pluripotent stem cell-derived dopaminergic neurons. Scientific Data 2017 4:1 4, 1–11 (2017).

139. Rakovic, A. et al. Electrophysiological Properties of Induced Pluripotent Stem Cell-Derived Midbrain Dopaminergic Neurons Correlate With Expression of Tyrosine Hydroxylase. Front Cell Neurosci 16, 817198 (2022).

140. Mirsadeghi, S. et al. Development of membrane ion channels during neural differentiation from human embryonic stem cells. Biochem Biophys Res Commun 491, 166–172 (2017).

141. Smith, R. S. & Walsh, C. A. Ion Channel Functions in Early Brain Development. Trends Neurosci 43, 103–114 (2020).

142. Ernsberger, U. Regulation of gene expression during early neuronal differentiation: Evidence for patterns conserved across neuron populations and vertebrate classes. Cell Tissue Res 348, 1–27 (2012).

143. Rasmussen, R. N., Asiminas, A., Carlsen, E. M. M., Kjaerby, C. & Smith, N. A. Astrocytes: integrators of arousal state and sensory context. Trends Neurosci 46, 418–425 (2023).

144. Vedam-Mai, V. et al. Deep brain stimulation and the role of astrocytes. Molecular Psychiatry 2012 17:2 17, 124–131 (2011).

145. McNeill, J., Rudyk, C., Hildebrand, M. E. & Salmaso, N. Ion Channels and Electrophysiological Properties of Astrocytes: Implications for Emergent Stimulation Technologies. Front Cell Neurosci 15, 644126 (2021).

146. Pasti, L., Volterra, A., Pozzan, T. & Carmignoto, G. Intracellular Calcium Oscillations in Astrocytes: A Highly Plastic, Bidirectional Form of Communication between Neurons and Astrocytes In Situ. Journal of Neuroscience 17, 7817–7830 (1997).

147. Chiareli, R. A. et al. The Role of Astrocytes in the Neurorepair Process. Front Cell Dev Biol 9, (2021).

148. de Majo, M., Koontz, M., Rowitch, D. & Ullian, E. M. An update on human astrocytes and their role in development and disease. Glia 68, 685–704 (2020).

149. Ahtiainen, A. et al. Stimulation of Neurons and Astrocytes via Temporally Interfering Electric Fields. bioRxiv 2023.10.30.564774 (2023) doi:10.1101/2023.10.30.564774.

150. Patel, R. et al. Ile-Lys-Val-ala-Val (IKVAV) peptide for neuronal tissue engineering. Polym Adv Technol 30, 4–12 (2019).

151. Silva, G. A. et al. Selective Differentiation of Neural Progenitor Cells by High-Epitope Density Nanofibers. Science (1979) 303, 1352–1355 (2004).

152. Peressotti, S., Koehl, G. E., Goding, J. A. & Green, R. A. Self-Assembling Hydrogel Structures for Neural Tissue Repair. ACS Biomater Sci Eng 7, 4136–4163 (2021).

153. Hyvärinen, T. et al. Co-stimulation with IL-1β and TNF-α induces an inflammatory reactive astrocyte phenotype with neurosupportive characteristics in a human pluripotent stem cell model system. Scientific Reports 2019 9:1 9, 1–15 (2019).

154. Iaccarino, H. F. et al. Gamma frequency entrainment attenuates amyloid load and modifies microglia. Nature 2016 540:7632 540, 230–235 (2016).

155. Liu, Q. et al. Intensity-dependent gamma electrical stimulation regulates microglial activation, reduces beta-amyloid load, and facilitates memory in a mouse model of Alzheimer’s disease. Cell Biosci 13, 1–14 (2023).

156. Prichard, A. et al. Brain rhythms control microglial response and cytokine expression via NF-κB signaling. Sci Adv 9, (2023).

157. Song, J. et al. Neuronal circuitry mechanism regulating adult quiescent neural stem-cell fate decision. Nature 2012 489:7414 489, 150–154 (2012).

158. Remmers, C. L., Castillon, C. C. M., Armstrong, J. N. & Contractor, A. Recruitment of parvalbumin and somatostatin interneuron inputs to adult born dentate granule neurons. Scientific Reports 2020 10:1 10, 1–11 (2020).

159. Strüber, M., Sauer, J. F. & Bartos, M. Parvalbumin expressing interneurons control spike-phase coupling of hippocampal cells to theta oscillations. Scientific Reports 2022 12:1 12, 1–10 (2022).

160. Amilhon, B. et al. Parvalbumin Interneurons of Hippocampus Tune Population Activity at Theta Frequency. Neuron 86, 1277–1289 (2015).

161. Wessel, M. J. et al. Noninvasive theta-burst stimulation of the human striatum enhances striatal activity and motor skill learning. Nature Neuroscience 2023 26:11 26, 2005–2016 (2023).

162. Missey, F. et al. Orientation of Temporal Interference for Non-invasive Deep Brain Stimulation in Epilepsy. Front Neurosci 15, 633988 (2021).

163. Acerbo, E. et al. Focal non-invasive deep-brain stimulation with temporal interference for the suppression of epileptic biomarkers. Front Neurosci 16, 945221 (2022).

164. Ohira, K., Hagihara, H., Miwa, M., Nakamura, K. & Miyakawa, T. Fluoxetine-induced dematuration of hippocampal neurons and adult cortical neurogenesis in the common marmoset. Mol Brain 12, (2019).

165. Vallejo-Giraldo, C. et al. Attenuated Glial Reactivity on Topographically Functionalized Poly(3,4-Ethylenedioxythiophene):P-Toluene Sulfonate (PEDOT:PTS) Neuroelectrodes Fabricated by Microimprint Lithography. Small 14, 1800863 (2018).

166. Thompson, L. H. & Parish, C. L. Transplantation of fetal midbrain dopamine progenitors into a rodent model of Parkinson’s disease. Methods in Molecular Biology 1059, 169–180 (2013).

167. Panayotov, I. V., Orti, V., Cuisinier, F. & Yachouh, J. Polyetheretherketone (PEEK) for medical applications. J Mater Sci Mater Med 27, 1–11 (2016).

168. Merrill, D. R., Bikson, M. & Jefferys, J. G. R. Electrical stimulation of excitable tissue: design of efficacious and safe protocols. J Neurosci Methods 141, 171–198 (2005).

169. Álvarez, Z. et al. Bioactive scaffolds with enhanced supramolecular motion promote recovery from spinal cord injury. Science (1979) 374, 848–856 (2021).

170. Adler, J. & Parmryd, I. Quantifying colocalization by correlation: The pearson correlation coefficient is superior to the Mander’s overlap coefficient. Cytometry Part A 77, 733–742 (2010).

171. Rattus_norvegicus - Ensembl genome browser 114. https://www.ensembl.org/Rattus_norvegicus/Info/Index.

172. Mortazavi, A., Williams, B. A., McCue, K., Schaeffer, L. & Wold, B. Mapping and quantifying mammalian transcriptomes by RNA-Seq. Nature Methods 2008 5:7 5, 621–628 (2008).

173. Anders, S. & Huber, W. Differential expression analysis for sequence count data. Genome Biol 11, 1–12 (2010).

174. Benjamini, Y. Discovering the false discovery rate. J R Stat Soc Series B Stat Methodol 72, 405–416 (2010).

175. Enrichr: a comprehensive gene set enrichment analysis web server 2016 update. vol. 44 https://maayanlab.cloud/Enrichr/.

176. Gee, K. R. et al. Chemical and physiological characterization of fluo-4 Ca2+-indicator dyes. Cell Calcium 27, 97–106 (2000).

177. scipy.signal.find_peaks — SciPy v1.11.4 Manual. https://docs.scipy.org/doc/scipy-1.11.4/reference/generated/scipy.signal.find_peaks.html.

178. Hurst, J. L. & West, R. S. Taming anxiety in laboratory mice. Nature Methods 2010 7:10 7, 825–826 (2010).

179. Grill, W. M. Temporal pattern of electrical stimulation is a new dimension of therapeutic innovation. Curr Opin Biomed Eng 8, 1–6 (2018).

180. Albensi, B. C., Oliver, D. R., Toupin, J. & Odero, G. Electrical stimulation protocols for hippocampal synaptic plasticity and neuronal hyper-excitability: Are they effective or relevant? Exp Neurol 204, 1–13 (2007).

181. van Goethem, N. P., van Hagen, B. T. J. & Prickaerts, J. Assessing spatial pattern separation in rodents using the object pattern separation task. Nat Protoc 13, 1763–1792 (2018).

182. Blackmore, D. G., Brici, D. & Walker, T. L. Protocol for three alternative paradigms to test spatial learning and memory in mice. STAR Protoc 3, (2022).

183. Coscia, F. et al. A streamlined mass spectrometry–based proteomics workflow for large-scale FFPE tissue analysis. Journal of Pathology 251, 100–112 (2020).

184. Schindelin, J. et al. Fiji - an Open Source platform for biological image analysis. Nat Methods 9, 676–682 (2012).

185. Sun, X. et al. Histogram-based normalization technique on human brain magnetic resonance images from different acquisitions. Biomed Eng Online 14, 1–17 (2015).

186. Li, P., Wang, H., Yu, M. & Li, Y. Overview of Image Smoothing Algorithms. J Phys Conf Ser 1883, 012024 (2021).

187. Romero-Zaliz, R. & Reinoso-Gordo, J. F. An updated review on watershed algorithms. Studies in Fuzziness and Soft Computing 358, 235–258 (2018).

188. Yu, Z. et al. Beyond t test and ANOVA: applications of mixed-effects models for more rigorous statistical analysis in neuroscience research. Neuron 110, 21–35 (2022).

189. Keselman, H. J., Kowalchuk, R. K., Algina, J. & Wolfinger, R. D. The analysis of repeated measurements: A comparison of mixed-model Satterthwaite F tests and a nonpooled adjusted degrees of freedom multivariate test. Commun Stat Theory Methods 28, 2967–2999 (1999).

190. Morrell, C. H. Likelihood Ratio Testing of Variance Components in the Linear Mixed-Effects Model Using Restricted Maximum Likelihood. Biometrics 54, 1560 (1998).

191. Sture Holm. A Simple Sequentially Rejective Multiple Test Procedure. Scandinavian Journal of Statistics 6, 65–70 (1979).

192. Lilliefors, H. W. On the Kolmogorov-Smirnov Test for Normality with Mean and Variance Unknown. J Am Stat Assoc 62, 399–402 (1967).

193. MacFarland, T. W. & Yates, J. M. Mann–Whitney U Test. Introduction to Nonparametric Statistics for the Biological Sciences Using R 103–132 (2016) doi:10.1007/978-3-319-30634-6_4.

194. Blanca, M. J., Arnau, J., García-Castro, F. J., Alarcón, R. & Bono, R. Non-normal Data in Repeated Measures ANOVA: Impact on Type I Error and Power. Psicothema 35, 21–29 (2023).

