## Supplementary material for "Temporal Interference Stimulation Enhances Neural Regeneration": Document S1: Document_S1.docx

**SUPPLEMENTAL INFORMATION – Document S1**

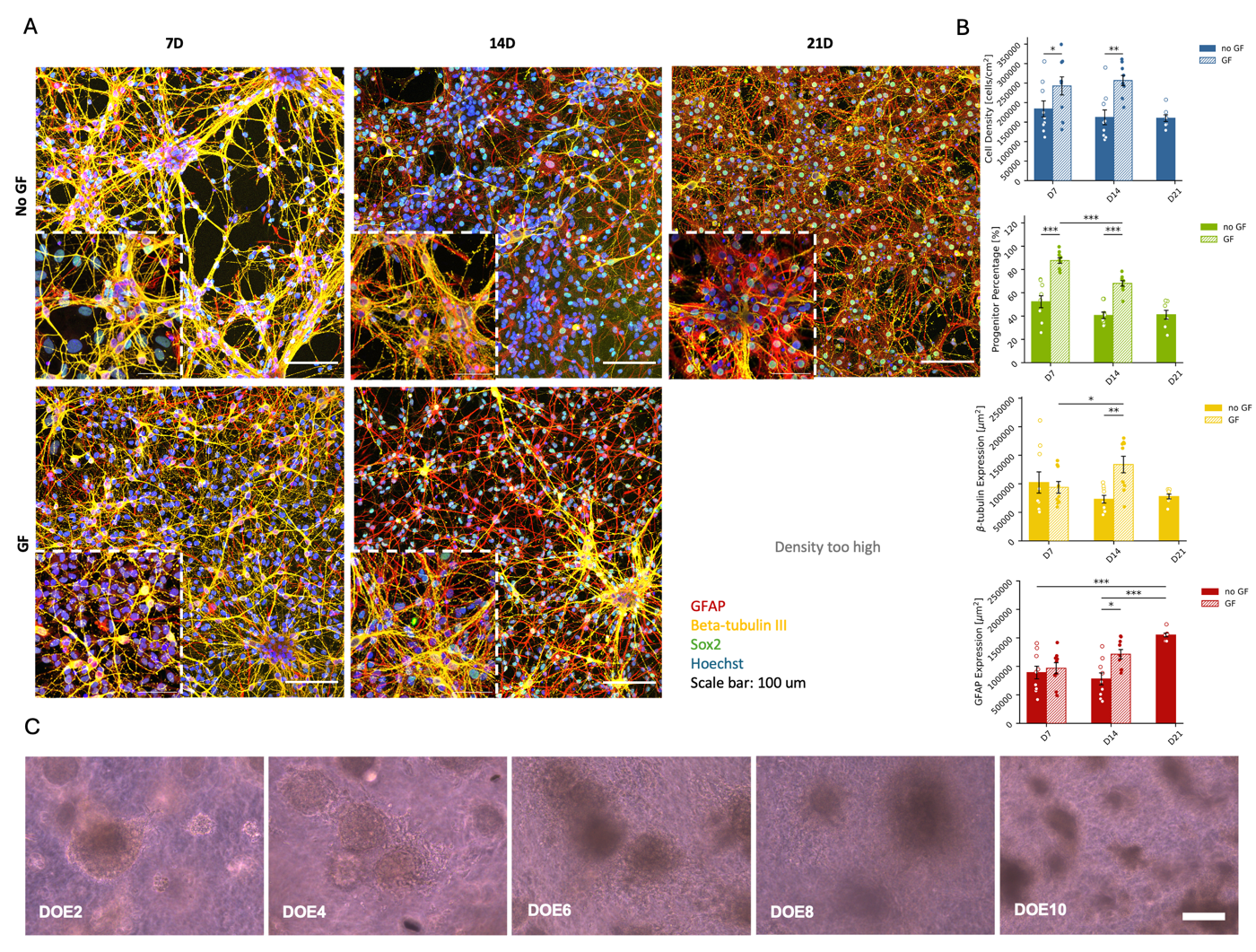

**Figure S1:** **Characterisation of primary NPC monolayer cultures, related to Figure 1 and Figure 2.**

(A) Representative immunofluorescence images of the culture at DIV7, DIV14 and DIV21. The “no GF” condition refers to the medium that were used in Figure 1. In the “GF” condition, the cells were kept in the proliferation medium (see methods for details). β-tubulin III: yellow, GFAP: red, Sox2: green, nuclear and Hoechst: blue, nuclei. Scale bar: 100 µm.
(B) Total cell density, progenitor cell (Sox2+) Percentage, β-tubulin III and GFAP expression at 7, 14 and 21 DIV. A t-test and a post-hoc Benjamini-Hockberg test was used to adjust for multiple comparisons between the means (values expressed as mean±SEM, biological repeats n=3, technical repeats N=7-9; *: p-value<0.05, **: p-value<0.01, ***: p-value<0.001).
(C) Representative phase-contrast pictures of neurospheres from day of ecapsulation (DOE) 0 to DOE10. Scale bar: 100 µm.

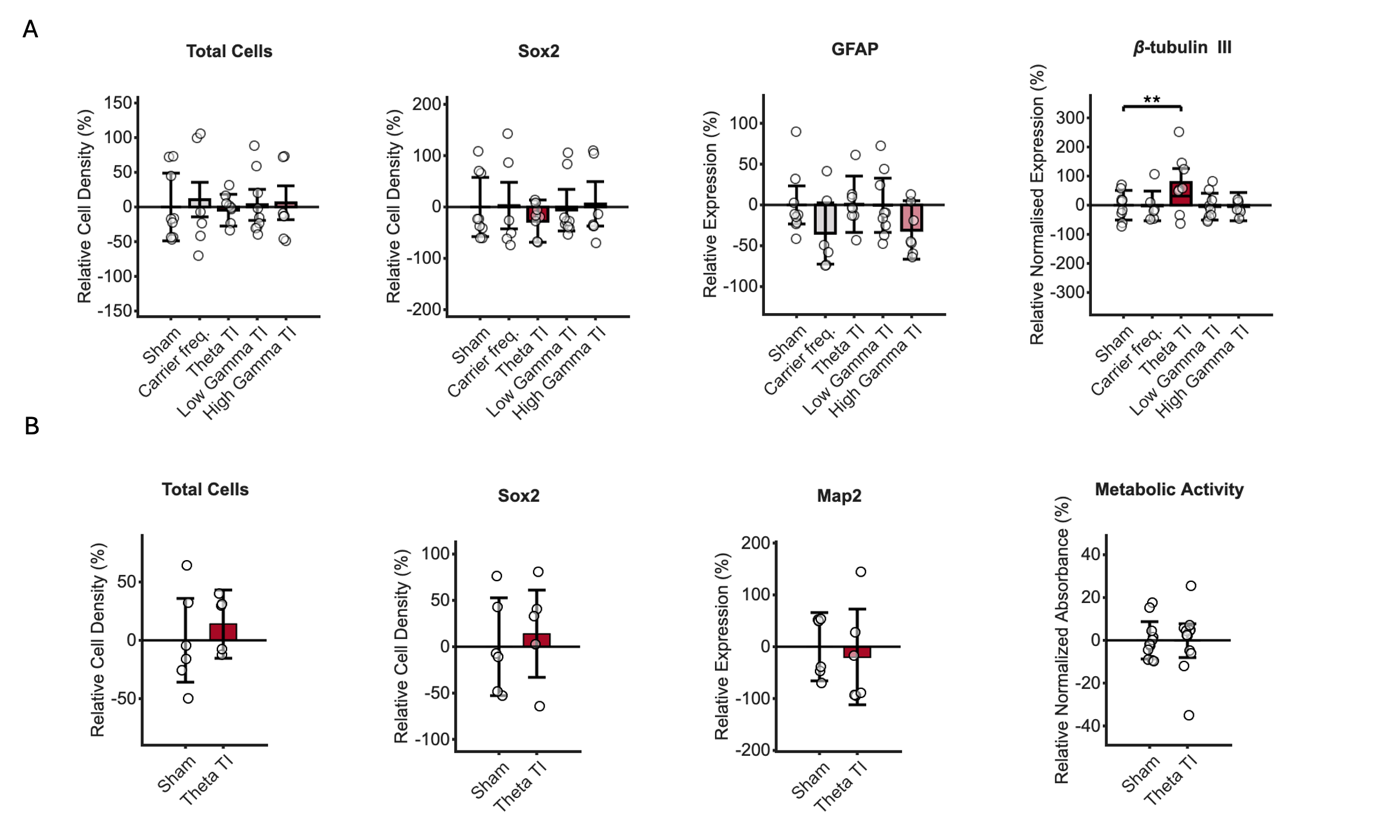

**Figure S2:** **Supplemental IHC quantifications related to Figure 1 and Figure 2.**

(A) Supplemental experimental outcomes of the NPC stimulation study across TI frequencies, related to Figure 1: total cell density, sox2+ cell density, GFAP expression in µm^2^.
(B) Supplemental experimental outcomes of the NPC stimulation study in the 3D model, related to Figure 3: Quantification of total cell density, sox2+ cell density, GFAP expression in µm^2^, metabolic activity.

To compute the statistical significance, a LMM was implemented. The data is presented as the percentual change of each metric compared to the sham control (data points are the percentual change from the average of the sham, bar plots show the model predicted-means±95% CI, n=3 number of animals, N=6-9 number of well-plates per condition considered as data points. ^∗∗^p<0.01, with p displaying the p-value adjusted for multiple comparisons with BH correction where required).

**Table S1: results of LMM statistical analyses related to the immunofluorescence quantifications of the embryonic NPCs cultured in a monolayer, Figure 1B, S2 and methods section.** A linear mixed effect t mode (LMM) was implemented to compare the data across frequencies, where the fixed effect was set as the stimulation condition (Carrier, Theta, Low Gamma band, High Gamma band) with sham as a reference, and the random effect was set as the biological repeat (n=3) from which each data point was generated (total replicas N=6-9). The F statistics and degrees of freedom were calculated with an ANOVA, Satterthwaite method^1^. To evaluate the significance of the biological repeat effect, we performed a likelihood ratio test^2^ by comparing the full model to a reduced model excluding the biological repeat (random effect, where c^2^=2 × (log-likelihood of the full model – log-likelihood of the model without random effect). This test determined whether inclusion of the biological effect significantly improved model fit. The estimate represents the distance from the intercept, and the standard error of the mean (SE), t-statistic (tStat), freedom (DF) and p values (pValue) are reported. In case of multiple comparisons, the results were corrected with Bonferroni-Holm (BH) post-hoc correction^3^ . We identified outliers using a threshold of z-score >3 on the model residuals. If outliers were detected, we ran the analysis both with and without outliers. Here, we report the most conservative results.

| Fixed effect – Alamar blue | DF1 | DF2 | F | pValue |  |
| --- | --- | --- | --- | --- | --- |
| (Intercept) | 1 | 4.622791447 | 395.5636309 | 1.17472E-05 |  |
| Treatment | 4 | 25.00775194 | 2.864300251 | 0.044114755 |  |
| Random Effect | **DF** | c^2^ | **pValue** |  |  |
| Biological Repeat | 1 | 5.41973173 | 0.019910419 |  |  |
| Name | **Estimate** | **SE** | **tStat** | **pValue** | **Adj pValue** |
| (Intercept) | 0.389225624 | 0.019570109 | 19.88878153 | 1.49709E-15 |  |
| Treatment_Carrier | 0.009764525 | 0.020770814 | 0.470107946 | 0.642904708 | 2.571618831 |
| Treatment_Theta | 0.050292657 | 0.018577981 | 2.707111042 | 0.012871466 | 0.012871466 |
| Treatment_Low Gamma Band | 0.047776053 | 0.019518227 | 2.447766076 | 0.022817936 | 0.030423915 |
| Treatment_High Gamma Band | 0.013625103 | 0.018577981 | 0.733400667 | 0.471057753 | 0.942115505 |
| Fixed effect – β-tubulin III expression | **DF1** | **DF2** | **F** | **pValue** |  |
| (Intercept) | 1 | 7.59756811 | 15.93557376 | 0.00443654 |  |
| Treatment | 4 | 36.0733165 | 2.953409537 | 0.032907519 |  |
| Random Effect | **DF** | c^2^ | **pValue** |  |  |
| Biological Repeat | 1 | 6.76672828 | 0.009287294 |  |  |
| Name | **Estimate** | **SE** | **tStat** | **pValue** | **Adj pValue** |
| (Intercept) | 43165.4507 | 10813.155 | 3.991938597 | 0.00033139 |  |
| Treatment_Carrier | 2301.52097 | 11978.7706 | 0.19213332 | 0.848780737 | 1.697561475 |
| Treatment_Theta | 32485.765 | 11059.8109 | 2.937280321 | 0.005906133 | 0.005906133 |
| Treatment_Low Gamma Band | 576.620859 | 10714.1381 | 0.053818688 | 0.957394613 | 3.829578451 |
| Treatment_High Gamma Band | 5009.63605 | 11472.7372 | 0.436655696 | 0.665121658 | 0.886828877 |
| Fixed effect – Hoechst & Sox2 correlation | **DF1** | **DF2** | **F** | **pValue** |  |
| (Intercept) | 1 | 39.00000077 | 236.7918439 | 3.71044E-18 |  |
| Treatment | 4 | 39.00000077 | 1.632702084 | 0.185459311 |  |
| Random Effect | **DF** | c^2^ | **pValue** |  |  |
| Biological Repeat | 1 | 0 | 1 |  |  |
| Name | **Estimate** | **SE** | **tStat** | **pValue** | **Adj pValue** |
| (Intercept) | 0.611362022 | 0.039729682 | 15.38804224 | 6.93047E-17 |  |
| Treatment_Carrier | -0.11593166 | 0.062818143 | -1.845512389 | 0.073686679 | 0.098248906 |
| Treatment_Theta | -0.136952105 | 0.057915466 | -2.364689687 | 0.02389473 | 0.02389473 |
| Treatment_Low Gamma Band | -0.064815535 | 0.056186255 | -1.15358347 | 0.256716111 | 1.026864445 |
| Treatment_High Gamma Band | -0.083362254 | 0.060065633 | -1.387852756 | 0.174210493 | 0.348420986 |
| Fixed effect – Total cell density | **DF1** | **DF2** | **F** | **pValue** |  |
| (Intercept) | 1 | 3.55112472 | 17.30586571 | 0.018061295 |  |
| Treatment | 4 | 36.0040309 | 0.411879442 | 0.798891631 |  |
| Random Effect | **DF** | c^2^ | **pValue** |  |  |
| Biological Repeat | 1 | 41.1953161 | 1.37752E-10 |  |  |
| Name | **Estimate** | **SE** | **tStat** | **pValue** | **Adj pValue** |
| (Intercept) | 208220.1884 | 50052.5457 | 4.160031936 | 0.000204208 |  |
| Treatment_Carrier | 21837.23873 | 25738.865 | 0.848414985 | 0.40214068 | 0.40214068 |
| Treatment_Theta | -9186.479162 | 23771.1026 | -0.386455744 | 0.701567213 | 1.403134426 |
| Treatment_Low Gamma Band | 6645.592869 | 23021.5407 | 0.288668467 | 0.774587792 | 3.098351168 |
| Treatment_High Gamma Band | 12358.10245 | 24659.027 | 0.50115937 | 0.619488364 | 0.825984485 |
| Fixed effect – Sox2+ cell density | **DF1** | **DF2** | **F** | **pValue** |  |
| (Intercept) | 1 | 4.53537654 | 12.47702341 | 0.019640054 |  |
| Treatment | 4 | 36.0042351 | 0.760862874 | 0.557634693 |  |
| Random Effect | **DF** | c^2^ | **pValue** |  |  |
| Biological Repeat | 1 | 19.0202479 | 1.29339E-05 |  |  |
| Name | **Estimate** | **SE** | **tStat** | **pValue** | **Adj pValue** |
| (Intercept) | 202223.3997 | 57250.0556 | 3.532283031 | 0.001208536 |  |
| Treatment_Carrier | 5005.529696 | 44966.73 | 0.111316293 | 0.912020091 | 3.648080365 |
| Treatment_Theta | -55881.78184 | 41525.6122 | -1.345718434 | 0.187299183 | 0.187299183 |
| Treatment_Low Gamma Band | -11395.10203 | 40219.466 | -0.283323056 | 0.778647232 | 1.038196309 |
| Treatment_High Gamma Band | 11324.26023 | 43076.5504 | 0.262886887 | 0.794224282 | 1.588448565 |
| Fixed effect – GFAP expression | **DF1** | **DF2** | **F** | **pValue** |  |
| (Intercept) | 1 | 39.000001 | 73.31930262 | 1.70872E-10 |  |
| Treatment | 4 | 39.0000009 | 1.907085884 | 0.128622668 |  |
| Random Effect | **DF** | c^2^ | **pValue** |  |  |
| Biological Repeat | 1 | 0 | 1 |  |  |
| Name | **Estimate** | **SE** | **tStat** | **pValue** | **Adj pValue** |
| (Intercept) | 49226.87258 | 5749.01025 | 8.56266913 | 5.29568E-10 |  |
| Treatment_Carrier | -17079.83038 | 9089.98334 | -1.878972683 | 0.068843221 | 0.068843221 |
| Treatment_Theta | 385.5262879 | 8380.55055 | 0.046002501 | 0.963577398 | 1.927154796 |
| Treatment_Low Gamma Band | -174.5965496 | 8130.32826 | -0.021474723 | 0.982992503 | 3.931970013 |
| Treatment_High Gamma Band | -15219.08735 | 8691.68652 | -1.750993587 | 0.088965729 | 0.118620972 |
| Fixed effect – β-tubulin III expression normalised | **DF1** | **DF2** | **F** | **pValue** |  |
| (Intercept) | 1 | 6.19887699 | 16.50326205 | 0.00619303 |  |
| Treatment | 4 | 35.9855826 | 4.690068764 | 0.003771956 |  |
| Random Effect | **DF** | c^2^ | **pValue** |  |  |
| Biological Repeat | 1 | 8.85508561 | 0.002922734 |  |  |
| Name | **Estimate** | **SE** | **tStat** | **pValue** | **Adj pValue** |
| (Intercept) | 208.4709404 | 51.3169253 | 4.062420713 | 0.000270671 |  |
| Treatment_Carrier | -5.113829672 | 51.9001818 | -0.098532018 | 0.922088475 | 3.6883539 |
| Treatment_Theta | 163.2256656 | 47.922511 | 3.406033247 | 0.001708233 | 0.001708233 |
| Treatment_Low Gamma Band | -9.377128828 | 46.4209338 | -0.202002158 | 0.841118858 | 1.12149181 |
| Treatment_High Gamma Band | -9.997176731 | 49.7120057 | -0.201101859 | 0.841817185 | 1.68363437 |

**Table S2: results of LLM statistical analyses related to the live-cell calcium imaging of the embryonic NPCs cultured in a monolayer, Figure 1D, 1E and methods section.** A linear mixed effect t mode (LMM) was implemented to compare the data across frequencies, where the fixed effect was set as the stimulation condition (Theta only) with sham as a reference, and the random effect was set as the biological repeat (n=3) from which each data point was generated (total replicas N=6-9). The F statistics and degrees of freedom were calculated with an ANOVA, Satterthwaite method^1^. To evaluate the significance of the biological repeat effect, we performed a likelihood ratio test^2^ by comparing the full model to a reduced model excluding the biological repeat (random effect, where c^2^=2 × (log-likelihood of the full model – log-likelihood of the model without random effect). This test determined whether inclusion of the biological effect significantly improved model fit. The estimate represents the distance from the intercept, and the standard error of the mean (SE), t-statistic (tStat), degrees of freedom (DF) and p values (pValue) are reported. We identified outliers using a threshold of z-score >3 on the model residuals. If outliers were detected, we ran the analysis both with and without outliers. Here, we report the most conservative results.

| Fixed effect – Peak number | DF1 | DF2 | F | pValue |
| --- | --- | --- | --- | --- |
| (Intercept) | 1 | 15 | 4.8711 | 0.04330745 |
| Treatment | 1 | 15 | 4.6186 | 0.04835884 |
| Random Effect | **DF** | c^2^ | **pValue** |  |
| Biological Repeat | 1 | 0 | 1 |  |
| Name | **Estimate** | **SE** | **tStat** | **pValue** |
| (Intercept) | 3 | 1.36 | 2.2071 | 0.045897906 |
| Treatment_Theta | 4 | 1.86 | 2.1491 | 0.051041746 |
| Fixed effect – peak length | **DF1** | **DF2** | **F** | **pValue** |
| (Intercept) | 1 | 15 | 192.51 | 5.80548E-10 |
| Treatment | 1 | 15 | 7.2981 | 0.016409559 |
| Random Effect | **DF** | c^2^ | **pValue** |  |
| Biological Repeat | 1 | 0 | 1 |  |
| Name | **Estimate** | **SE** | **tStat** | **pValue** |
| (Intercept) | 1134.946572 | 81.8 | 13.875 | 3.59802E-09 |
| Treatment_Theta | -302.5873214 | 112 | -2.7015 | 0.018142219 |

**Table S3: results of statistical analyses of the stimulated 3D in vitro model of embryonic NPCs, related to Figure 2.** A linear mixed effect t mode (LMM) was implemented to compare the data across frequencies, where the fixed effect was set as the stimulation condition (Theta ony) with sham as a reference, and the random effect was set as the biological repeat (n=3) from which each data point was generated (total replicas N=6-9). The F statistics and degrees of freedom were calculated with an ANOVA, Satterthwaite method^1^. To evaluate the significance of the biological repeat effect, we performed a likelihood ratio test^2^ by comparing the full model to a reduced model excluding the biological repeat (random effect, where c^2^=2 × (log-likelihood of the full model – log-likelihood of the model without random effect). This test determined whether inclusion of the biological effect significantly improved model fit. The estimate represents the distance from the intercept, and the standard error of the mean (SE), t-statistic (tStat), freedom (DF) and p values (pValue) are reported. We identified outliers using a threshold of z-score >3 on the model residuals. If outliers were detected, we ran the analysis both with and without outliers. Here, we report the most conservative results.

| Fixed effect – β-tubulin III expression | DF1 | DF2 | F | pValue |
| --- | --- | --- | --- | --- |
| (Intercept) | 1 | 12 | 30.986495 | 0.000122494 |
| Treatment | 1 | 12 | 0.92348564 | 0.355525743 |
| Random Effect | **DF** | c^2^ | **pValue** |  |
| Biological Repeat | 1 | 0 | 1 |  |
| Name | **Estimate** | **SE** | **tStat** | **pValue** |
| (Intercept) | 41281.734 | 7416.03386 | 5.56655144 | 0.000238532 |
| Treatment_Theta | -10078.6364 | 10487.8557 | -0.9609816 | 0.359221296 |
| Fixed effect – β-tubulin III & Map2 correlation | **DF1** | **DF2** | **F** | **pValue** |
| (Intercept) | 1 | 7.275304383 | 67.8557004 | 6.09369E-05 |
| Treatment | 1 | 9.40260111 | 6.22670591 | 0.033078933 |
| Random Effect | **DF** | c^2^ | **pValue** |  |
| Biological Repeat | 1 | 0.007149005 | 0.93261773 |  |
| Name | **Estimate** | **SE** | **tStat** | **pValue** |
| (Intercept) | 0.20046392 | 0.024335655 | 8.23745716 | 2.78342E-06 |
| Treatment_Theta | 0.090340581 | 0.036203762 | 2.49533683 | 0.028155204 |
| Fixed effect – GFAP expression | **DF1** | **DF2** | **F** | **pValue** |
| (Intercept) | 1 | 11 | 24.1863745 | 0.000458341 |
| Treatment | 1 | 11 | 13.6247499 | 0.003555886 |
| Random Effect | **DF** | c^2^ | **pValue** |  |
| Biological Repeat | 1 | 0 | 1 |  |
| Name | **Estimate** | **SE** | **tStat** | **pValue** |
| (Intercept) | 48607.6979 | 9883.70253 | 4.91796447 | 0.000826921 |
| Treatment_Theta | 54112.2107 | 14659.9 | 3.69117189 | 0.004988317 |
| Fixed effect – Hoechst & Sox2 correlation | **DF1** | **DF2** | **F** | **pValue** |
| (Intercept) | 1 | 5.041598309 | 152.653993 | 5.82188E-05 |
| Treatment | 1 | 7.745054979 | 0.29360491 | 0.603153921 |
| Random Effect | **DF** | c^2^ | **pValue** |  |
| Biological Repeat | 1 | 0.435459802 | 0.509322 |  |
| Name | **Estimate** | **SE** | **tStat** | **pValue** |
| (Intercept) | 0.47514778 | 0.038456931 | 12.3553225 | 6.003E-07 |
| Treatment_Theta | 0.024891104 | 0.045936987 | 0.54185322 | 0.601074929 |
| Fixed effect – total cell density | **DF1** | **DF2** | **F** | **pValue** |
| (Intercept) | 1 | 4.05991566 | 40.2306652 | 0.003008572 |
| Treatment | 1 | 8.14594155 | 1.1648321 | 0.311382925 |
| Random Effect | **DF** | c^2^ | **pValue** |  |
| Biological Repeat | 1 | 3.75148242 | 0.0527607 |  |
| Name | **Estimate** | **SE** | **tStat** | **pValue** |
| (Intercept) | 631742.979 | 99600.5686 | 6.34276479 | 0.000134017 |
| Treatment_Theta | 88487.9878 | 81988.4458 | 1.07927388 | 0.308539158 |
| Fixed effect – Sox2+ cell density | **DF1** | **DF2** | **F** | **pValue** |
| (Intercept) | 1 | 4.19970791 | 18.1987322 | 0.011695124 |
| Treatment | 1 | 8.11455852 | 0.44563556 | 0.522942957 |
| Random Effect | **DF** | c^2^ | **pValue** |  |
| Biological Repeat | 1 | 2.82798465 | 0.09263454 |  |
| Name | **Estimate** | **SE** | **tStat** | **pValue** |
| (Intercept) | 307893.414 | 72173.8432 | 4.26599722 | 0.002092597 |
| Treatment_Theta | 42712.2677 | 63982.7214 | 0.66755941 | 0.52116265 |
| Fixed effect – Map2 expression | **DF1** | **DF2** | **F** | **pValue** |
| (Intercept) | 1 | 12.0000005 | 11.6437559 | 0.005151362 |
| Treatment | 1 | 12.0000005 | 0.23970179 | 0.633249169 |
| Random Effect | **DF** | c^2^ | **pValue** |  |
| Biological Repeat | 1 | 0 | 1 |  |
| Name | **Estimate** | **SE** | **tStat** | **pValue** |
| (Intercept) | 50910.8618 | 14919.831 | 3.41229482 | 0.006632674 |
| Treatment_Theta | -10330.3382 | 21099.8273 | -0.48959349 | 0.63498333 |
| Fixed effect Metabolic activity | **DF1** | **DF2** | **F** | **pValue** |
| (Intercept) | 1 | 6.272475323 | 566.261017 | 2.18592E-07 |
| Treatment | 1 | 20.00000059 | 0.00112638 | 0.973559472 |
| Random Effect | **DF** | c^2^ | **pValue** |  |
| Biological Repeat | 1 | 3.911059657 | 0.04796934 |  |
| Name | **Estimate** | **SE** | **tStat** | **pValue** |
| (Intercept) | 0.515141033 | 0.021648002 | 23.7962396 | 3.42571E-17 |
| Treatment_Theta | -0.000660806 | 0.019689374 | -0.03356156 | 0.97352942 |

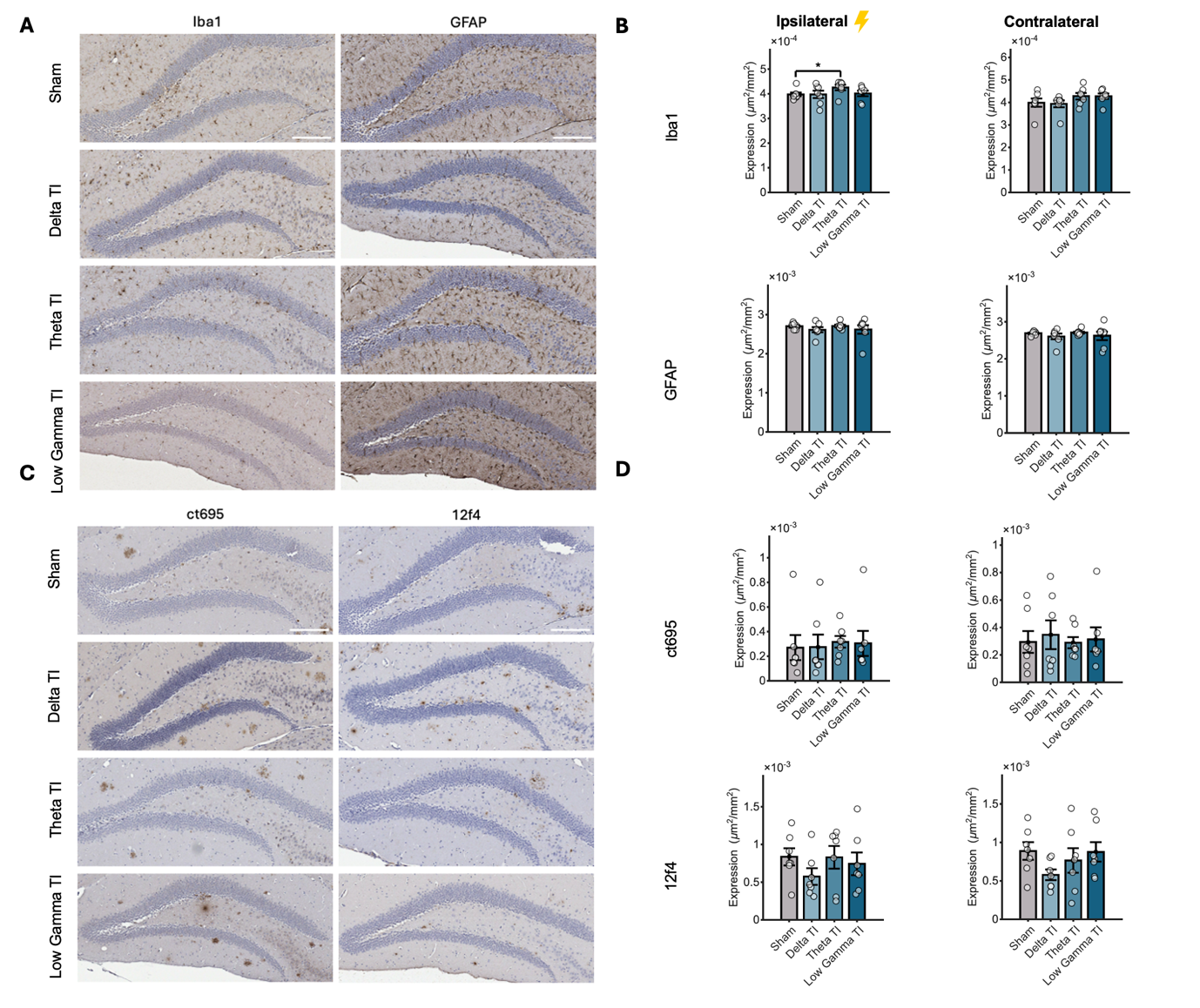

**Figure S3: TI stimulations lead to no significant difference in glial and amyloid-beta expression bilaterally in the short term, except for a theta-specific increase in Iba1 expression in the ipsilateral DG region. Related to Figure 3.**

(A) Representative brightfield images of ipsilateral DG stained with glial markers across conditions. Iba1 and GFAP were stained with DAB and counterstained with haematoxylin to quantify microglia and astrocytes. Scale bar: 200µm.
(B) Quantification of glial expressions in the ipsilateral and contralateral DG. The percentage of Iba1 and GFAP area coverage across frequencies is compared with the sham control. Each data point represents an individual mouse.
(C) Representative brightfield images of ipsilateral DG stained with amyloid-beta markers across conditions. ct695 and 12f4 were stained with DAB and counterstained with haematoxylin. Scale bar: 200µm.
(D) Quantification of amyloid-beta expression in the ipsilateral and contralateral DG. The percentage of ct695 and 12f4 area coverage across frequencies is compared with the sham control. Each data point represents an individual mouse.

Results are expressed as mean±standard error of the mean (SEM). Significance in (B) was calculated using an independent t-test if the data passed the normality test (Shapiro), or a Wilcoxon rank-sum test otherwise, and a post-hoc BH correction (n=24; sham: n=7, delta: n=6, theta: n=6, gamma: n=5; ∗p<0.05, ∗∗p<0.01, ∗∗∗p<0.001, with p displaying the p-value adjusted for multiple comparisons). Statistical analyses results can be found in **Table S4.**

**Table S4: results of statistical analyses of the IHC quantifications related to Figure 3 and S3.** The short-term study dataset was assessed for normality with the Shapiro test. If the condition passed the normality test (p-value of the Shapiro test > 0.05), an independent t-test was used. Otherwise, a non-parametric Wilcoxon rank-sum test was used to compare the means of each stimulation group to the sham group, for a total of four comparisons^4^. The results were corrected for multiple comparisons with Bonferroni correction **(**for all quantifications, n=24; sham: n=7, delta: n=6, theta: n=6, gamma: n=5).

| DCX expression | Test | Ipsilateral pValue | Ipsilateral Adj. pValue | Test | Contralateral pValue | Contralateral Adj. pValue |
| --- | --- | --- | --- | --- | --- | --- |
| Gamma vs Sham | t-test | 0.347382244 | 0.694764487 | t-test | 0.139868599 | 0.279737198 |
| Theta vs Sham | t-test | 0.004012178 | 0.012036533 | t-test | 0.000918052 | 0.002754157 |
| Delta vs Sham | t-test | 0.496563013 | 0.694764487 | t-test | 0.326135442 | 0.326135442 |
| Ki67+ cell density | **Test** | **Ipsilateral pValue** | **Ipsilateral Adj. pValue** | **Test** | **Contralateral pValue** | **Contralateral Adj. pValue** |
| Gamma vs Sham | Wilcoxon | 0.287878788 | 0.575757576 | Wilcoxon | 0.924242424 | 1 |
| Theta vs Sham | Wilcoxon | 0.016317016 | 0.048951049 | Wilcoxon | 0.123543124 | 0.370629371 |
| Delta vs Sham | Wilcoxon | 0.743589744 | 0.743589744 | Wilcoxon | 0.597902098 | 1 |
| Iba1 expression | **Test** | **Ipsilateral pValue** | **Ipsilateral Adj. pValue** | **Test** | **Contralateral pValue** | **Contralateral Adj. pValue** |
| Gamma vs Sham | Wilcoxon | 0.045454545 | 0.090909091 | t-test | 0.094436971 | 0.283310914 |
| Theta vs Sham | Wilcoxon | 0.01048951 | 0.031468531 | t-test | 0.128566395 | 0.283310914 |
| Delta vs Sham | Wilcoxon | 0.381118881 | 0.381118881 | t-test | 0.647858076 | 0.647858076 |
| GFAP expression | **Test** | **Ipsilateral pValue** | **Ipsilateral Adj. pValue** | **Test** | **Contralateral pValue** | **Contralateral Adj. pValue** |
| Gamma vs Sham | t-test | 0.263568333 | 0.790704999 | Wilcoxon | 0.318181818 | 0.954545455 |
| Theta vs Sham | t-test | 0.800063466 | 0.913243219 | Wilcoxon | 0.944055944 | 1 |
| Delta vs Sham | t-test | 0.45662161 | 0.913243219 | Wilcoxon | 0.757575758 | 1 |
| Ct695 expression | **Test** | **Ipsilateral pValue** | **Ipsilateral Adj. pValue** | **Test** | **Contralateral pValue** | **Contralateral Adj. pValue** |
| Gamma vs Sham | t-test | 0.833384073 | 1 | t-test | 0.50973733 | 1 |
| Theta vs Sham | t-test | 0.032617864 | 0.097853593 | t-test | 0.888566452 | 1 |
| Delta vs Sham | t-test | 0.60691593 | 1 | t-test | 0.458755863 | 1 |
| 12f4 expression | **Test** | **Ipsilateral pValue** | **Ipsilateral Adj. pValue** | **Test** | **Contralateral pValue** | **Contralateral Adj. pValue** |
| Gamma vs Sham | t-test | 0.57310044 | 1 | t-test | 0.723991928 | 1 |
| Theta vs Sham | t-test | 0.617007568 | 1 | t-test | 0.87510652 | 1 |
| Delta vs Sham | t-test | 0.112984962 | 0.338954886 | t-test | 0.056876759 | 0.170630277 |

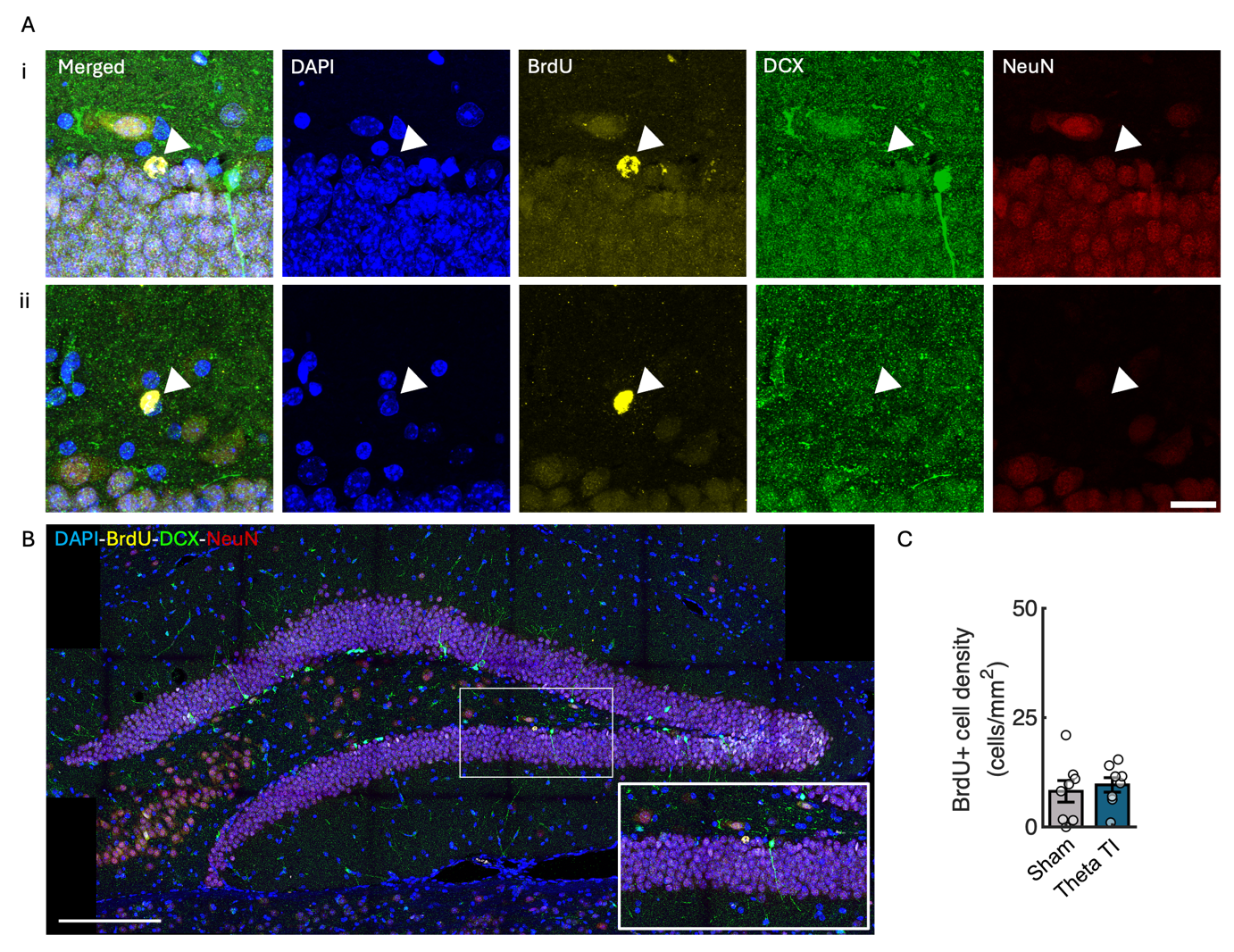

**Figure S4: No aberrant migration is detected after applying theta TI stimulation. Related to Figure 5.**

(A) Representative fluorescent micrographs of the BrdU+ cell quantification in the DG region, showing (i) mature GC cells co-expressing NeuN, and (ii) “other fate” cells not expressing NeuN nor DCX. Scale bar: 20 µm.
(B) Representative fluorescent photograph of the entire DG region quantified, including a zoomed in area. Scale bar: 200 µm.
(C) Quantification of BrdU+ cell density in the hilus of the DG to detect aberrant cell migration. Cell density in cells/mm^2^ for sham and theta stimulated mice. Each data point represents an individual mouse. Results are expressed as mean ± SEM (sham: n=8, theta TI: n=7).

**Table S5: Repeated measures ANOVA results of IHC data related to Figure 4.** The results were analysed with repeated measures ANOVA using the function 'aov_car’ from the ‘afex’ package in R. We We assessed the normality of the residuals with the Shapiro-Wilk test. We report deegrees pf freedom (DF), F statistic (F-value) and p-value for the model.

| Repeated measures ANOVA statistical results associated to DCX+ cell density | | | | | | |
| --- | --- | --- | --- | --- | --- | --- |
| GCL DCX+ cell density | **N_Sham_** | **N_Theta_** | **DF1** | **DF2** | **F-value** | ***p*-value** |
| Treatment (Sham, Theta) | 7 | 6 | 1 | 11 | 5.407995284 | 0.0402 |
| ROI (Ipsilateral, Contralateral) | 7 | 6 | 1 | 11 | 0.454196328 | 0.514 |
| Treatment x ROI | 7 | 6 | 1 | 11 | 0.049708116 | 0.828 |
| dGCL DCX+ cell density | **N_Sham_** | **N_Theta_** | **DF1** | **DF2** | **F-value** | ***p*-value** |
| Treatment (Sham, Theta) | 7 | 6 | 1 | 11 | 14.60078847 | 0.00284 |
| ROI (Ipsilateral, Contralateral) | 7 | 6 | 1 | 11 | 1.013753854 | 0.336 |
| Treatment x ROI | 7 | 6 | 1 | 11 | 0.062016076 | 0.808 |
| vGCL DCX+ cell density | **N_Sham_** | **N_Theta_** | **DF1** | **DF2** | **F-value** | ***p*-value** |
| Treatment (Sham, Theta) | 7 | 6 | 1 | 11 | 0.144739048 | 0.711 |
| ROI (Ipsilateral, Contralateral) | 7 | 6 | 1 | 11 | 0.008602732 | 0.928 |
| Treatment x ROI | 7 | 6 | 1 | 11 | 0.345651821 | 0.568 |
| Proliferative stage | **N_Sham_** | **N_Theta_** | **DF1** | **DF2** | **F-value** | ***p*-value** |
| Treatment (Sham, Theta) | 7 | 6 | 1 | 11 | 6.595775576 | 0.0261 |
| ROI (Ipsilateral, Contralateral) | 7 | 6 | 1 | 11 | 2.058519733 | 0.179 |
| Treatment x ROI | 7 | 6 | 1 | 11 | 0.05343599 | 0.821 |
| Intermediate stage | **N_Sham_** | **N_Theta_** | **DF1** | **DF2** | **F-value** | ***p*-value** |
| Treatment (Sham, Theta) | 7 | 6 | 1 | 11 | 22.3347698 | 0.000623 |
| ROI (Ipsilateral, Contralateral) | 7 | 6 | 1 | 11 | 2.48574047 | 0.143 |
| Treatment x ROI | 7 | 6 | 1 | 11 | 2.82896046 | 0.121 |
| Postmitotic stage | **N_Sham_** | **N_Theta_** | **DF1** | **DF2** | **F-value** | ***p*-value** |
| Treatment (Sham, Theta) | 7 | 6 | 1 | 11 | 4.92456963 | 0.0484 |
| ROI (Ipsilateral, Contralateral) | 7 | 6 | 1 | 11 | 1.29865673 | 0.279 |
| Treatment x ROI | 7 | 6 | 1 | 11 | 0.82236279 | 0.384 |

**Table S6: Statistical results related to Figure 5.** The results were analysed with repeated measures ANOVA using the function 'aov_car’ from the ‘afex’ package in R. We We assessed the normality of the residuals with the Shapiro-Wilk test. We report deegrees pf freedom (DF), F statistic (F-value) and p-value for the model. All results obtained with the manual quantification of BrdU+ cells (Figure 5F) were analysed with two-sided unpaired two-samples Wilcoxon rank-sum tests in R due to the presence of outliers.

| Repeated measures ANOVA statistical results associated to DCX+ cell density | | | | | | | | | |
| --- | --- | --- | --- | --- | --- | --- | --- | --- | --- |
| GCL DCX+ cell density | | **N_Sham_** | **N_Theta_** | **DF1** | **DF2** | | **F-value** | | ***p*-value** |
| Outliers included | Treatment (Sham, Theta) | 8 | 8 | 1 | 14 | | 1.44548604 | | 0.249 |
|  | ROI (Ipsilateral, Contralateral) | 8 | 8 | 1 | 14 | | 1.32267637 | | 0.269 |
|  | Treatment x ROI | 8 | 8 | 1 | 14 | | 0.51366871 | | 0.485 |
| Outliers removed | Treatment (Sham, Theta) | 8 | 7 | 1 | 13 | | 0.38609369 | | 0.545 |
|  | ROI (Ipsilateral, Contralateral) | 8 | 7 | 1 | 13 | | 3.74726458 | | 0.0749 |
|  | Treatment x ROI | 8 | 7 | 1 | 13 | | 0.02921728 | | 0.867 |
| dGCL DCX+ cell density | | **N_Sham_** | **N_Theta_** | **DF1** | **DF2** | | **F-value** | | ***p*-value** |
| Outliers included | Treatment (Sham, Theta) | 8 | 8 | 1 | 14 | | 2.96046942 | | 0.107 |
|  | ROI (Ipsilateral, Contralateral) | 8 | 8 | 1 | 14 | | 5.07206819 | | 0.0409 |
|  | Treatment x ROI | 8 | 8 | 1 | 14 | | 0.63644349 | | 0.438 |
| Outliers removed | Treatment (Sham, Theta) | 8 | 7 | 1 | 13 | | 1.93752809 | | 0.187 |
|  | ROI (Ipsilateral, Contralateral) | 8 | 7 | 1 | 13 | | 9.96411609 | | 0.00757 |
|  | Treatment x ROI | 8 | 7 | 1 | 13 | | 2.32129142 | | 0.152 |
| vGCL DCX+ cell density | | **N_Sham_** | **N_Theta_** | **DF1** | **DF2** | | **F-value** | | ***p*-value** |
| Outliers included | Treatment (Sham, Theta) | 8 | 8 | 1 | 14 | | 0.07377716 | | 0.79 |
|  | ROI (Ipsilateral, Contralateral) | 8 | 8 | 1 | 14 | | 2.35762838 | | 0.147 |
|  | Treatment x ROI | 8 | 8 | 1 | 14 | | 7.11488635 | | 0.0184 |
| Outliers removed | Treatment (Sham, Theta) | 8 | 7 | 1 | 13 | | 0.9169635 | | 0.356 |
|  | ROI (Ipsilateral, Contralateral) | 8 | 7 | 1 | 13 | | 1.693705 | | 0.216 |
|  | Treatment x ROI | 8 | 7 | 1 | 13 | | 5.6071532 | | 0.0341 |
| Proliferative stage | | **N_Sham_** | **N_Theta_** | **DF1** | **DF2** | | **F-value** | | ***p*-value** |
| Treatment (Sham, Theta) | | 8 | 8 | 1 | 14 | | 2.27363494 | | 0.154 |
| ROI (Ipsilateral, Contralateral) | | 8 | 8 | 1 | 14 | | 6.34741509 | | 0.0245 |
| Treatment x ROI | | 8 | 8 | 1 | 14 | | 0.85037211 | | 0.372 |
| Intermediate stage | | **N_Sham_** | **N_Theta_** | **DF1** | **DF2** | | **F-value** | | ***p*-value** |
| Treatment (Sham, Theta) | | 8 | 8 | 1 | 14 | | 4.92635508 | | 0.0435 |
| ROI (Ipsilateral, Contralateral) | | 8 | 8 | 1 | 14 | | 2.16895423 | | 0.163 |
| Treatment x ROI | | 8 | 8 | 1 | 14 | | 0.16718734 | | 0.689 |
| Postmitotic stage | | **N_Sham_** | **N_Theta_** | **DF1** | **DF2** | | **F-value** | | ***p*-value** |
| Outliers included | Treatment (Sham, Theta) | 8 | 8 | 1 | 14 | | 1.31333457 | | 0.271 |
|  | ROI (Ipsilateral, Contralateral) | 8 | 8 | 1 | 14 | | 0.49762384 | | 0.492 |
|  | Treatment x ROI | 8 | 8 | 1 | 14 | | 0.04016785 | | 0.844 |
| Outliers removed | Treatment (Sham, Theta) | 8 | 7 | 1 | 13 | | 0.287475613 | | 0.601 |
|  | ROI (Ipsilateral, Contralateral) | 8 | 7 | 1 | 13 | | 0.115460676 | | 0.739 |
|  | Treatment x ROI | 8 | 7 | 1 | 13 | | 0.027334001 | | 0.871 |
| Post-hoc statistical comparisons associated to DCX+ cell density | | | | | | | | | |
| vGCL DCX+ cell density | | **N_Sham_** | **N_Theta_** | **Estimate** | | **SEM** | **t** | ***p*-value** | |
| Outliers  included | Theta (contra) vs  Theta (ipsi) | 8 | 8 | -45.1251 4837 | | 15.18419118 | -2.97185 0646 | 0.0606 | |
|  | Theta (contra) vs  Sham (contra) | 8 | 8 | -24.0020 5195 | | 16.83162228 | -1.42600 9422 | 0.852 | |
|  | Theta (contra) vs  Sham (ipsi) | 8 | 8 | -11.8488 3108 | | 20.16776095 | -0.58751 3463 | 0.938 | |
|  | Theta (ipsi) vs  Sham (contra) | 8 | 8 | 21.12309641 | | 20.16776095 | 1.047369436 | 0.938 | |
|  | Theta (ipsi) vs Sham (ipsi) | 8 | 8 | 33.27631728 | | 23.02550011 | 1.445194116 | 0.852 | |
|  | Sham (contra) vs Sham (ipsi) | 8 | 8 | 12.15322087 | | 15.18419118 | 0.80038645 | 0.938 | |
| Outliers  removed | Theta (contra) vs  Theta (ipsi) | 8 | 7 | -41.8132 681 | | 16.6437934 | -2.51224 387 | 0.13 | |
|  | Theta (contra) vs  Sham (contra) | 8 | 7 | -36.6187 535 | | 11.9673764 | -3.05988 149 | 0.0547 | |
|  | Theta (contra) vs  Sham (ipsi) | 8 | 7 | -24.4655 326 | | 15.0080911 | -1.63015 619 | 0.508 | |
|  | Theta (ipsi) vs  Sham (contra) | 8 | 7 | 5.19451467 | | 15.393525 | 0.33744803 | 1 | |
|  | Theta (ipsi) vs Sham (ipsi) | 8 | 7 | 17.3477355 | | 17.86016 | 0.97130908 | 1 | |
|  | Sham (contra) vs Sham (ipsi) | 8 | 7 | 12.1532209 | | 15.5688432 | 0.78061168 | 1 | |
| Wilcoxon rank sum test comparisons associated to BrdU+ cell density | | | | | | | | | |
| Variable | | **N_Sham_** | **N_Theta_** | **U** | | | ***p*-value** | | |
| BrdU+ NeuN+ DCX- cell density in the dGCL | | 8 | 8 | 27 | | | 0.637 | | |
| BrdU+ NeuN- DCX- cell density in the dGCL | | 8 | 8 | 34 | | | 0.875 | | |
| Total BrdU+ cell density in hilus | | 8 | 8 | 38 | | | 0.564 | | |
| Repeated measures ANOVA statistical results associated to the OPS behavioural test | | | | | | | | | |
| Position 1 | | **N_Sham_** | **N_Theta_** | **DF1** | **DF2** | | **F-value** | | ***p*-value** |
| Treatment (Sham, Theta) | | 8 | 7 | 1 | 13 | | 0.0090144 | | 0.926 |
| Session (Training, Test) | | 8 | 7 | 1 | 13 | | 8.28622166501848 | | 0.0129 |
| Treatment x Session | | 8 | 7 | 1 | 13 | | 0.0362233712267633 | | 0.852 |
| Position 2 | | **N_Sham_** | **N_Theta_** | **DF1** | **DF2** | | **F-value** | | ***p*-value** |
| Outliers included | Treatment (Sham, Theta) | 8 | 7 | 1 | 13 | | 1.05756746253476 | | 0.323 |
|  | Session (Training, Test) | 8 | 7 | 1 | 13 | | 2.19474142999942 | | 0.162 |
|  | Treatment x Session | 8 | 7 | 1 | 13 | | 0.314900397290363 | | 0.584 |
| Outliers removed | Treatment (Sham, Theta) | 7 | 7 | 1 | 12 | | 0.527595007593732 | | 0.482 |
|  | Session (Training, Test) | 7 | 7 | 1 | 12 | | 1.24821626868522 | | 0.286 |
|  | Treatment x Session | 7 | 7 | 1 | 12 | | 0.054111077989305 | | 0.82 |
| Position 3 | | **N_Sham_** | **N_Theta_** | **DF1** | **DF2** | | **F-value** | | ***p*-value** |
| Treatment (Sham, Theta) | | 8 | 7 | 1 | 13 | | 2.85455288782472 | | 0.115 |
| Session (Training, Test) | | 8 | 7 | 1 | 13 | | 25.1663935593637 | | 0.000236 |
| Treatment x Session | | 8 | 7 | 1 | 13 | | 1.30444962308038 | | 0.274 |

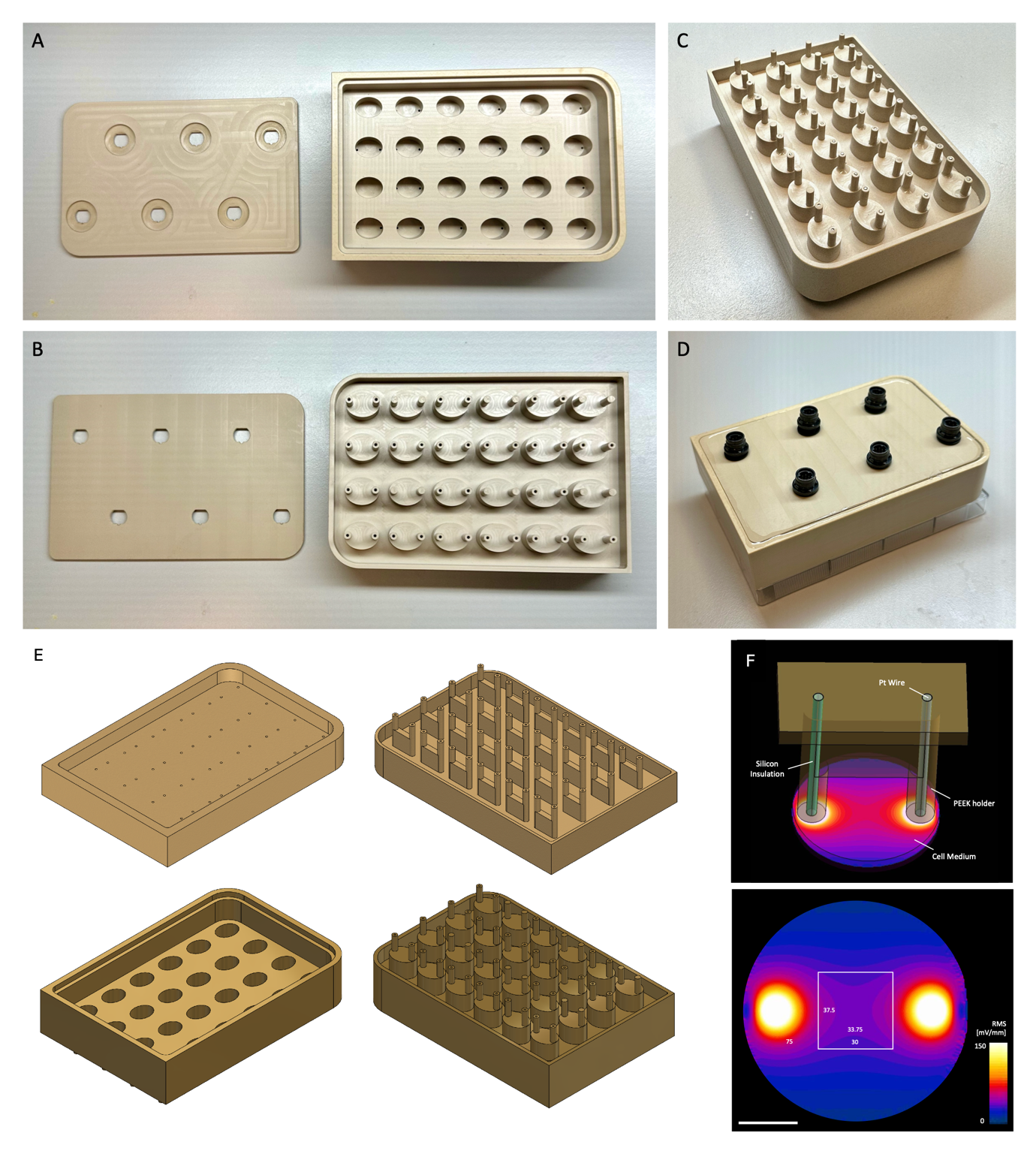

**Figure S5: Representative images of the in vitro TI stimulation device, related to Figure 1, Figure 3, and methods section.**

(A) PEEK components respectively showing the inner facing side and the outer facing side;
(B) Assembled device from the well plate facing side.
(C) Final setup placed on a well plate.
(D) 3D models produced in Solidworks of the first iteration of the device.
(E) Electric field distribution FEM modelling. The spatial distribution of the electric field and current density across the culture medium were modelled to understand the variability of the electrical cue delivered to the cells in the culture. Above: Schematic depicting the components of a single stimulation unit; below: current density distribution in A/m^2^ RMS at the bottom of the well (1 V at 1 kHz applied between Pt electrodes, the square indicates the area considered for quantification in the IHC quantifications in **Figure 1**).

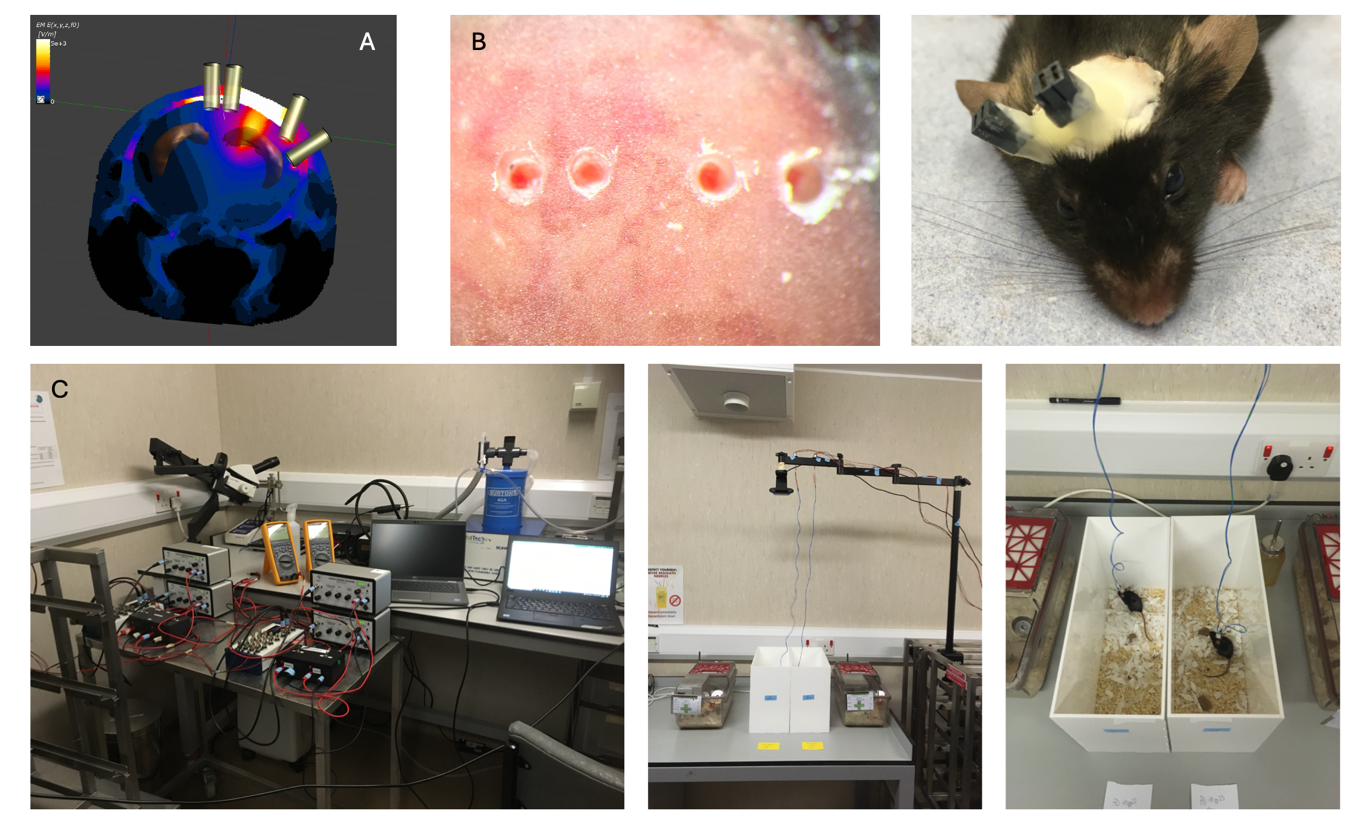
**Figure S6: TI stimulation setup, related to methods section, Figure 3, 4 and 5.**

(A) Finite element modelling of the TI stimulation targeting the CA1 of the right hippocampus in adult mice, developed with Sim4Life as described in the methods section, replicating a procedure from from Missey^5^ and Acerbo et al^6^ (coordinates: AP -2, ML +0.7, -0.7, -3.1 and -4.5 mm).
(B) Representative images of the TI stimulation electrode implantation as described in the methods section. Briefly, four 0.7 mm holes in the marked locations were drilled through the cement and the skull down to the dura mater. Four stainless steel screws (TX000-1.5FH, 0.86 mm diameter, Component Supply), previously attached in pairs to micro-connector sockets, were screwed into the holes.
(C) TI stimulation setup optimised for freely-moving animals.

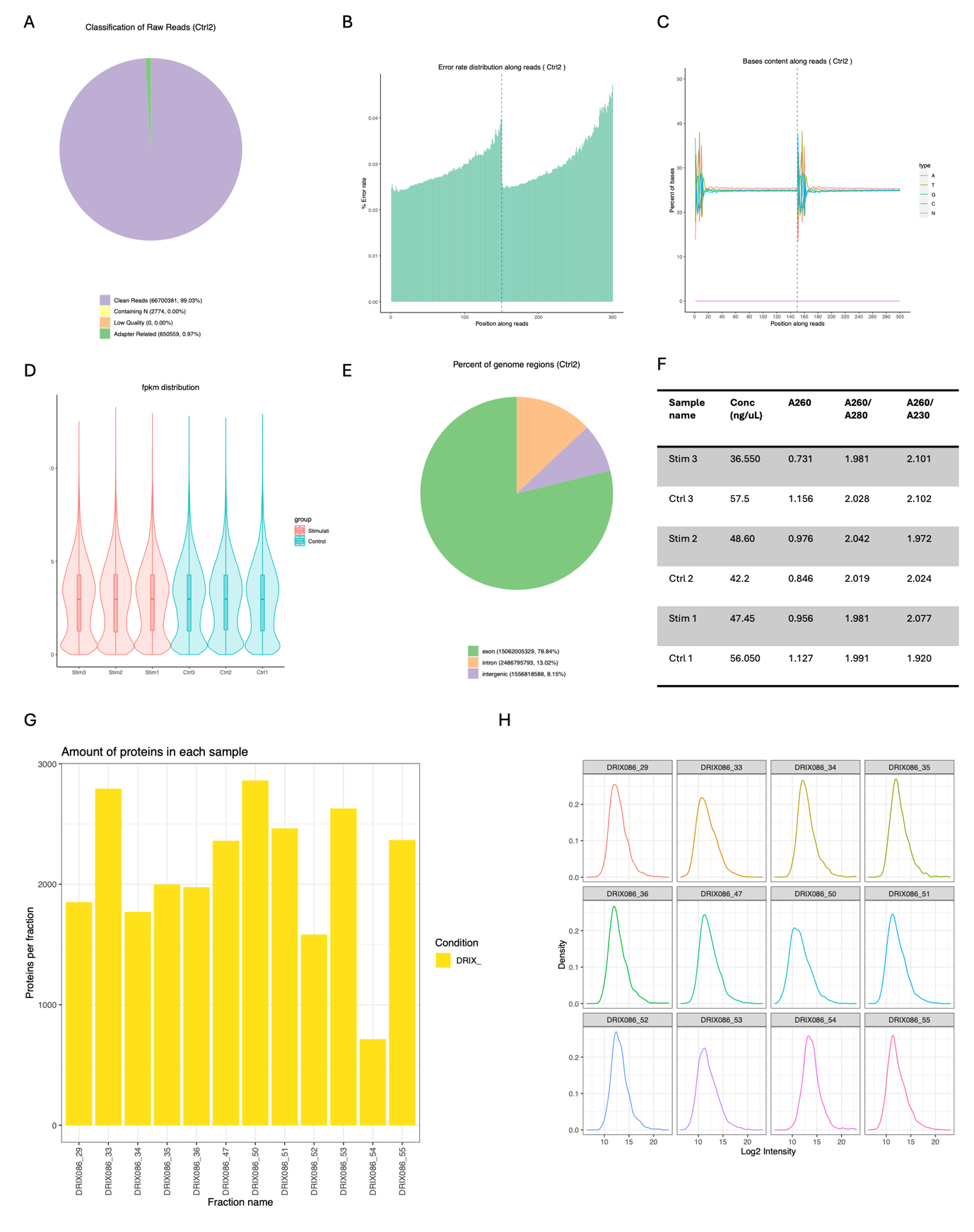
**Figure S7: Quality control measures for the mRNA sequencing analysis related to Figure 1, and the proteomics analysis related to Figure 3.**

(A) Representative classification of the raw reads for Ctrl2: reads were removed when uncertain nucleotides were more than 10 per cent of either read (N >10%), when low-quality nucleotides (Base Quality less than 5) constituted more than 50 per cent of the read, when reads were contaminated by an adapter;
(B) Representative error rate distribution along reads of semple Ctrl2;
(C) Representative Bases content along reads for Ctrl2;
(D) Representative FKPM distribution across controls and stimulation conditions;
(E) Representative percentages of mapped genome regions in Ctrl2 (exons, introns and intergenic).
(F) RNA purity values from the mRNA sequencing samples. The ratio A260/A280 is a metric for RNA and DNA content, and the A260/A230 was used for quantifying purity measures purity over salts and contaminants at neutral pH. Only samples with a purity ratio above 1.9 were included in the dataset.
(G) Amount of protein in each sample of ipsilateral (right) dentate gyrus (DG) tissue, analysed with mass spectrometry-based proteomics, related to Figure 3 and proteomics method section.
(H) Protein density spectra of all samples included in the proteomics analysis (Figure 3).

**Supplemental References**

1.	Keselman, H. J., Kowalchuk, R. K., Algina, J. & Wolfinger, R. D. The analysis of repeated measurements: A comparison of mixed-model Satterthwaite F tests and a nonpooled adjusted degrees of freedom multivariate test. *Commun Stat Theory Methods* **28**, 2967–2999 (1999).

2.	Morrell, C. H. Likelihood Ratio Testing of Variance Components in the Linear Mixed-Effects Model Using Restricted Maximum Likelihood. *Biometrics* **54**, 1560 (1998).

3.	Sture Holm. A Simple Sequentially Rejective Multiple Test Procedure. *Scandinavian Journal of Statistics* **6**, 65–70 (1979).

4.	MacFarland, T. W. & Yates, J. M. Mann–Whitney U Test. *Introduction to Nonparametric Statistics for the Biological Sciences Using R* 103–132 (2016) doi:10.1007/978-3-319-30634-6_4.

5.	Missey, F. *et al.* Orientation of Temporal Interference for Non-invasive Deep Brain Stimulation in Epilepsy. *Front Neurosci* **15**, 633988 (2021).

6.	Acerbo, E. *et al.* Focal non-invasive deep-brain stimulation with temporal interference for the suppression of epileptic biomarkers. *Front Neurosci* **16**, 945221 (2022).
